# Novel regulators of islet function identified from genetic variation in mouse islet Ca^2+^ oscillations

**DOI:** 10.1101/2022.11.26.517741

**Authors:** Christopher H. Emfinger, Lauren E. Clark, Brian Yandell, Kathryn L. Schueler, Shane P. Simonett, Donnie S. Stapleton, Kelly A. Mitok, Matthew J. Merrins, Mark P. Keller, Alan D. Attie

## Abstract

Insufficient insulin secretion to meet metabolic demand results in diabetes. The intracellular flux of Ca^2+^ into β-cells triggers insulin release. Since genetics strongly influences variation in islet secretory responses, we surveyed islet Ca^2+^ dynamics in eight genetically diverse mouse strains. We found high strain variation in response to four conditions: 1) 8 mM glucose; 2) 8 mM glucose plus amino acids; 3) 8 mM glucose, amino acids, plus 10 nM GIP; and 4) 2 mM glucose. These stimuli interrogate β-cell function, α-cell to β-cell signaling, and incretin responses. We then correlated components of the Ca^2+^ waveforms to islet protein abundances in the same strains used for the Ca^2+^ measurements. To focus on proteins relevant to human islet function, we identified human orthologues of correlated mouse proteins that are proximal to glycemic-associated SNPs in human GWAS. Several orthologues have previously been shown to regulate insulin secretion (e.g. ABCC8, PCSK1, and GCK), supporting our mouse-to-human integration as a discovery platform. By integrating these data, we nominated novel regulators of islet Ca^2+^ oscillations and insulin secretion with potential relevance for human islet function. We also provide a resource for identifying appropriate mouse strains in which to study these regulators.

## INTRODUCTION

The majority of gene loci responsible for the genetic variation in type 2 diabetes (T2D) susceptibility affect the function of endocrine cells of pancreatic islets, primarily β-cells (1, 2). Variation in β-cell mass and function places boundaries on their capacity to respond to acute and chronic demands for insulin, such as those of overnutrition and insulin resistance (1, 2). Therefore, metabolic challenges are useful in genetic screens because they expose phenotypes that would otherwise remain silent.

The large collection of inbred mouse strains provides us with a wide repertoire of genetic and phenotype diversity, comparable to that of the entire human population (3). Yet, most mouse studies have been confined to a small number of highly inbred strains (3, 4). It is becoming widely appreciated that gene deletions, nutritional interventions, and drug effects vary widely among mouse strains, as they do in humans (3, 5). Thus, characterization of the basis for this high level of phenotype variation is a path to gain deeper insights into the pathophysiology and genetics of a wide range of physiological processes.

The pancreatic β-cell is a nutrient sensor. In response to particular nutrient stimuli (e.g. glucose, amino acids), the β-cells generate ATP and close ATP-dependent K^+^ channels (K_ATP_), resulting in plasma membrane depolarization (6–8). This leads to an oscillatory influx of Ca^2+^ ions, triggering insulin secretion. The process of secreting insulin and re-compartmentalizing Ca^2+^ ions consumes ATP, and the drop in the ATP/ADP ratio reopens K_ATP_ channels, repolarizing the membrane, and closing membrane Ca^2+^ channels. Consequently, oscillations in metabolism, insulin secretion, and Ca^2+^ are intrinsically linked (8–12), and the capacity to maintain functional Ca^2+^ handling has been suggested to be critical for islet compensation (13).

In this study, we utilized the extraordinary genetic and phenotypic diversity represented in the eight founder mouse strains (which we subsequently refer to as “founders”) used to generate the Collaborative Cross (CC) recombinant inbred mouse panel and the Diversity Outbred (DO) stock (14, 15). These strains capture most of the genetic diversity of all inbred mouse strains (14, 15). While studies of these mice have provided significant insight into genetic regulators of islet function (16), determining the appropriate model system for evaluating genes of interest is often difficult as most deletion models are made in only a small number of strains, primarily the C57BL/6J or C57BL/6N.

We explored the diversity of nutrient-evoked islet Ca^2+^ responses across the eight founder mouse strains, uncovering a remarkable diversity of Ca^2+^ oscillations. Our prior proteomics studies showed that the protein abundance from islets of the founder mouse strains is also highly diverse, as is their insulin secretory response to different stimuli (17). By correlating the strain and sex variation in protein abundance with the variation in Ca^2+^ oscillations, we identified a small number of islet proteins that are highly correlated with islet Ca^2+^ oscillations. The human orthologues of many of these proteins are encoded by genes with nearby SNPs linked to glycemic traits (e.g., fasting blood glucose, see **Table 2** for terms) in genome-wide association studies (GWAS). By integrating these data, we nominated novel regulators of islet Ca^2+^ oscillations and insulin secretion with potential relevance for human islet function. We provide a web-based resource that integrates proteomic and Ca^2+^ data for identifying appropriate mouse strains in which to study these regulators.

**Table 1:**
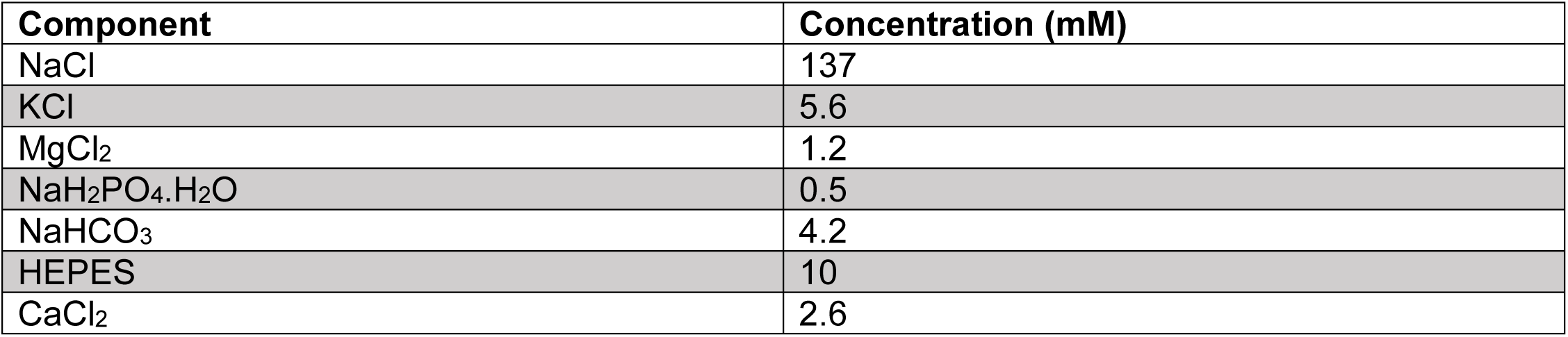
Imaging medium formula. Components are indicated by chemical abbreviation on the left and final concentration in mM is indicated in the right column.

**Table 2:** Categories included in SNP queries. These terms were considered as glycemia-related and are categorized as such on the Common Metabolic Diseases Knowledge portal, which was queried for the relevant SNPs. Also included but not listed here were variations of these terms that were adjusted for BMI.

| Fasting hormones | Glucose-related | Tolerance test | Diabetes risk |
| --- | --- | --- | --- |
| insulin | fasting glucose | 2hr glucose | T1D |
| proinsulin | random glucose | 2hr insulin | T2D |
| c-peptide | Hba1c | 2hr c-peptide |  |
| fasting glc-BMI interaction |  | 2hr insulin |  |
| fasting ins-BMI interaction |  | Acute insulin response |  |
| Gestational diabetes/altered fast glucose in pregnancy |  | SI-adjusted acute ins. Resp. |  |
|  |  | AUC insulin |  |
|  |  | AUC insulin/AUC glucose |  |
|  |  | Corrected insulin response |  |
|  |  | HOMA-B |  |
|  |  | HOMA-IR |  |
|  |  | Ins. Secretion rate |  |
|  |  | Ins. Sensitivity |  |
|  |  | Incremental ins. @ 30 min OGTT |  |
|  |  | Insulin @ 30min OGTT |  |
|  |  | Peak ins. response |  |
|  |  | Peak ins. Response adj SI |  |

## RESULTS

### Genetics exerts a strong influence on islet Ca^2+^ dynamics

Glucose metabolism, β-cell Ca^2+^ flux, and insulin secretion are pulsatile, and have been found to oscillate in both humans and mice (8, 9, 18–20). Because they are interconnected, understanding the factors governing oscillation patterns can inform about the mechanisms that regulate insulin secretion (6, 11, 21, 22). To explore the influence of genetic background on Ca^2+^ oscillations, we measured Ca^2+^ in islets of the eight CC founder strains, that together harbor as much genetic diversity as humans: A/J, C57BL/6J (B6), 129S1/SvlmJ (129), NOD/ShiLtJ (NOD), NZO/HILtJ (NZO), CAST/EiJ (CAST), PWK/PhJ (PWK), and WSB/EiJ (WSB).

All mice were maintained on a Western-style diet (WD) high in fat and sucrose for 16 weeks, prior to isolating their islets for Ca^2+^ imaging with Fura Red, a Ca^2+^-sensitive fluorescent dye (**Figure 1A**). We measured Ca^2+^ dynamics in response to four conditions: 1) 8 mM glucose (8G); 2) 8 mM glucose + 2 mM glutamine, 0.5 mM leucine, and 1.5 mM alanine (8G/QLA); 3) 8G/QLA + 10 nM glucose-dependent insulinotropic polypeptide (8G/QLA/GIP); and 4) 2 mM glucose (2G) (**Figure 1B**). There was a high degree of similarity between three of the five classical strains (A/J, B6, 129), which were dominated by slow oscillations (period 2-10 minutes) in 8G and 8G/QLA/GIP, and had relatively fewer islets reach plateau (continuous peak activity without oscillation) in 8G/QLA. Likewise, the wild-derived strains (CAST, WSB, and PWK) closely matched one another, while differing from the classic strains. The wild-derived mouse islets were dominated by fast oscillations (period <2 minutes) in 8G, resulting in plateaus for 8G/QLA and 8G/QLA/GIP.

**Figure 1.**
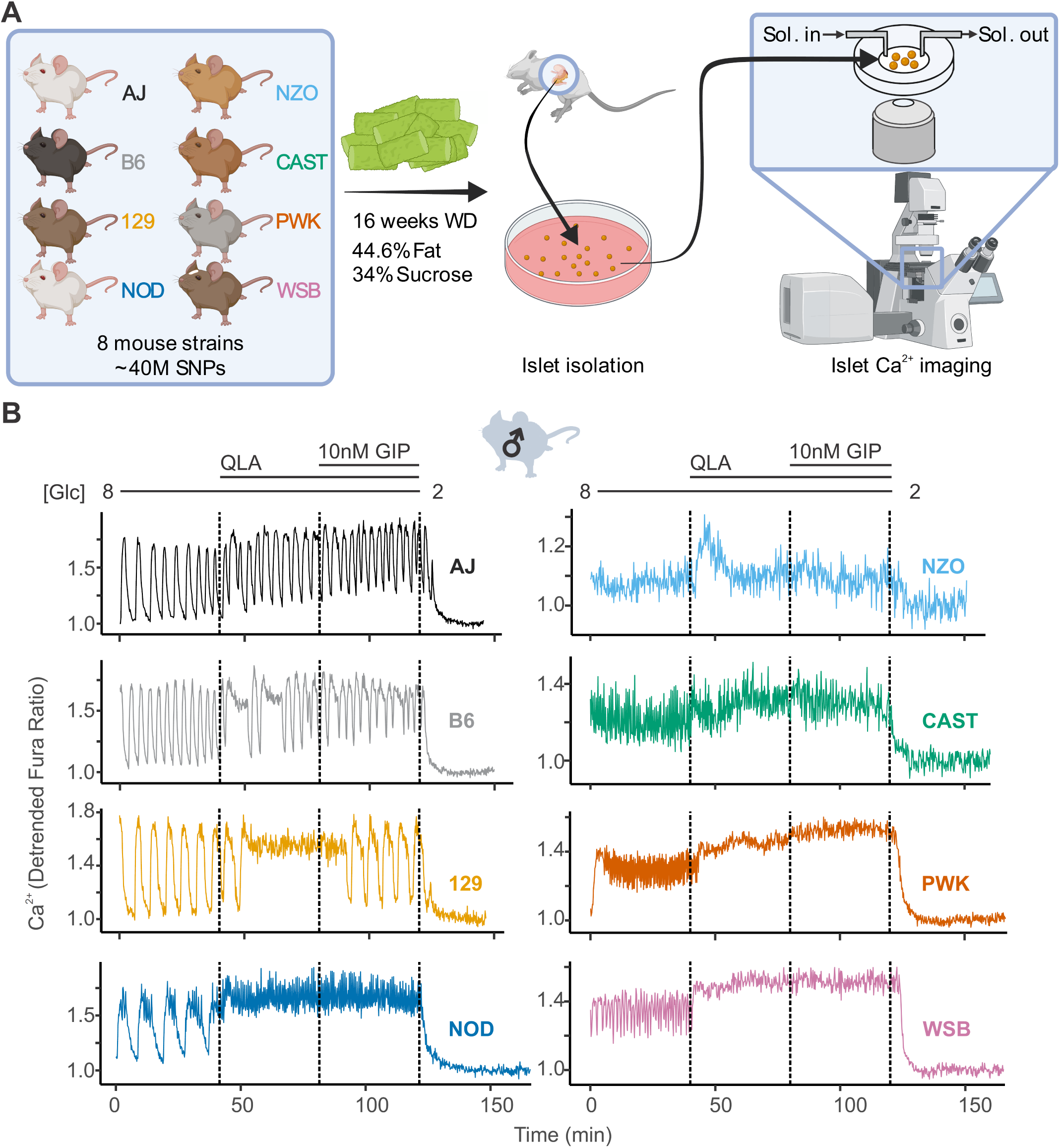
High diversity in Ca^2+^ oscillation across eight genetically distinct mouse strains. **(A)** Male and female mice from eight strains (A/J; C57BL/6J (B6); 129S1/SvImJ (129); NOD/ShiLtJ (NOD); NZO/HILtJ (NZO); CAST/EiJ (CAST); PWK/PhJ (PWK); and WSB/EiJ (WSB)) were placed on a Western Diet (WD) for 16 weeks, before their islets were isolated. The islets were then imaged on a confocal microscope using Fura Red dye under conditions of 8 mM glucose; 8 mM glucose + 2 mM L-glutamine, 0.5 mM L-leucine, and 1.25 mM L-alanine (QLA); 8 mM glucose + QLA + 10 nM GIP; and 2 mM glucose. **(B)** Representative Ca^2+^ traces for male mice (n = 3-8 mice per strain, and 15-83 islets per mouse), with the transitions between solution conditions indicated by dashed lines. Abbreviations: ‘[Glu]’ = ‘concentration of glucose in mM’ ; ‘Sol.’ = ‘solution’; ‘SNPs’= ‘single-nucleotide polymorphisms’

Two strains stood out from the others. Islets from NOD mice showed characteristics from both the wild-derived and classic strains; slow oscillations in 8G and a sustained plateau in response to 8G/AA and 8G/QLA/GIP with fast oscillations superimposed. The NZO mice also differed from the other classic strains, likely because they were all diabetic (blood glucose >250 mg/dL). Their islets were minimally responsive to 8G but did respond with a strong pulse in 8G/QLA and Ca^2+^ remained elevated in 8G/QLA/GIP.

Many of the strain differences seen in the male mice were maintained in the females (**Supplemental Figure 1A**). The classic strains were once again highly similar to one another, as were the wild-derived strains. Furthermore, the NZO females, of which all but one were diabetic, mirrored the behavior of the male islets. One interesting observation that emerged from the female islets is that the NOD females displayed a greater variation in their Ca^2+^ oscillations than the NOD males (**Supplemental Figure 2**). Some of the islets maintained slow oscillations throughout the various conditions, while some demonstrated fast oscillations and plateaued like the wild-derived strains. Yet others appeared strikingly similar to the islets from diabetic NZO mice, despite none of the NOD mice being diabetic. Finally, the one non-diabetic female NZO displayed oscillatory behavior comparable to that of the other classic strains, with clear, slow oscillations (**Supplemental Figure 1B**).

### Dissecting islet Ca^2+^ dynamics

An understanding of the mechanisms regulating insulin secretion, including the roles of specific metabolic pathways, ion channels, and hormones, has been derived from the shape and frequency of islet Ca^2+^ oscillations (6, 9, 11, 19, 23–26). To elucidate strain differences in Ca^2+^ dynamics, we focused on six parameters of the Ca^2+^ waveform (**Figure 2A**): 1) peak Ca^2+^ (the maximum value of each oscillation); 2) period (the length of time between two peaks); 3) active duration (the length of time for each Ca^2+^ oscillation measured at half of the peak height, also known the oxidative “secretory” phase, or “Mito_Ox_” (8); 4) pulse duration (active duration plus extra time for Ca^2+^ extrusion); 5) silent duration (the electrically-silent “triggering” phase, also known as “Mito_Cat_” (8), which culminates in K_ATP_ closure and membrane depolarization); and 6) plateau fraction (the active duration divided by the period, or the fraction of time spent in the active “secretory” phase).

**Figure 2.**
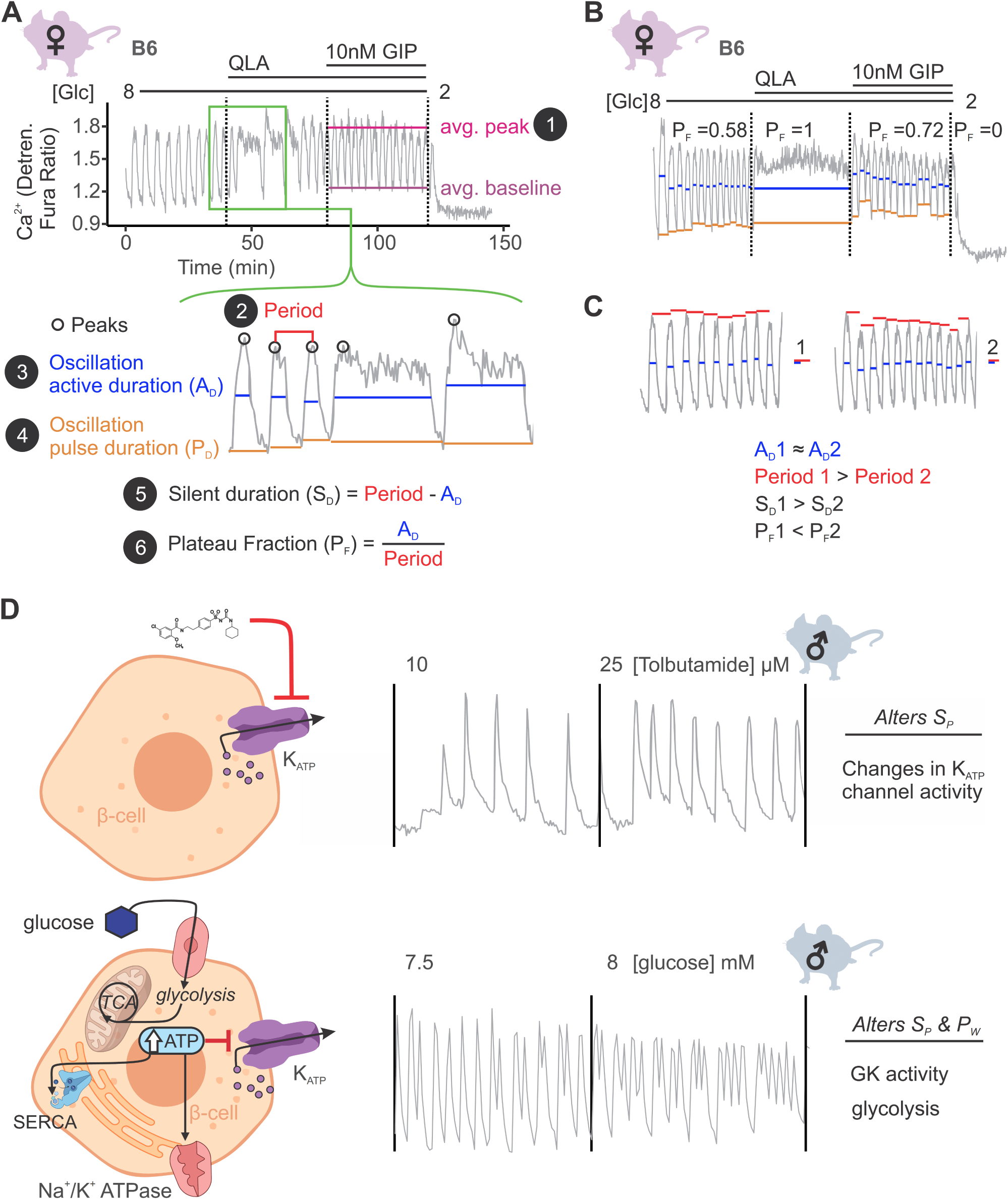
Ca^2+^ wave breakdown reveals mechanisms underlying Ca^2+^ responses. **(A)** In the example B6 female Ca^2+^ wave, the islet oscillations can change in their average peak (1) and average baseline in response to different nutrients. Additionally, shifts in wave shape (green box) can be broken down into changes in time between peaks (period, **2**), the time in the active phase (active duration, A_D_, **3**), and the length of the oscillation (pulse duration, P_D_, **4**). From these, the time inactive between oscillations (silent duration, S_D_, **5**), and the relative time in the active phase, or plateau-fraction (P_F_, **6**), can be calculated. Each parameter can be changed by different underlying mechanisms. **(B)** For islets that plateaued, as in the example islet in 8/QLA, they were assigned a plateau-fraction of one and a period of zero. For islets that ceased to oscillate, such as the example islet in 2 mM glucose, they were assigned a plateau-fraction of zero and a period of the time of measurement (40 minutes). **(C)** For trace 1 (left), which has a longer period (red bars) than trace 2 (right), but the same active duration (blue bars), the silent duration is greater and consequently the P_F_ is shorter, in contrast to the trace in (**A**) where the P_F_ increases between 8mM and 8/QLA are largely due to increases in A_D_. **(D)** Changes in specific Ca^2+^ wave parameters can reflect different mechanisms in β-cells. For example, changing K_ATP_ activity pharmacologically (upper panels) predominantly increases P_F_ by altering S_D_, whereas increasing glucose concentrations by elevating glucose or activating GK cause significant alterations in both A_D_ and S_D_ to increase P_F_. Abbreviations: ‘[Glc]’ = ‘concentration of glucose in mM’ ; ‘GK’ = ‘glucokinase’

We also assessed the spectral density for every islet to extract additional information from complex oscillations where multiple components were visible (**Supplemental Figure 3A**). We analyzed each trace to determine the top two frequencies contributing to the trace (1^st^ and 2^nd^ component frequencies, **Supplemental Figure 3B**) and their respective contributions (1^st^ and 2^nd^ component amplitudes). Because certain features, such as metabolically-driven (slow) and electrically-driven (fast) oscillations have characteristic frequencies (27), extracting the top two frequencies may highlight additional information beyond that previously collected.

A representative Ca^2+^ dynamic from a female B6 islet is illustrated in **Figure 2A**. The transition from 8G to 8G/QLA resulted in an increased active duration, yielding a longer period and increased plateau fraction. For an islet that plateaued at the peak, as seen in 8G/QLA (**Figure 2B**), we computed a plateau-fraction of one, an active and pulse duration of 40 minutes (the measurement time), and a period of zero minutes. An islet that returned to baseline and ceased to oscillate, as seen in 2 mM glucose (**Figure 2B**), was determined to have a plateau-fraction, active duration, and pulse duration of zero, and a period of 40 minutes. Dissecting the strain-dependence of these key parameters of the Ca^2+^ oscillations is important for identifying underlying mechanisms, as illustrated in **Figure 2C**. While both traces have a similar active duration (blue bars), trace 1 has a longer period (red bars), resulting in an increased silent duration and a decreased plateau-fraction.

Examples of pathways altering specific components of Ca^2+^ oscillations have previously been established (**Figure 2D**) (9, 11, 18, 28, 29). For example, when K_ATP_ channels are pharmacologically closed with tolbutamide, the silent duration is shortened, resulting in increased frequency without a change in pulse shape (upper panel). The addition of glucose leads to increased glucose metabolism and glucokinase (GK) activity (11). The resulting rise in ATP inhibits K_ATP_ channels (6) and is used as a substrate for additional processes that affect Ca^2+^, such as SERCA pumps (30, 31). Thus, glucose alters both the active and silent durations, resulting in a change in both frequency and shape of the Ca^2+^ oscillations (lower panel).

### Parameters significantly correlated with insulin secretion show remarkable variance by strain and sex

Average Ca^2+^ is commonly used for analyzing Ca^2+^ dynamics and is frequently assumed to be highly correlated to insulin secretion. To determine whether average Ca^2+^ is predictive of insulin secretion, we performed *ex vivo* perifusion studies on islets from WSB and 129 male mice, two strains that showed similar average Ca^2+^ (**Supplemental Figure 4**) but exhibited vastly different Ca^2+^ oscillations (**Figure 1B**). WSB mice had significantly higher insulin secretion in each of the secretory conditions (**Figure 3A**), suggesting another Ca^2+^ parameter better predicts insulin secretion.

**Figure 3.**
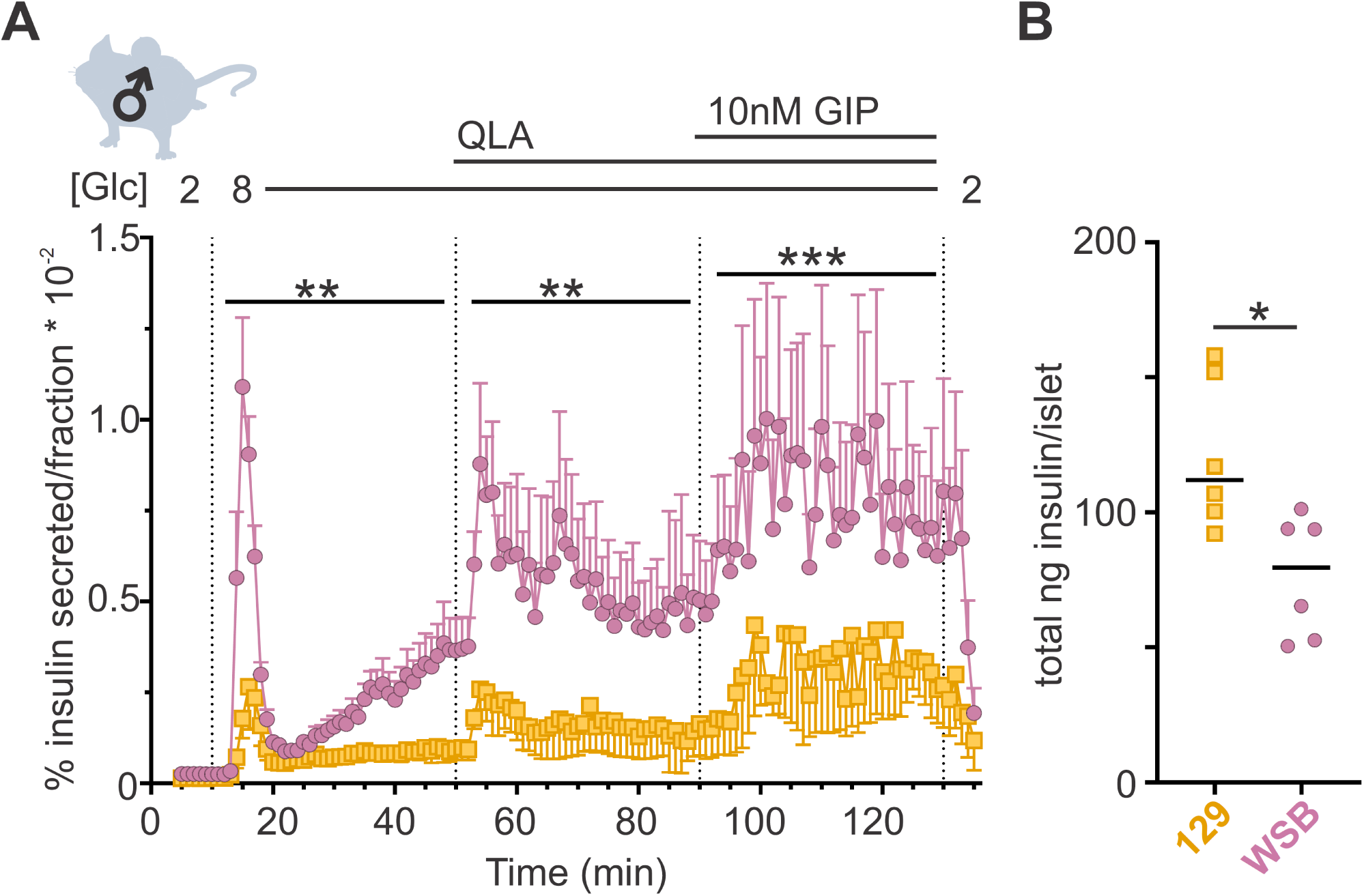
WSB mice secrete significantly more insulin than 129 mice. (**A**) Insulin secretion was measured for perifused islets from WSB (n=6, magenta circles) and 129 (n=5, yellow squares) male mice in 2 mM glucose, 8 mM glucose, 8 mM glucose + QLA, and 8 mM glucose + QLA + GIP. Transitions between solutions are indicated by dotted lines and the conditions for each are indicated above the graph. “[Glc]” denotes the concentration of glucose in mM. Data are shown as a percentage of total islet insulin (mean ± SEM). (**B**) Average total insulin per islet for the WSB and 129 males used in (**A**) with one exception: islets from one of the 129 mice were excluded from perifusion analysis due to technical issues with perifusion system on the day those animals’ islets were perifused. Dots represent individual values, and the mean is denoted by the black line. For (**A**), asterisks denote strain effect for the area-und-the-curve of the section determined by 2-way ANOVA, mixed effects model; ** p < 0.01, *** p < 0.001. For (**B**), asterisk denotes p < 0.05 from Student’s T-test with Welch’s correction.

To identify parameters of the Ca^2+^ dynamics most strongly correlated to insulin secretion, we computed the correlation between the Ca^2+^ oscillation parameters and our previously published insulin secretion in similar conditions (8.3G, 8.3G/QLA, basal) for the same sexes and strains (**Figure 4A**, **Supplemental Figure 5**, and **Supplemental Figure 6**) (17). Consistent with our observations from the perifusion data in the WSB and 129 islets, we found that average Ca^2+^ was not strongly correlated to insulin secretion. Other metrics, such as active duration in 8G, and the silent durations in 8G/QLA, were more highly correlated to insulin secretion. Meanwhile, the 1^st^ component frequency in 8G from the spectral density analysis was highly correlated with *decreased* insulin secretion. These metrics were also the most highly correlated with multiple clinical measures in the founder mice, particularly plasma insulin (**Figure 4B, Supplemental Figure 5**, and **Supplemental Figure 7**), for which silent duration in 8G/QLA/GIP had the strongest correlations.

**Figure 4.**
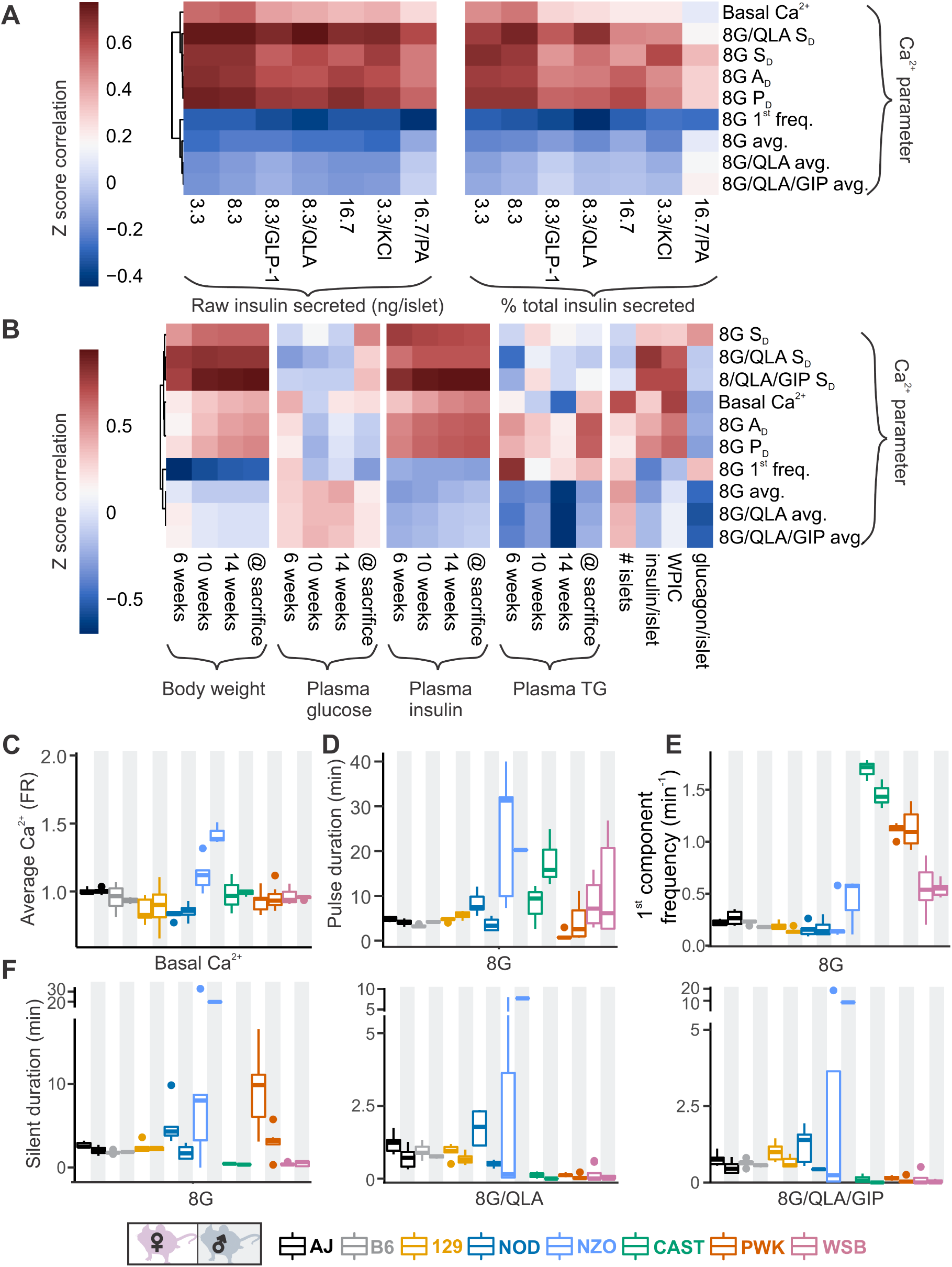
Comparing sex and strain patterns for Ca^2+^ metrics, insulin secretion, and clinical traits nominates Ca^2+^ metrics of interest. **(A)** The z-score correlation coefficient was calculated for Ca^2+^ parameters and raw insulin secreted and % total insulin secreted. Insulin measurements were previously collected for seven different secretagogues (16.7 mM glucose + 0.5 mM palmitic acid (16.7G/PA); 3.3 mM glucose + 50 mM KCl (3.3G/KCl); 16.7 mM glucose (16.7G); 8.3 mM glucose + 1.25 mM L-alanine, 2 mM L-glutamine, and 0.5 mM L-leucine (8.3G/QLA); 8.3 mM glucose + 100 nM GLP-1 (8.3G/GLP-1); 8.3 mM glucose (8.3G); and 3.3 mM glucose (3.3G)) (17). (**B**) Correlation of the Ca^2+^ parameters to the clinical measurements in the founder mice which include 1) plasma insulin, triglycerides, and glucose at 6, 10, and 14 weeks as well as at time of sacrifice; 2) number of islets; 3) whole-pancreas insulin content (WPIC); and 5) islet content for insulin and glucagon. For (**A**) and (**B**), the Ca^2+^ parameters shown here include average Ca^2+^ in 2 mM glucose (basal Ca^2+^); average Ca^2+^ in 8 mM glucose (8G avg.); average Ca^2+^ in 8 mM glucose + 1.25 mM L-alanine, 2 mM L-glutamine, and 0.5 mM L-leucine (8G/QLA avg); average Ca^2+^ in 8 mM glucose + QLA + 10 nM GIP (8G/QLA/GIP avg.); pulse duration in 8 mM glucose (8G P_D_); active duration in 8G (8G A_D_); silent duration in 8G (8G S_D_), 8G/QLA (8G/QLA/S_D_), and 8G/QLA/GIP (8G/QLA/GIP S_D_); and 1^st^ component frequency in 8 mM glucose (8G 1^st^ freq.). Other parameters analyzed are indicated in **Supplemental Figures 6 and 7.** (**B-E**) Sex and strain variability for **(C)** average Ca^2+^ determined by the Fura-ratio (FR) in 2 mM glucose, **(D)** pulse duration of oscillations in 8G, **(E)** 1^st^ component frequency in 8G and **(F)** silent duration of oscillations in 8G, 8G/QLA, and 8G/QLA/GIP.

Several parameters of the Ca^2+^ oscillatory waveform showed strong strain and sex effects (**Figure 4C-D** and **Supplemental Figure 5**). For example, basal Ca^2+^ (average Ca^2+^ in 2G, **Figure 4C**) was relatively consistent among the strains, except NZO where it was highest in islets from male mice. For the overall pulse duration (**Figure 4D**), the NZO mice were once again the highest, followed by CAST and WSB. A noticeable sex-effect was measured for the CAST mice, where male mice had a longer pulse duration than the female mice. The 1^st^ component frequency (**Figure 4E**) is driven by the differences observed in the wild-derived strains, for which CAST has the highest frequency, followed by PWK and WSB. Finally, the trend for a sex effect in the classic strains on the silent duration (at 8G, 8G/QLA, and 8G/QLA/GIP) is absent in the NZO and wild-derived mice with the former having greater silent duration in males and the latter frequently having islets plateau in response to these stimuli.

Clustering the Ca^2+^ responses into distinct groups based on our observations of the waveforms (**Figure 1B**, **Figure 4C-E**, and **Supplemental Figures 1 and 2**) also occurs when correlating individual Ca^2+^ parameters to ex vivo secretion and clinical data (**Supplemental Figure 5**). For example, the anticorrelation between the 1^st^ frequency component in 8G and percent insulin secreted in 8.3G/QLA (**Supplemental Figure 5A**) separates the classic inbred, wild-derived, and diabetes-susceptible strains into distinct groups despite the variability in the trait. Correlation between the silent duration in 8G/QLA to insulin secretion in 8.3G/QLA, likewise groups by strain (**Supplemental Figure 5B**). Finally, some correlations, such as that between 8G/QLA/GIP silent duration and plasma insulin at sacrifice (**Supplemental Figure 5C**), can be strongly influenced by outlier strains; e.g., NZO. Collectively, these data demonstrate that genetics has a profound influence on key parameters of islet Ca^2+^ oscillations.

### Calcium oscillatory parameters correlate strongly to the abundance of specific islet proteins

To explore relationships between Ca^2+^ oscillations and islet proteins, we took advantage of our whole islet proteomic survey from the eight founder strains (17). To identify proteins that may underly the strain-differences in Ca^2+^ oscillations, we computed the correlation between islet protein abundance and Ca^2+^ dynamics across all mice used in our study (**Figure 5** and **Supplemental Figure 8**). Our previous survey of islet proteomics included both sexes for all strains, except NZO males, resulting in a quantitative measure of 4054 proteins (17). **Figure 5A** illustrates a heatmap of the correlation between islet proteins and several parameters of Ca^2+^ oscillations. Unsupervised clustering was used to show that groups of proteins showed strong positive or negative correlation to a given Ca^2+^ parameter, yielding distinct correlation architecture. For example, proteins highly correlated to the 8G 1^st^ component frequency tended to also be strongly anticorrelated to the silent duration conditions, which were very similar to one another. The active and pulse durations for 8G had nearly identical correlation structure. Additionally, the conditions with the fewest highly correlated proteins were the average Ca^2+^ measures for 8G, 8G/QLA, and 8G/QLA/GIP, and the structure for these was largely inverted from the active duration conditions. Finally, despite the differences in the overall correlations between the different metrics, there were proteins that did overlap (e.g. the block of proteins with high correlation to both 8G A_D_ and 8G/QLA S_D_) suggesting that while there were clusters of distinct proteins/pathways for any given metric some proteins may modify more than one metric.

**Figure 5.**
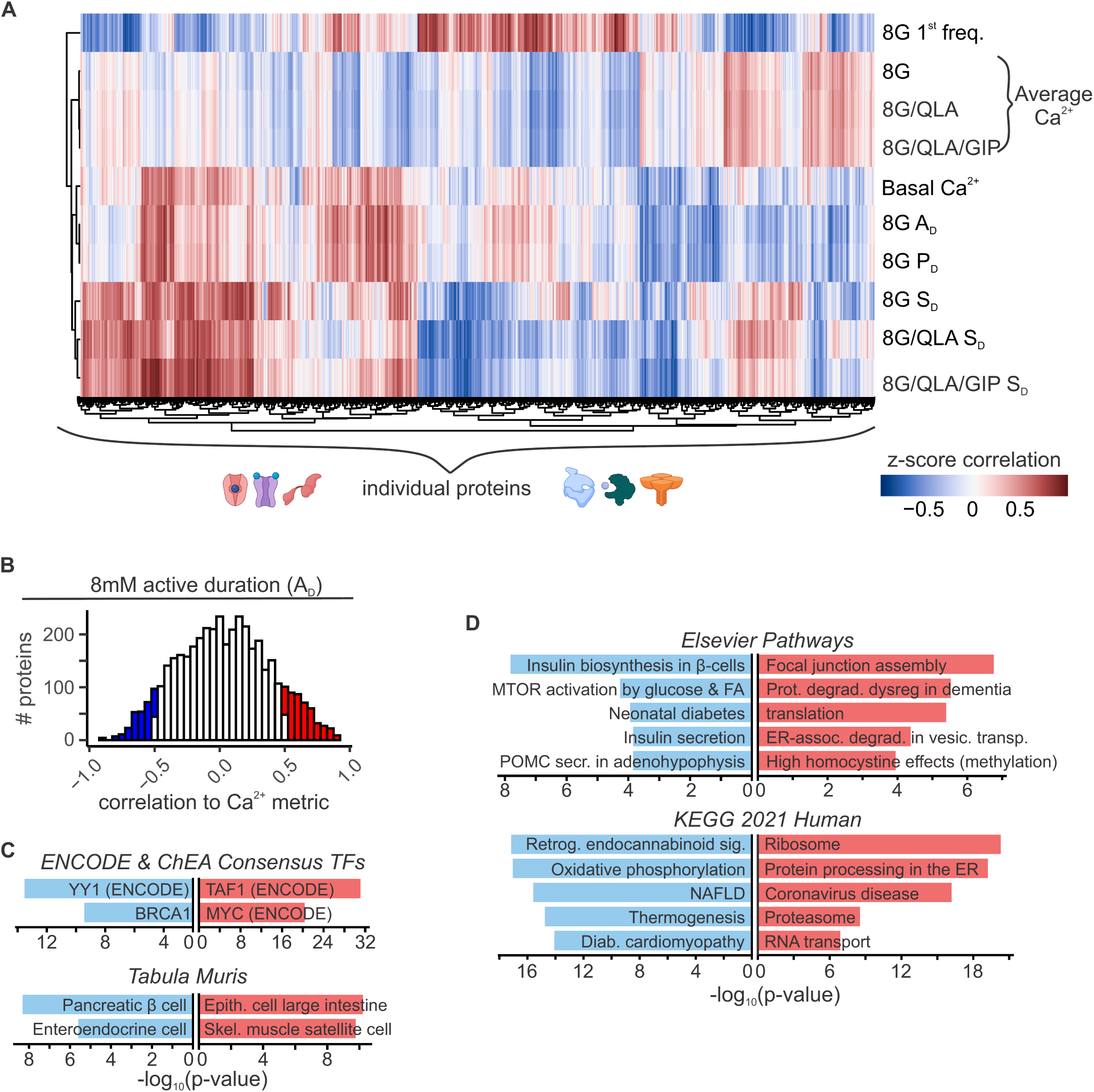
Islet proteins show correlation architecture to specific Ca^2+^ parameters. (**A**) Unsupervised clustering of correlation coefficients between protein abundance z-scores and z-scores for the Ca^2+^ parameters indicated. Islet proteins show differential correlation values to basal Ca^2+^, excitatory Ca^2+^ (detrended average values for 8mM, 8/QLA, and 8/QLA/GIP), active duration and pulse duration in 8mM glucose (8G P_D_ & A_D_), and silent durations (S_D_) in 8G, 8G/QLA, and 8G/QLA/GIP. Correlation coefficients for other parameters are indicated in **Supplemental Figure 8**. **(B)** Histograms representing the number of proteins that are correlated (red) and anti-correlated (blue) to 8G A_D_. TRRUST transcription factor motif database and ARCHS4 Tissue signature database (**C**) as well as pathway enrichments for the Elsevier Pathway database and KEGG 2021 Human pathway database (**D**) (-log_10_(p-values)), for the highly correlated (red) and anticorrelated (blue) proteins to 8 A_D_ metric. Databases were queried using Enrichr (37, 88).

Among the 4054 islet proteins, 363 had high absolute correlation coefficients (*r* > |0.5|) to 3 or more of the parameters our data suggest most strongly correlate to insulin secretion and plasma insulin (Basal Ca^2+^, 8G A_D_, 8G P_D_, 8G/QLA S_D_, 8G S_D_, 8G/QLA/GIP S_D_). Interestingly, of the proteins correlated to these traits, many have been previously implicated in islet biology, including PCSK1, GCK, SUR1, GLUT2, PDX1, and GLP-1 (28–36). Notably, the highly correlated proteins enriched for tissues, pathways, and transcription factors that support their role in insulin secretion (**Figure 5B, C, D**, Enrichr links in **Supplemental Table 1**) (37). For instance, proteins highly anti-correlated to active duration in 8G were enriched for components of oxidative metabolism and had their gene promoters enrich for binding to the islet transcription factor MAFA. These enrichment data provide a framework for discovering new genes of interest for their role in islet function.

### Integration of mouse genetics with human GWAS

The data presented in **Figure 5A** illustrates the *correlation* between islet proteins and Ca^2+^ dynamics. Importantly, a protein strongly correlated to Ca^2+^ does not necessarily reflect a causal relationship, i.e., a change in protein abundance may or may not cause a change in the Ca^2+^ signal. To take our analysis beyond correlation, we integrated our data with human GWAS of glycemia-related traits.

For each Ca^2+^ parameter, we focused on those proteins in the tails of the correlation histogram where *r* > |0.5| (e.g. **Figure 5B**). We identified human homologues for 3073 proteins that were correlated to Ca^2+^ in either direction for at least one of our parameters of interest. We then searched the Type 2 Diabetes Knowledge Portal (https://t2d.hugeamp.org/) for SNPs that are associated with one or more glycemia-related traits (see **Table 2**) with a P-value < 10^-8^, and are located within ±100 kbp of the homologous gene (e.g. COBLL1, **Figure 6A**), or in regions that contact the gene’s promoter region (determined using human islet promoter-capture HiC data (38)) as illustrated by ACP1 (**Figure 6B**). This yielded a list of 647 human genes strongly associated with diabetes-related SNPs. Among these genes, 478 were not previously associated with insulin secretion (see Methods), suggesting they may have understudied roles in islet function (**Figure 7A, B**; **Table 3**, **Table 4**). Our approach thereby leverages the genetic diversity of the eight CC founder strains and human GWAS for diabetes-related traits to highlight genes that may play a novel role in islet function and relate to diabetes risk.

**Table 3.**
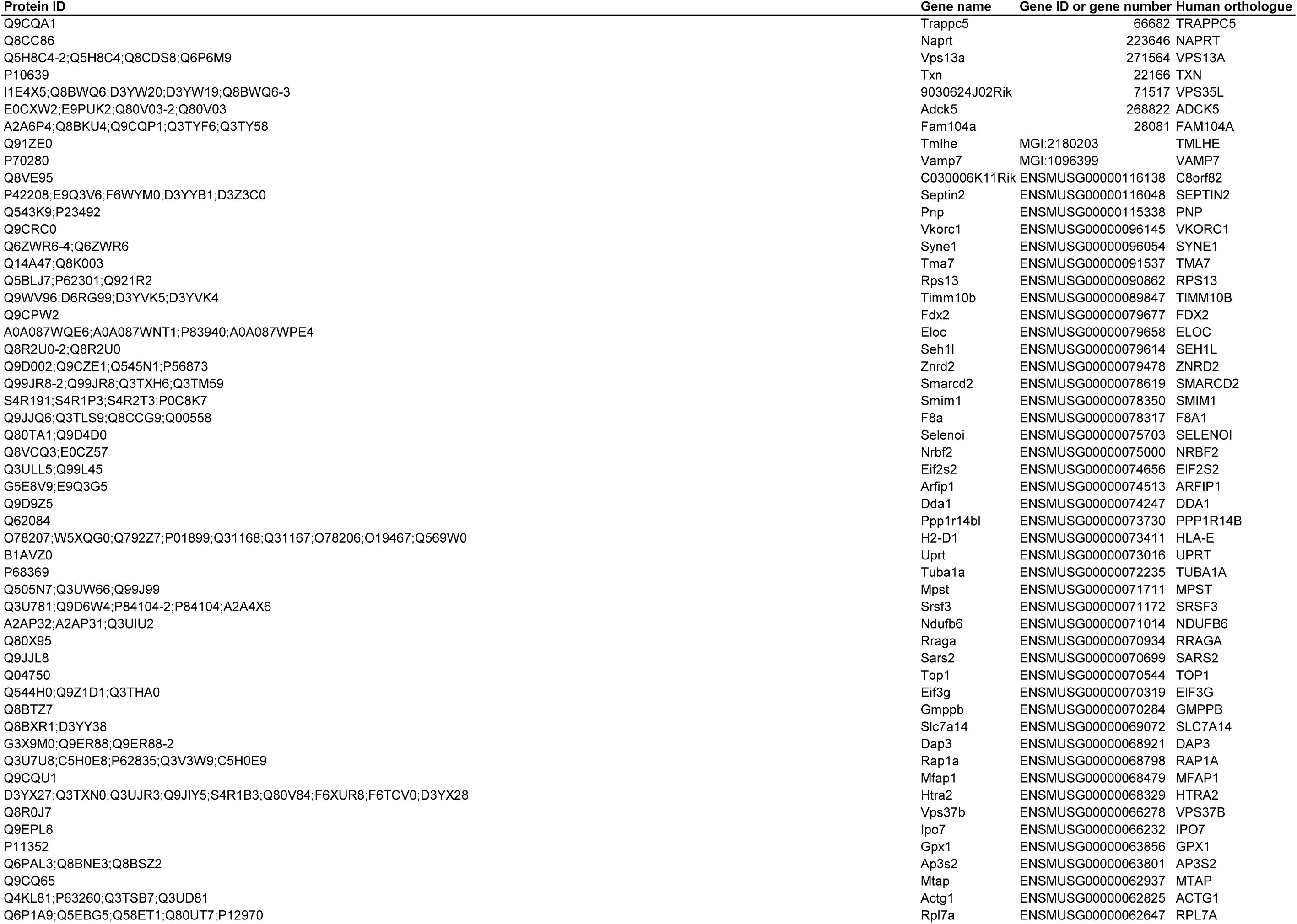
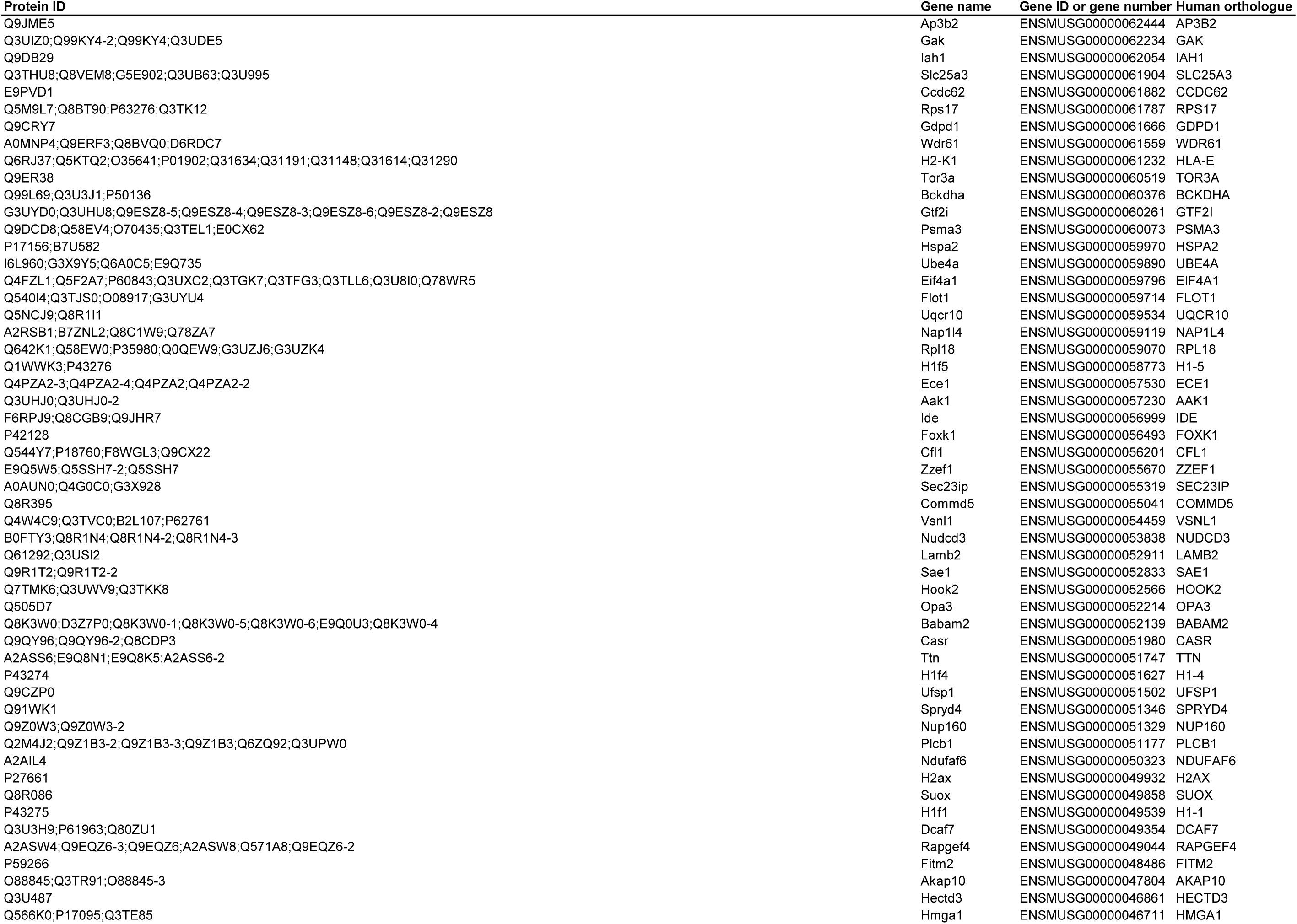
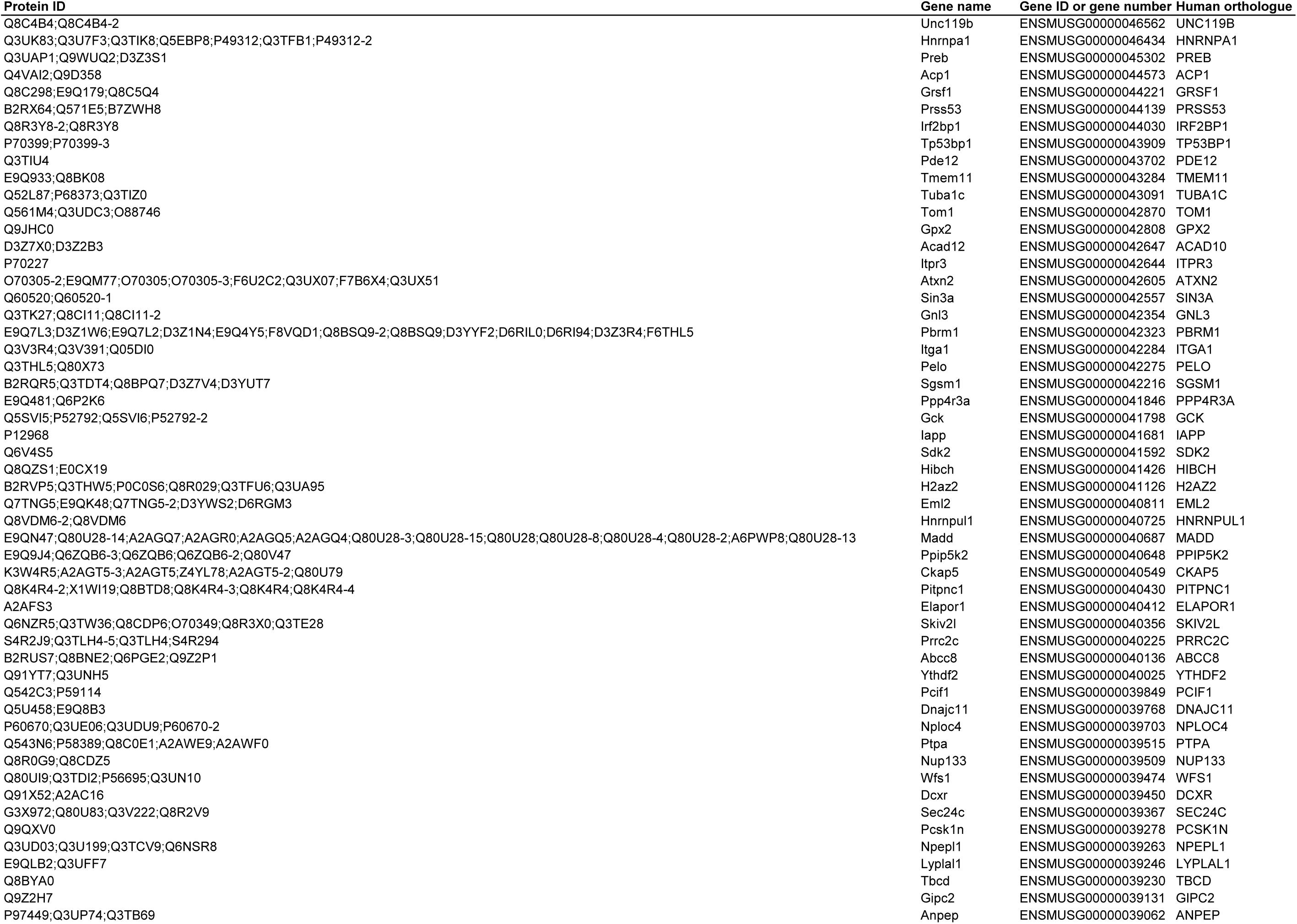
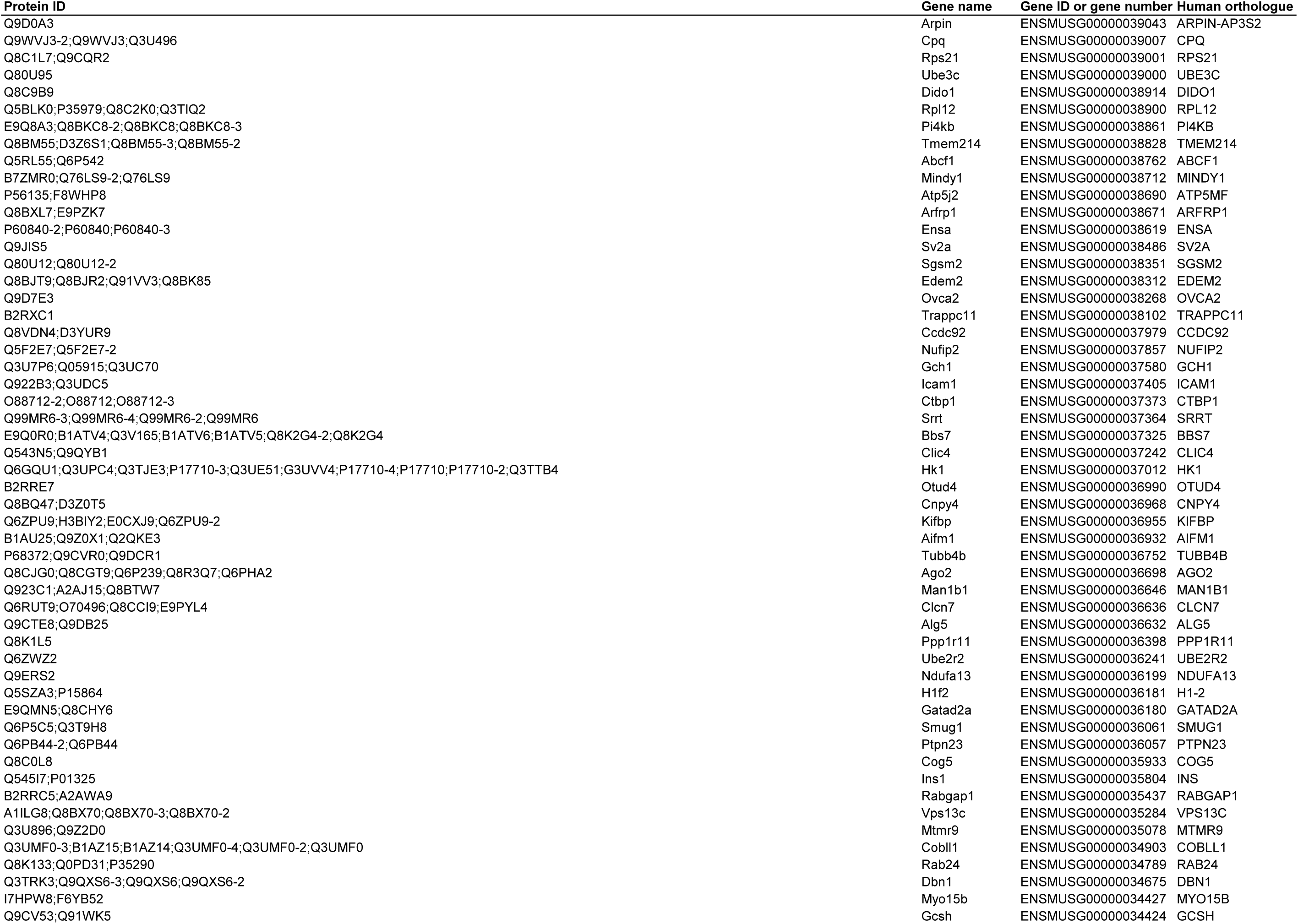
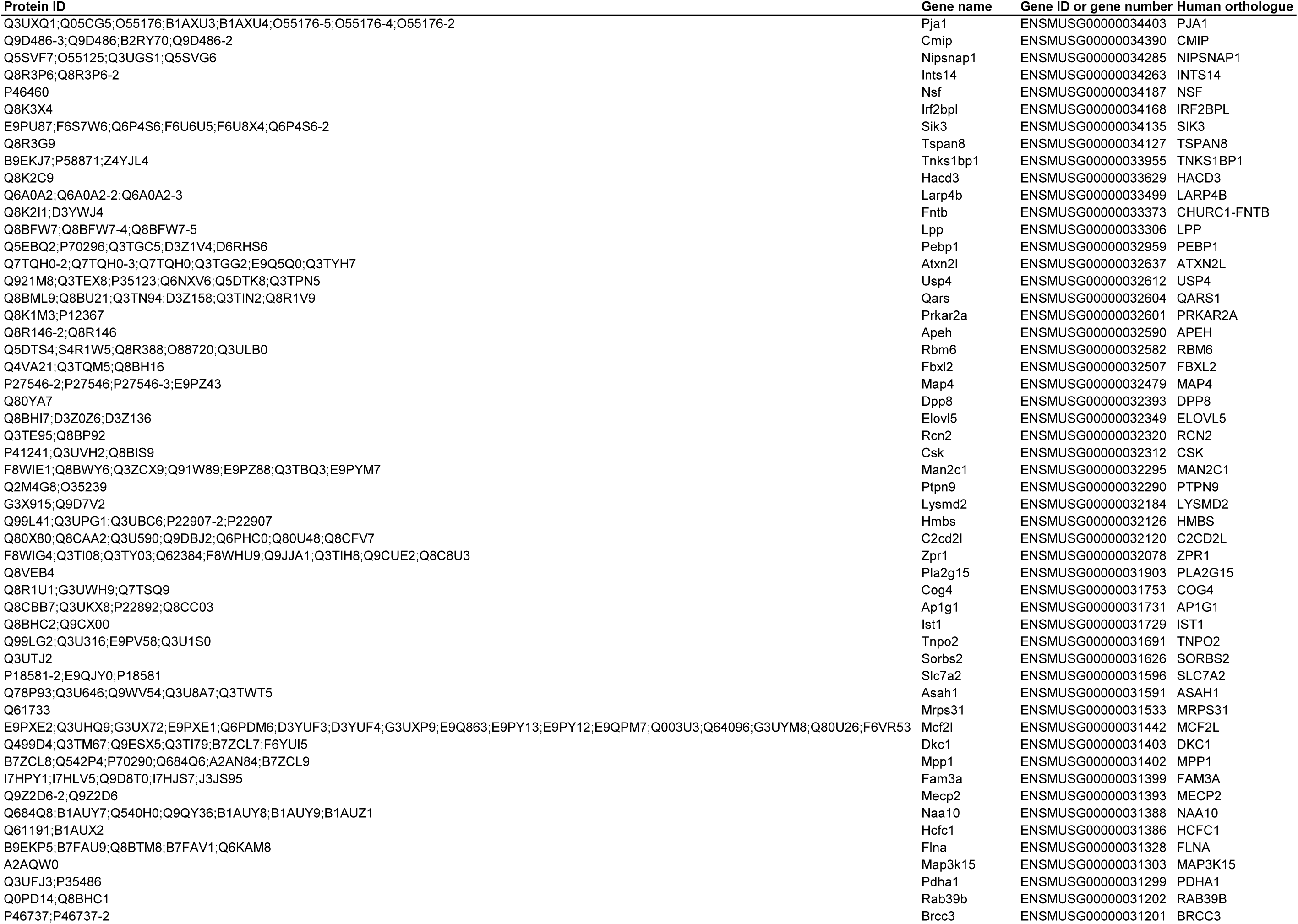
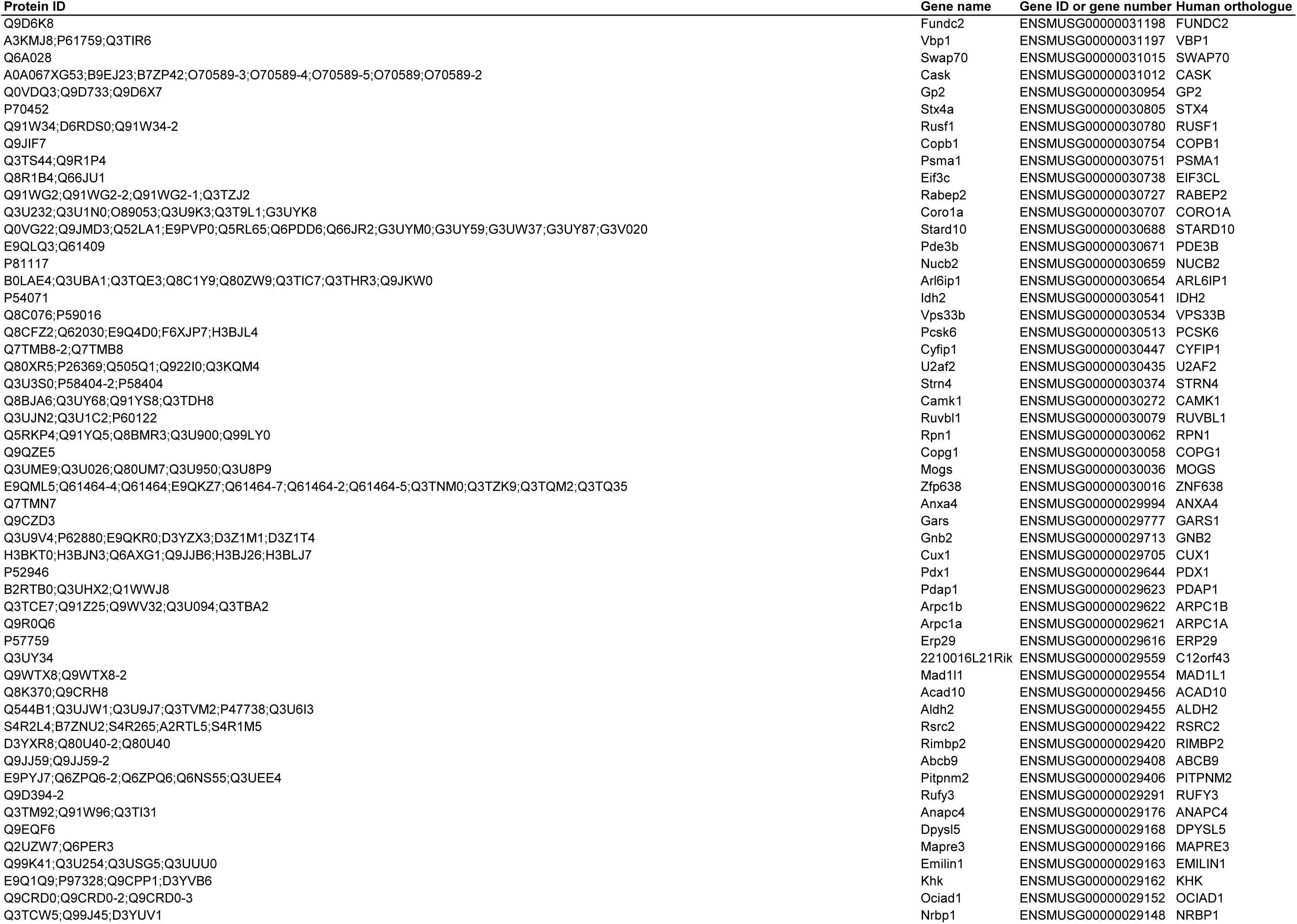
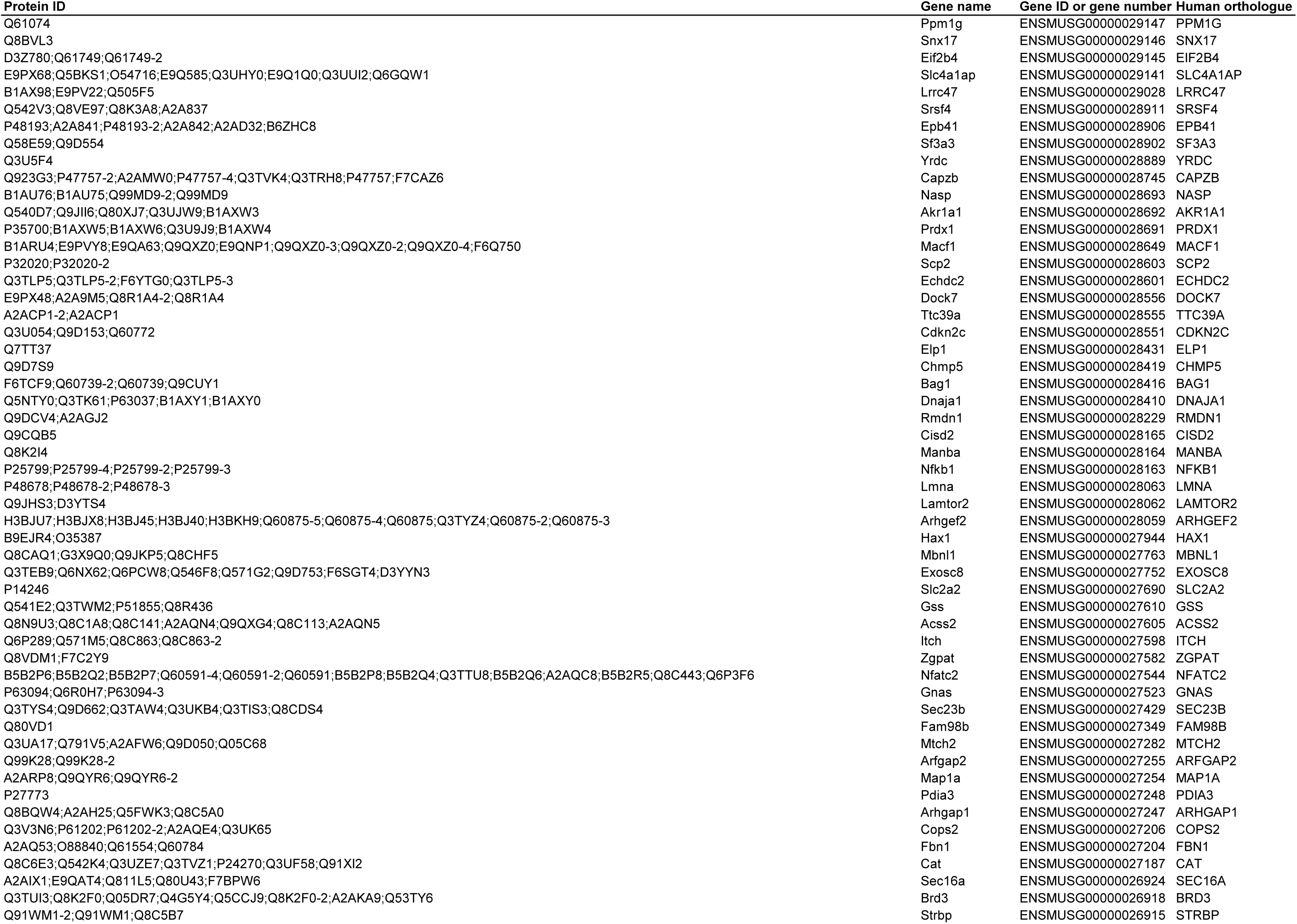
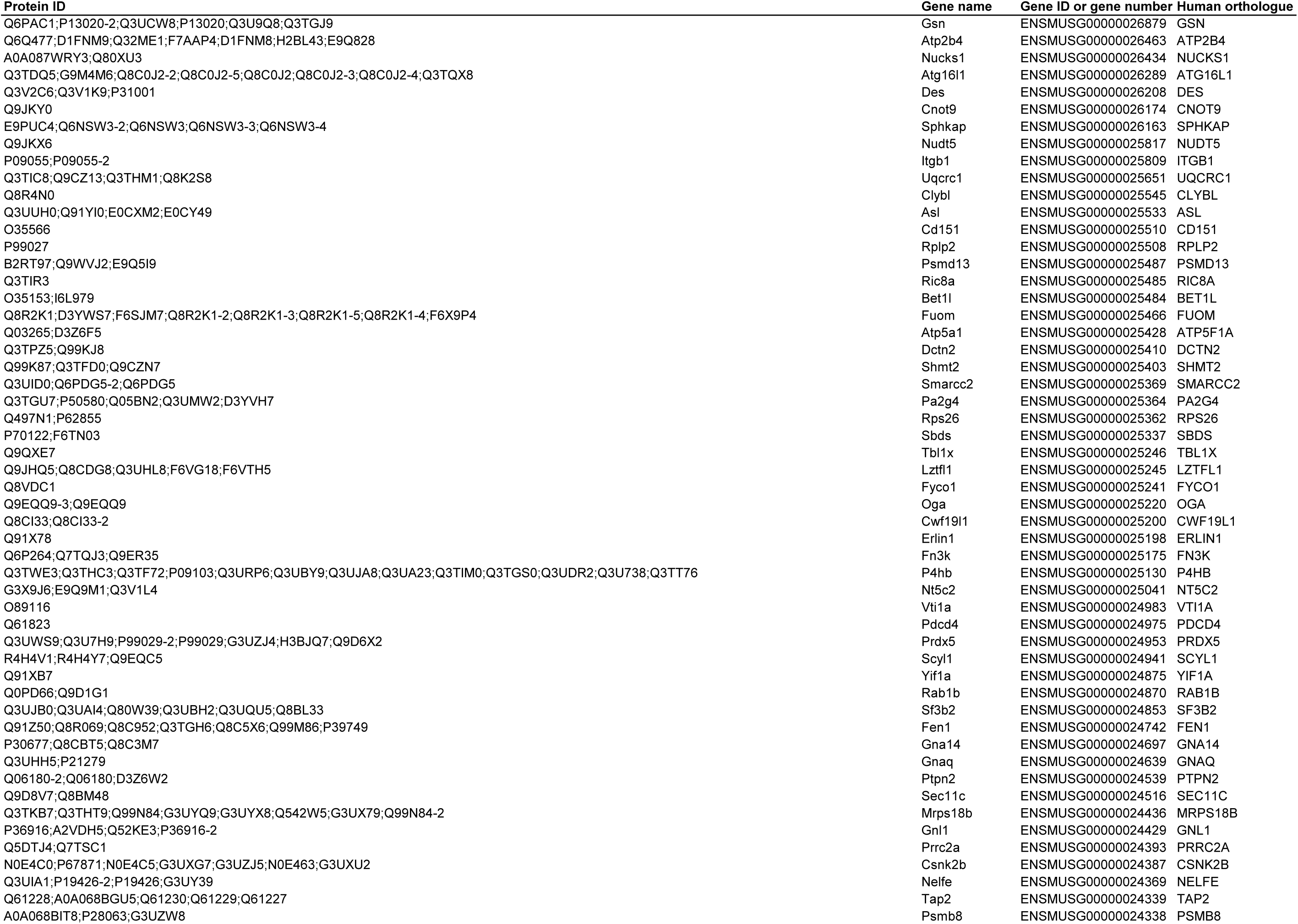
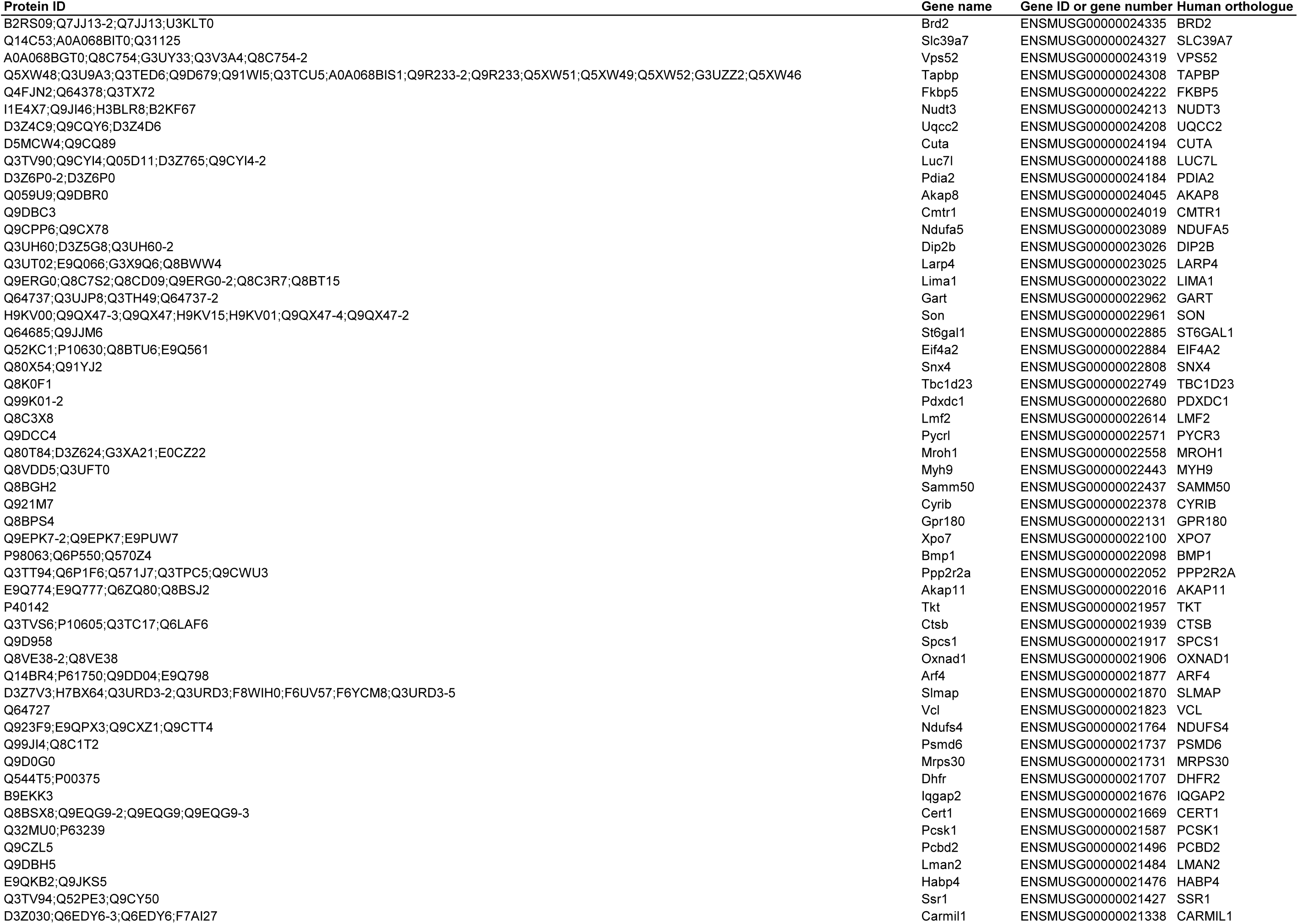
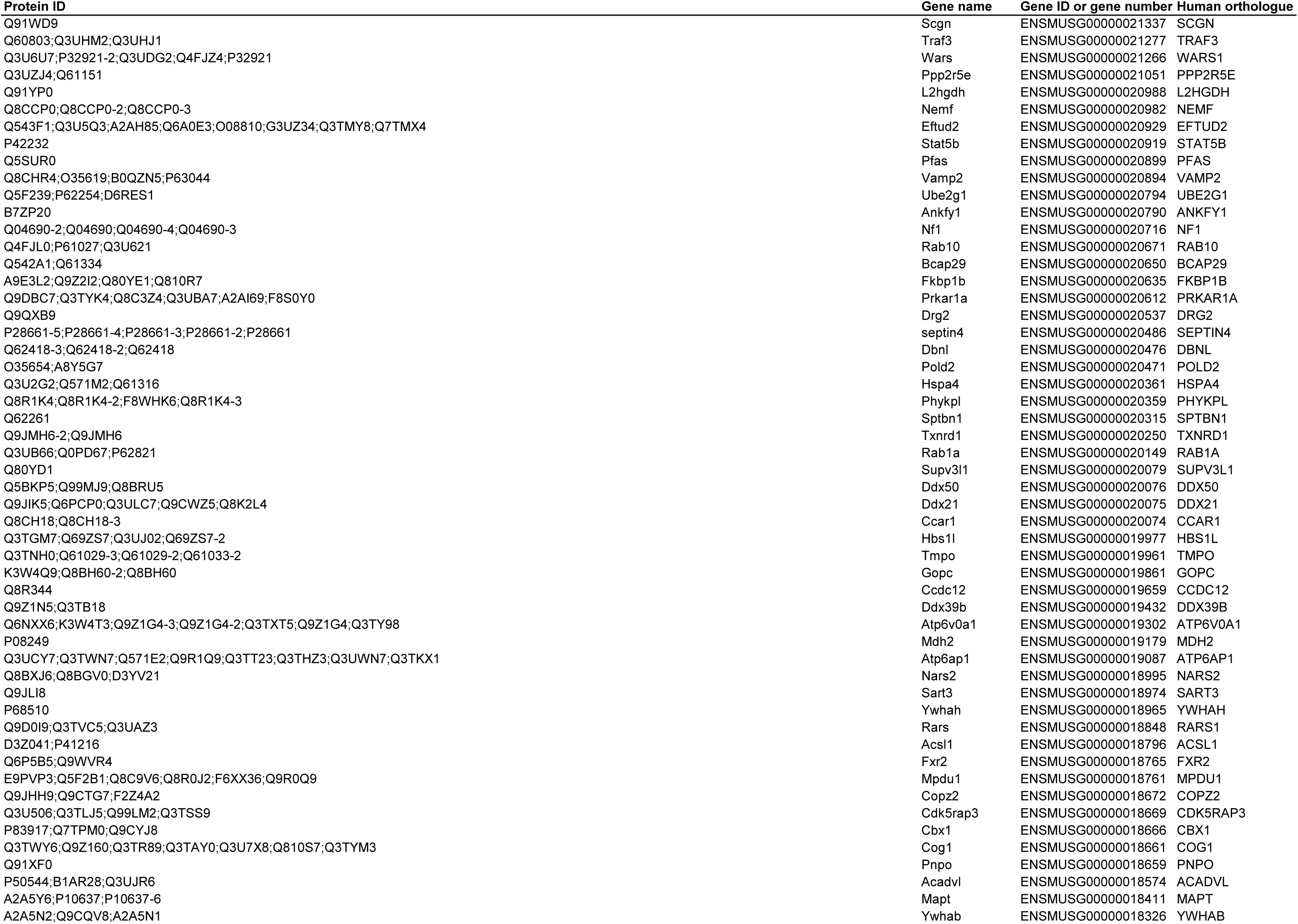
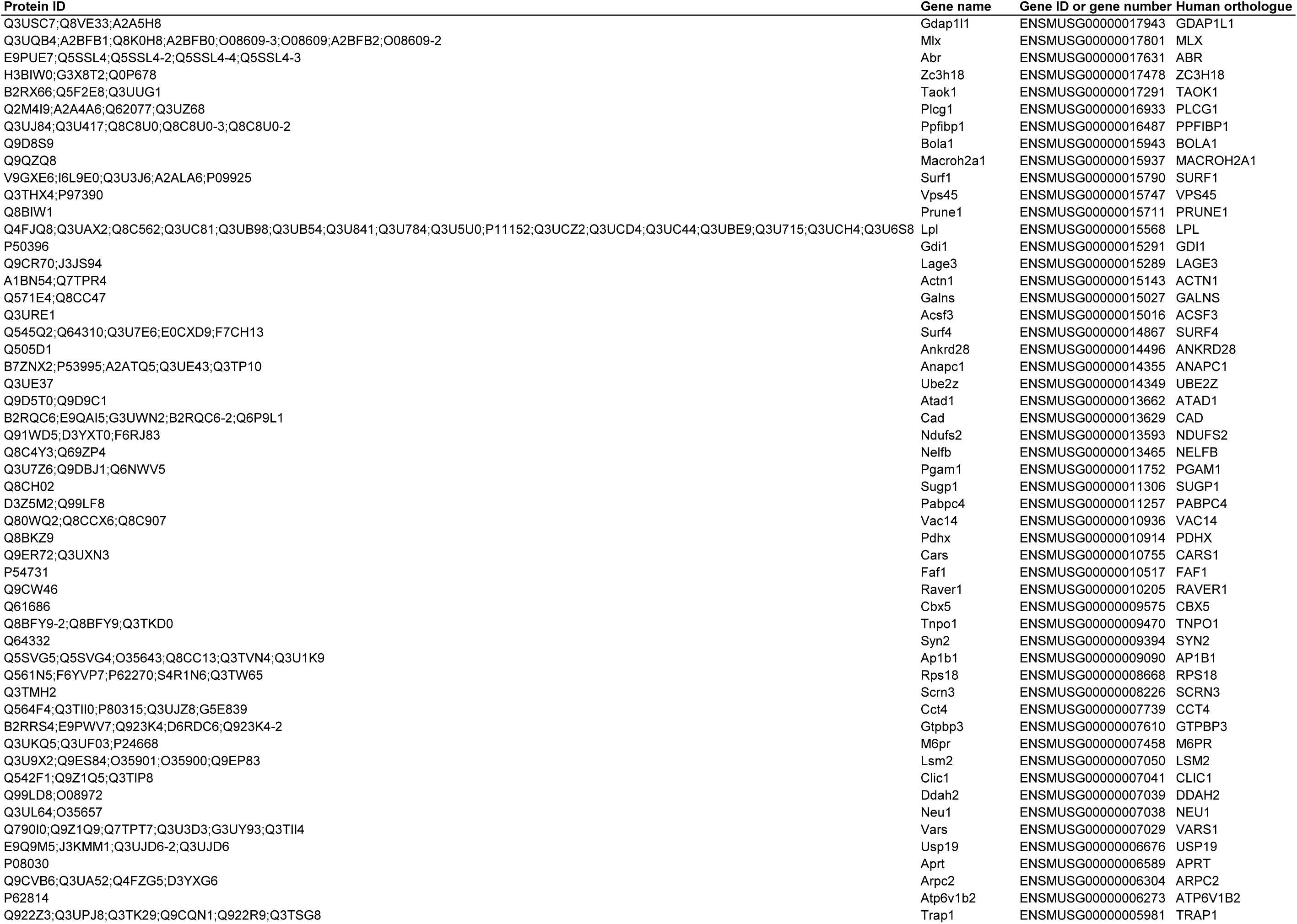
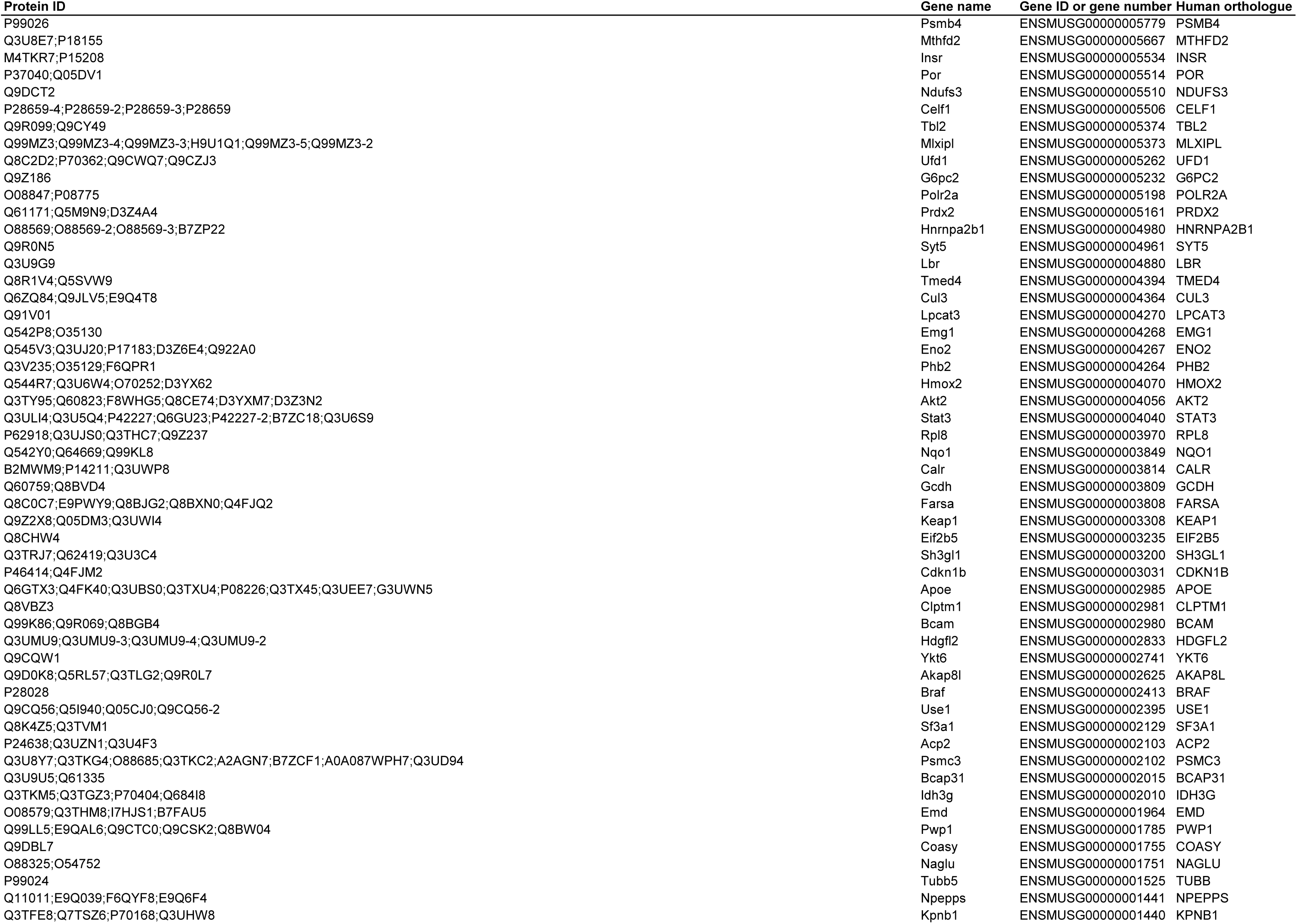
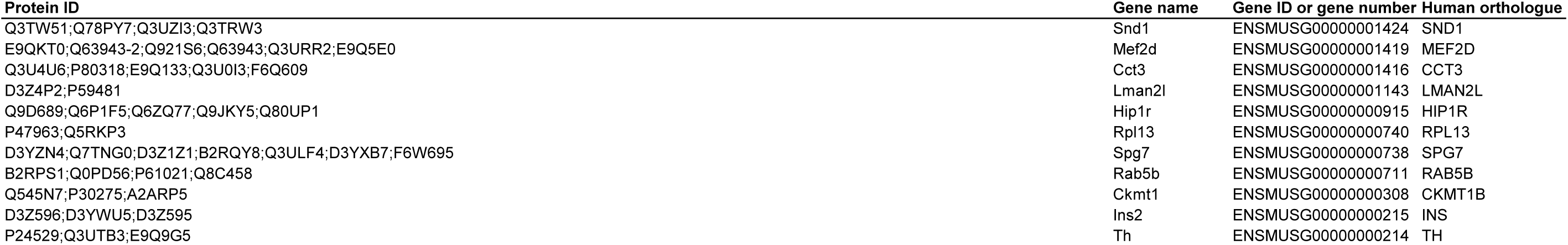
(on following pages): Proteins correlated with Ca^2+^ parameters that have glycemic-related SNPs. This includes protein IDs, gene names, gene IDs, and human orthologues for each of the proteins that correlate to one of the following metrics and have a glycemic-related SNP (see Table 2): basal Ca^2+^, 8G S_D_, 8G/QLA S_D_, 8G/QLA/GIP S_D_, 8G A_D_, 8G P_D_, and 8G 1^st^ freq.

**Figure 6.**
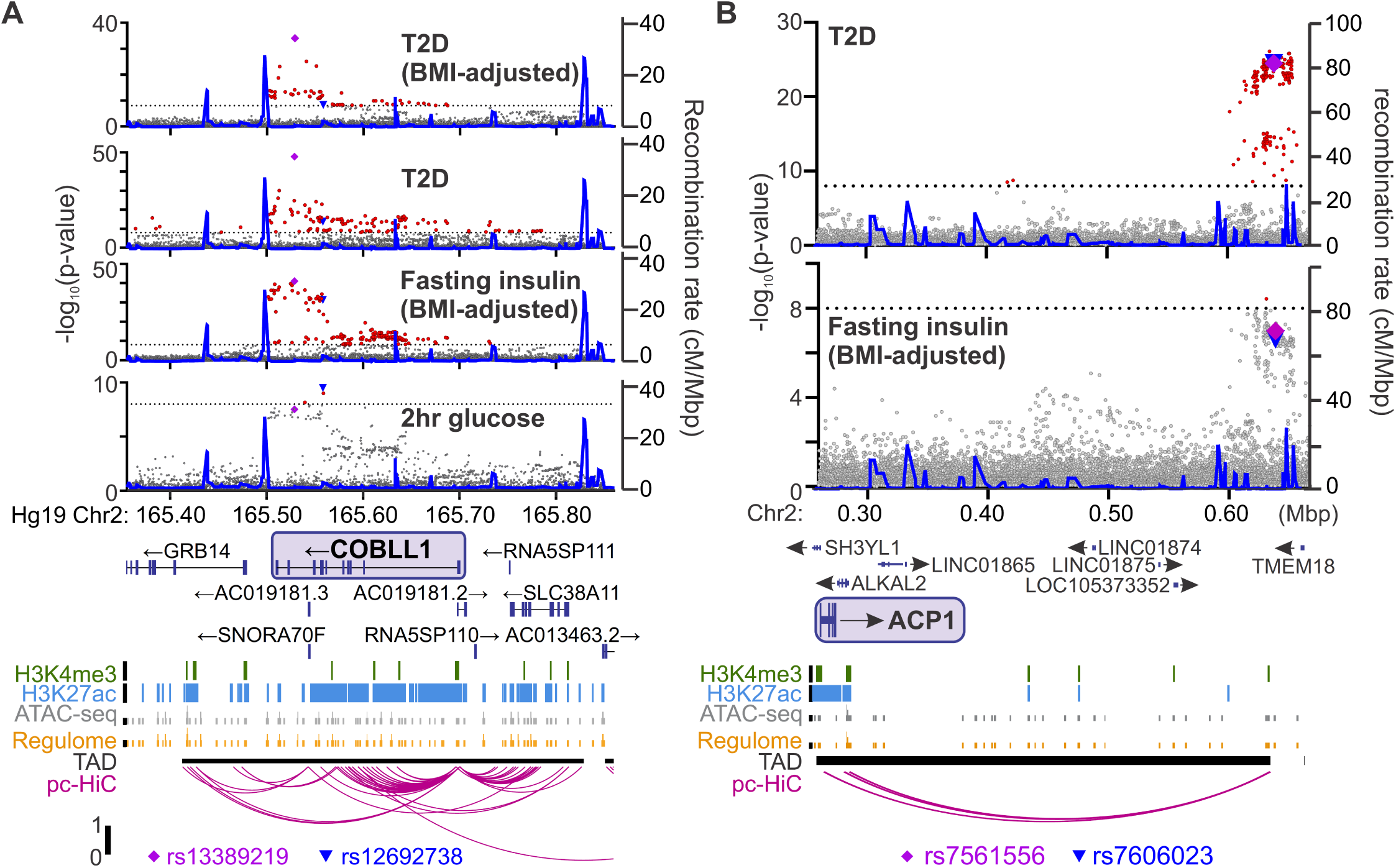
Identifying candidate protein targets by integrating human GWAS. (**A**) An example gene, *COBLL1*, orthologous to a gene coding for a protein identified as highly correlated to Ca^2+^ wave parameters in the founder mice. The recombination rate is indicated by the solid blue line. Significant SNPs (8 < -log_10_(p), red) decorate the gene body for multiple glycemia-related parameters (in bold). Human islet chromatin data (38) for histone methylation (H3K4me3), histone acetylation (H3K27ac), ATAC-sequencing (ATAC-seq), and regulome score suggest active transcription of the gene within a topologically-associated domain (TAD). Human islet promoter-capture HiC data (pc-HiC) (38) show contacts between the SNP-containing regions decorating the gene and its promoter. The highest SNP for 2hr glucose 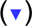 and the other parameters 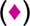 are indicated. (**B**) Some orthologues did not show SNPs decorating the gene itself but did show looping to regions with SNPs for glycemic traits. The promoter of *ACP1*, for example, loops to a region within its topologically associated domain (black bar) with strong SNPs for type 2 diabetes risk and near-threshold SNPs for fasting insulin adjusted for BMI. Some SNPs 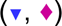 lie directly on the contact regions identified by HiC, whereas others lie immediately proximal to these contacts. For both panels, the significance of association (-log_10_ of the p-value) for the individual SNPs is on the left y axis and the recombination rate per megabasepair (Mbp) is on the right y axis. Chromosomal position in Mbp is aligned to Hg19. SNP data were provided by the Common Metabolic Diseases Knowledge Portal (cmdkp.org).

**Figure 7:**
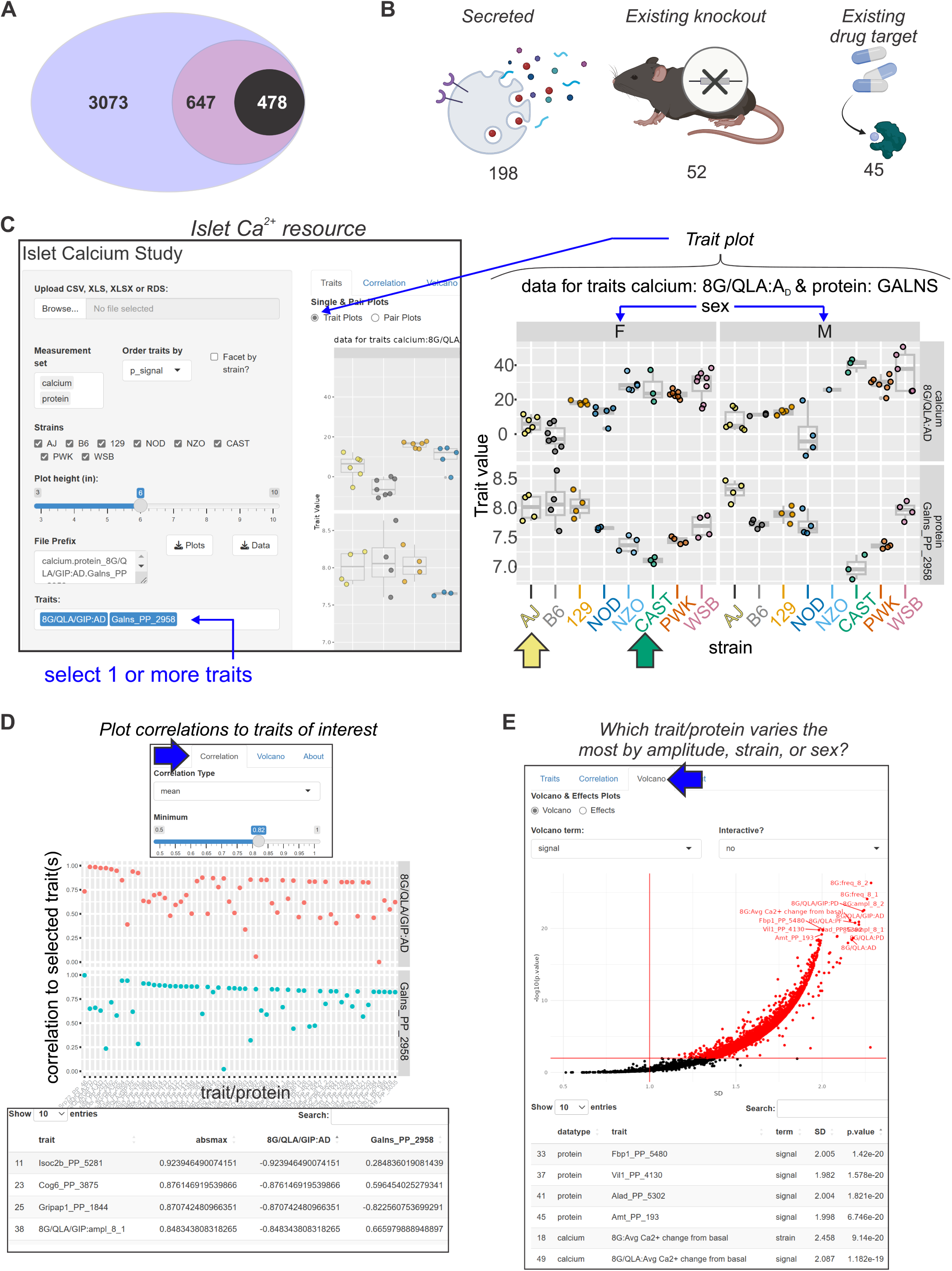
Mining Ca^2+^ data using a novel online resource (preceding page) (**A**) 3073 islet proteins significantly correlated to islet Ca^2+^ parameters of interest. Among the proteins, 647 had orthologues containing SNPs for glycemic traits. Of these, 478 showed no results in our starting triage (see Methods) under any alias, suggesting they may be understudied in islet biology. (**B**) Of these 478 proteins, 198 were found to be secreted either as soluble proteins or in exosomes (68, 69, 71–75), 52 have existing knockout mice with annotated glycemia or pancreatic phenotypes (76, 95), and 45 have existing compounds that target them (62–67). To make these data more accessible, we have developed an online resource that enables individuals to query the Ca^2+^ and proteomic data simultaneously. The user can select proteins and calcium traits (**C**) and display strain and sex distribution of these traits to determine the ideal backgrounds on which to test their traits or proteins of interest. In this example, GALNS is highly correlated to 8G/QLA/GIP A_D_, with the highest and lowest abundance strains for GALNS being AJ (yellow arrow) and CAST (green arrow), respectively. (**D**) The user can also query for the correlations between Ca^2+^ traits and proteins against one another or other traits of the same category. (**E**) The user can also see which of the traits or proteins has the largest change and most significant effects by sex, strain, or sex and strain.

**Table 4.**
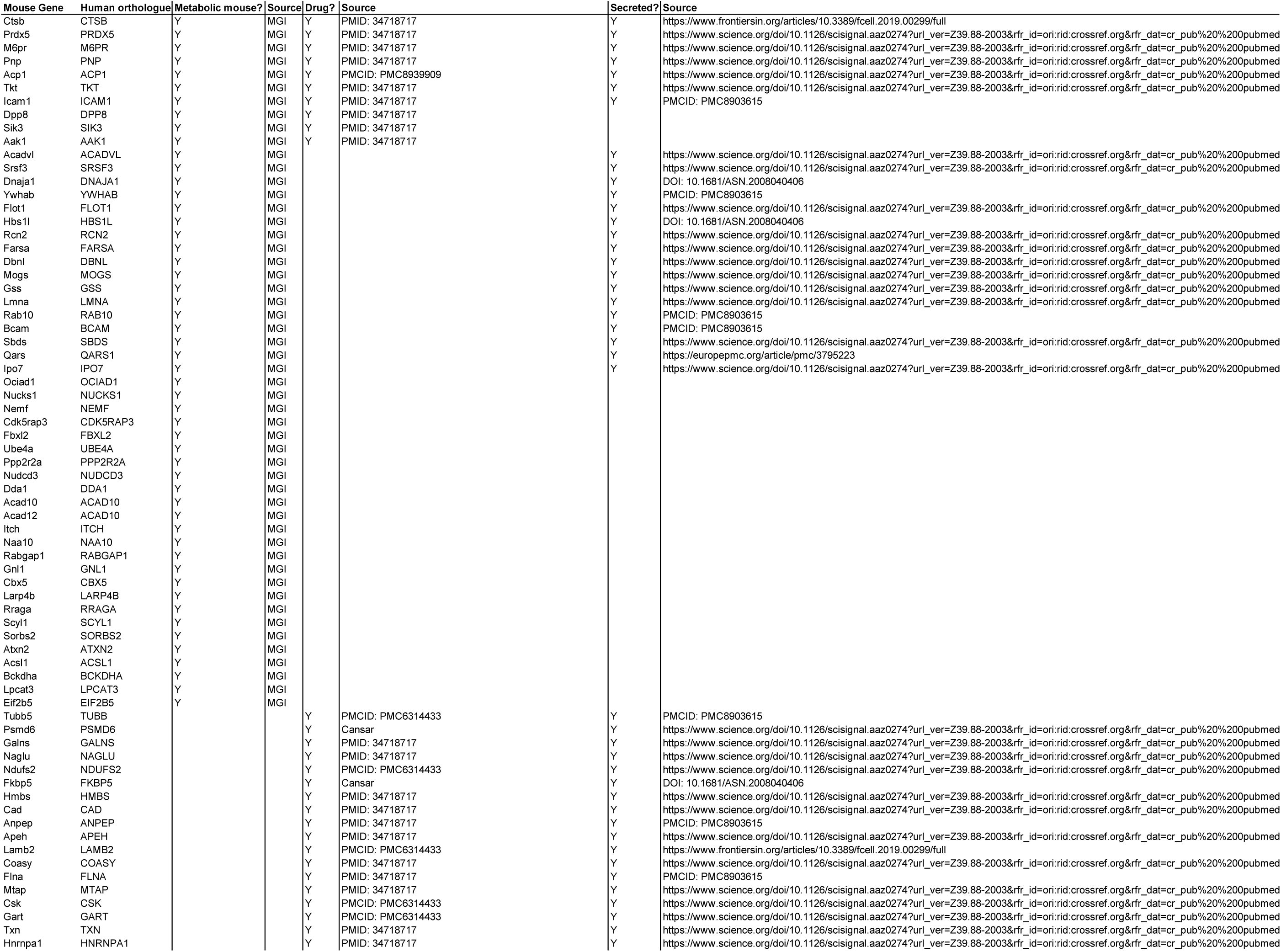
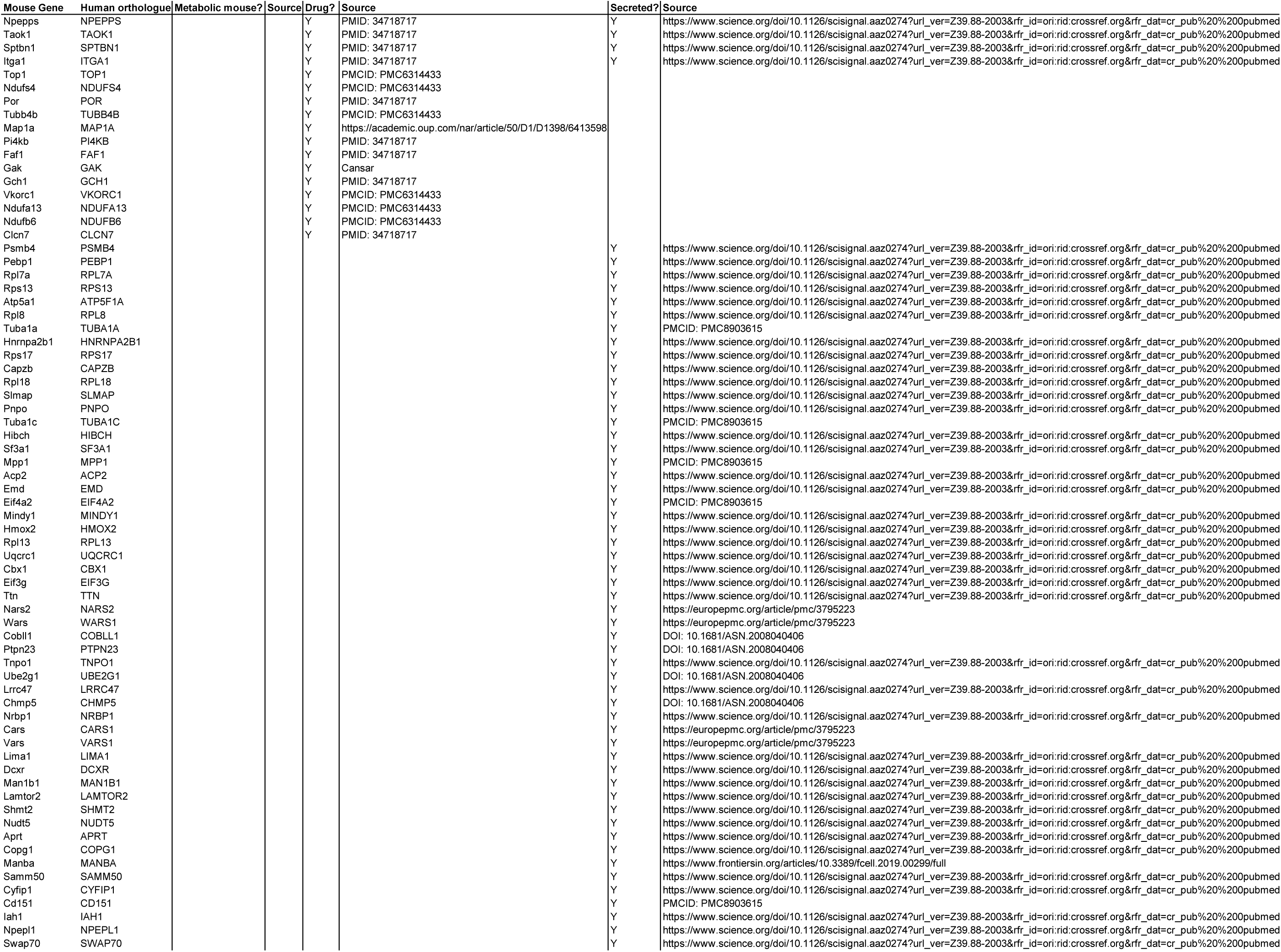
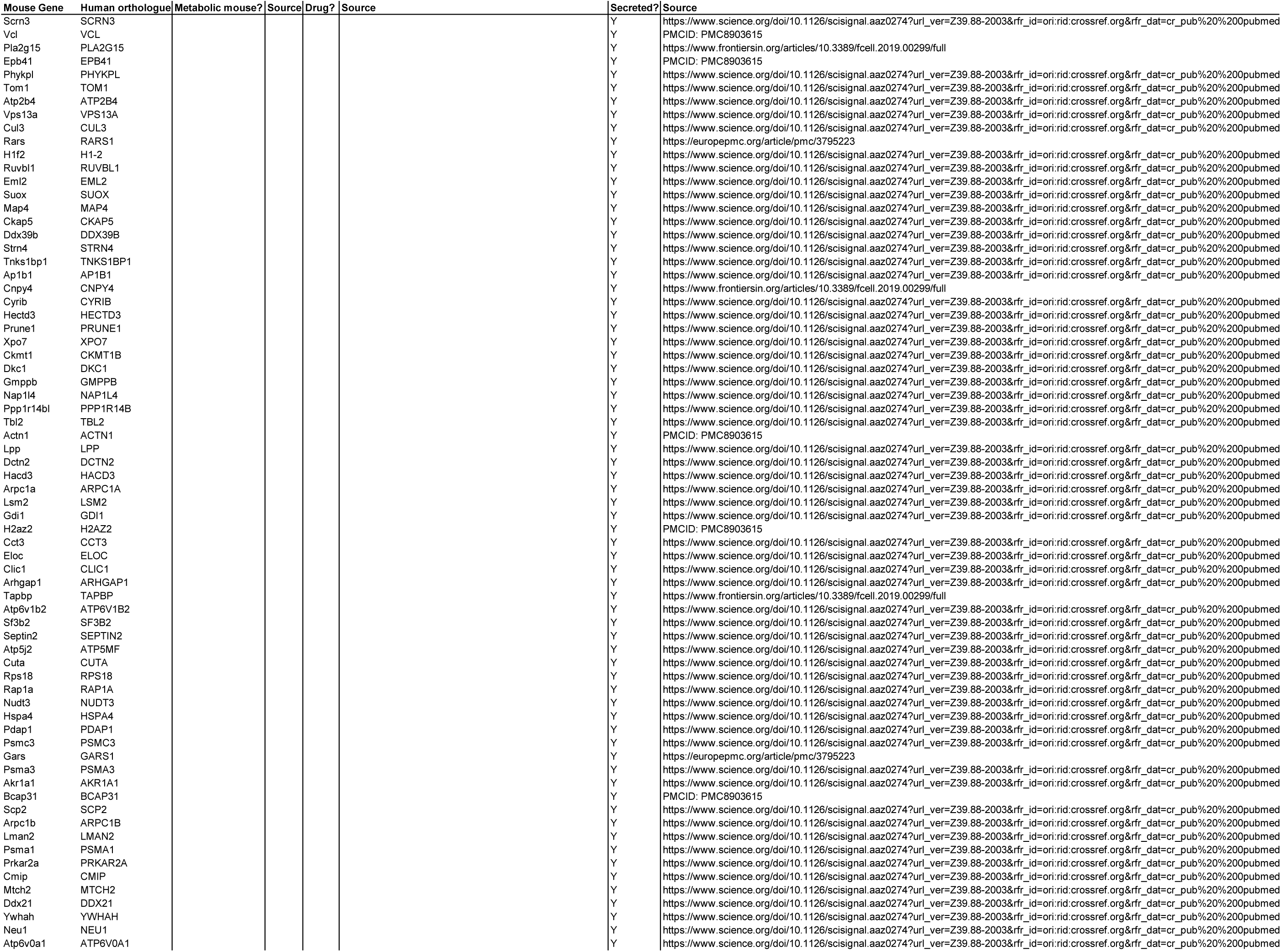
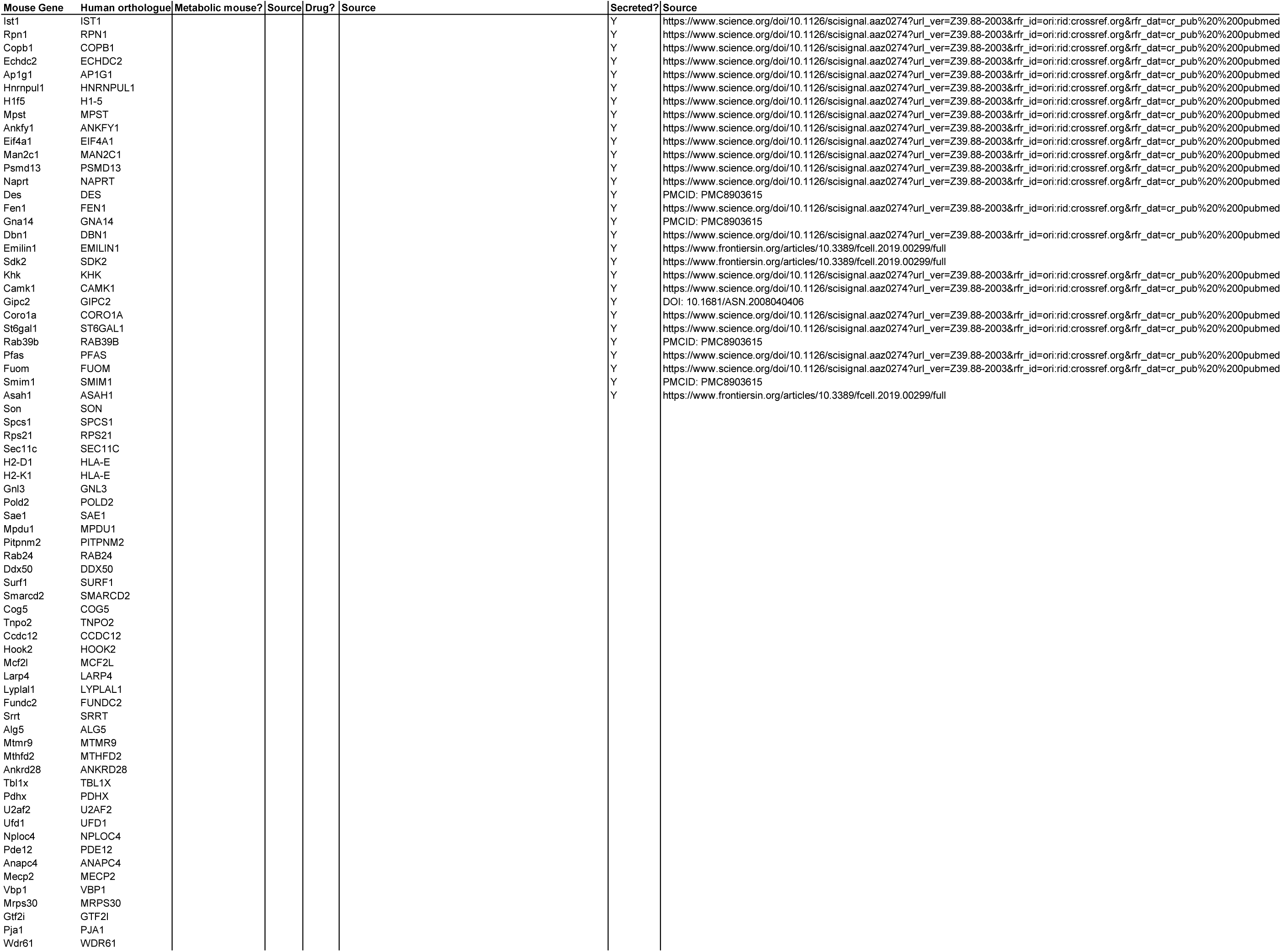
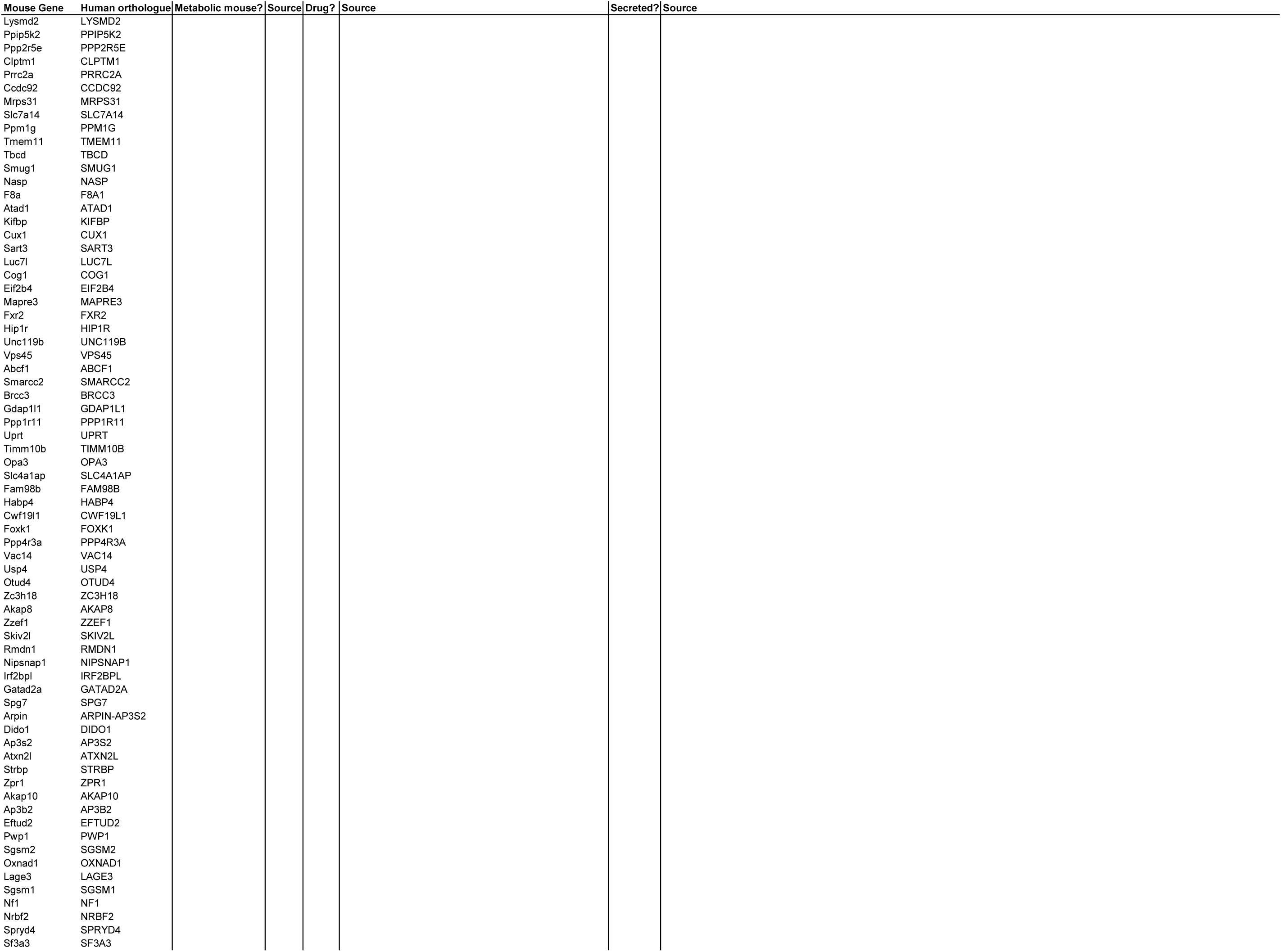
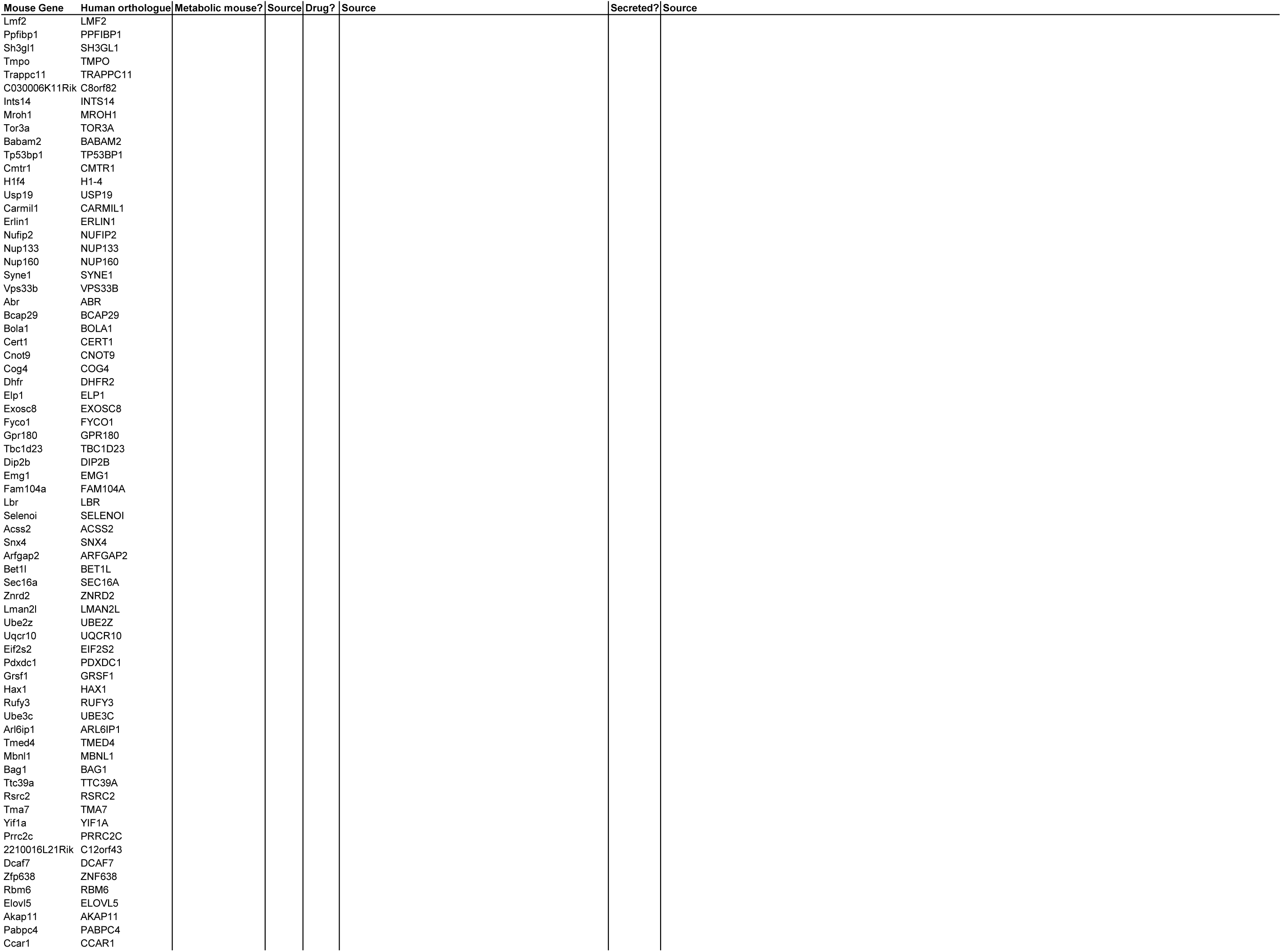
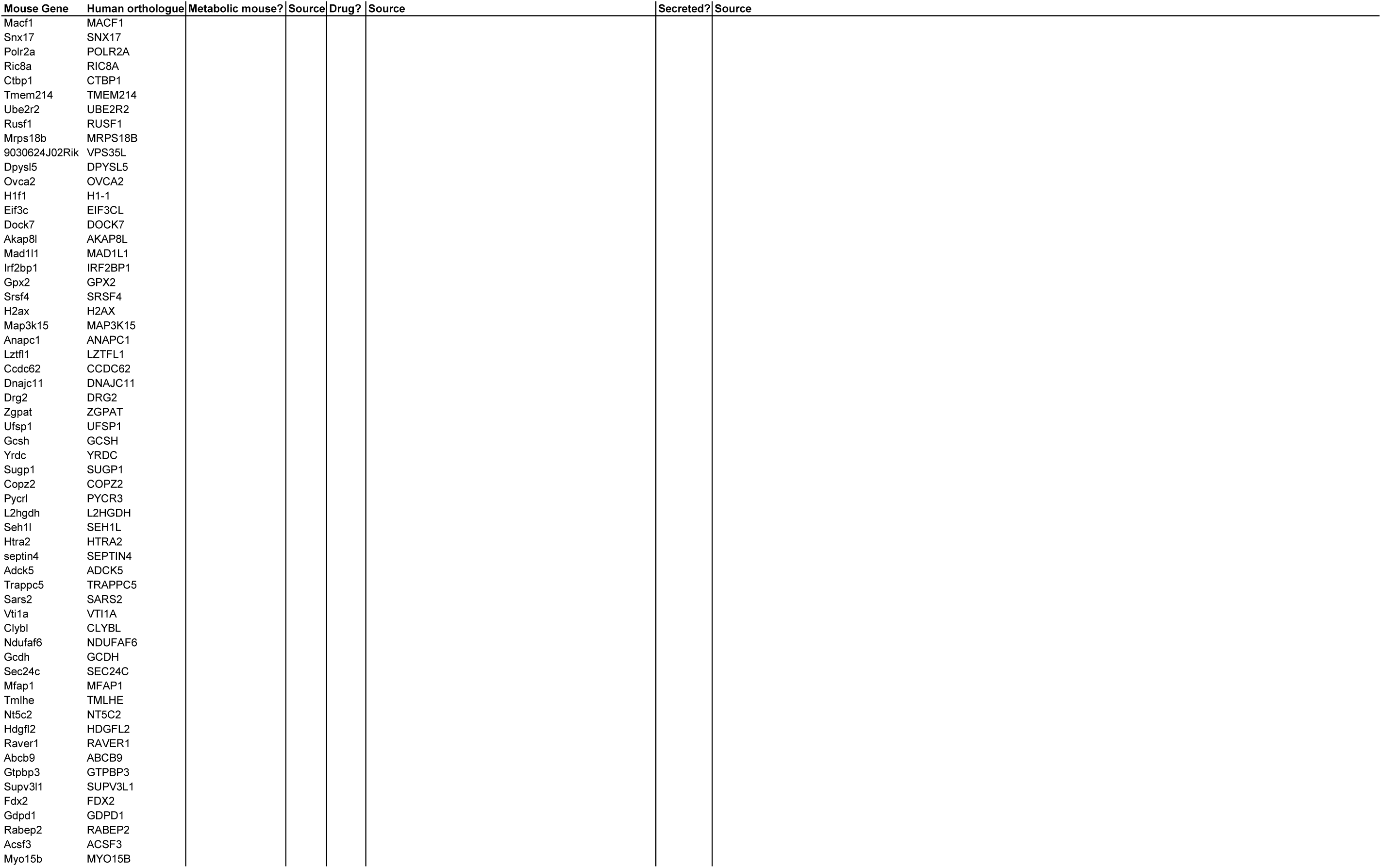
(on following pages): Proteins understudied in islet biology. This table of proteins indicates the subset of proteins meeting our selection criteria (Table 3) that did not have any results in Pubmed, Google Scholar, or Google for any alias and the term “insulin secretion,” suggesting that they may be understudied in islet biology. The gene symbols for the mouse gene and human orthologue are indicated. The “Mouse?” column indicates whether a knockout mouse with metabolic phenotypes exists (identified from sources in the subsequent “Source” column). The “Drug?” column similarly indicates whether any source (indicated in the following “Source” column) shows existing compound(s) targeting the protein. Finally, the “Secreted?” column indicates whether any source (indicated in the subsequent “Source” column) shows an isoform of the protein to be secreted.

To aid in the selection of mouse strains for validating potential candidate regulators, we provide a resource with proteomic and Ca^2+^ data (**Figure 7C, D, E**; https://data-viz.it.wisc.edu/FounderCalciumStudy/, https://doi.org/10.5061/dryad.j0zpc86jc, https://rstudio.it.wisc.edu/FounderCalciumStudy). This will enable the user to identify proteins correlated to other proteins or traits of interest, and from there, identify which strain(s) may be most appropriate for studies of their protein of interest. In the examples illustrated in **Figure 7C**, the user queries for the strain/sex distribution of the protein GALNS, which shows a high negative correlation to multiple traits, including the active duration time in 8G/QLA. Strains at the extremes of this trait are also extremes regarding GALNS protein abundance. Strains with high abundance (AJ, for example (yellow arrow)) would be ideal models for inhibition or knockout, whereas CAST mice (green arrow) could be a comparison strain for validating the role of the protein, as they express much less GALNS. Users can also query for the proteins or calcium parameters and see their correlation to the other proteins and/or calcium parameters (**Figure 7D**). Importantly, this includes the ability to look at the correlations between traits for individual strains or subsets of strains, enabling the user to see how the main clusters of strains (classic, wild-derived, and disease models) or individual outlier strains are drivers of specific traits. Finally, the user can see which traits or proteins show the strongest strain, sex, or sex-by-strain effects using the options in the volcano plot (**Figure 7E**). Together, these and other tools in the resource will allow researchers to explore their traits or proteins of interest as well as determine the appropriate model systems and conditions that may best interrogate their experimental questions of interest.

## DISCUSSION

### Genetic variability drives islet function

While the development and progression of T2D is potentiated by environmental factors, an estimated 50% of disease risk is driven by genetic factors (1, 3, 39). Therefore, to study the genetic variation contributing to T2D, we took advantage of the genetic diversity contained within the eight CC founder strains. These mice collectively contain a level of genetic diversity mirroring that seen in humans, making them an excellent experimental platform to link genetics with altered islet function (14, 15). We demonstrate that they also vary in their Ca^2+^ response to various insulin secretagogues, supporting the use of these mice to identify novel genes involved in regulating islet biology.

### Dissecting the calcium waveform highlights islet regulatory pathways

Variations in Ca^2+^ dynamics are highly complex, reflecting changes in metabolism, extra-islet signaling, and Ca^2+^ itself (8). We therefore selected stimulatory conditions to assess each of these components in islets of the eight mouse strains. 8 mM glucose was first used to survey glycolytic responses, because we have observed that several strains reliably oscillate at this glucose concentration. Furthermore, this glucose concentration remains close to the stimulatory threshold, thus reducing the possibility of oscillations plateauing if islet Ca^2+^ responses were left-shifted in any strains (23, 40, 41). We then added QLA as fuel to engage mitochondrial metabolism and paracrine signaling from α-cells, providing a survey of α-to-β cell communication in the islet (6, 7, 42, 43). Finally, we used GIP to interrogate the islet incretin responses and the cAMP amplification pathway (42), before returning to a low glucose concentration, which enabled us to establish baseline Ca^2+^ levels.

The variation in Ca^2+^ response to these conditions can be better understood by examining the multitude of pathways regulating Ca^2+^ dynamics. As mentioned previously, altering ionic pathways involved in regulating Ca^2+^, such as K_ATP_ channels, has a very different effect on Ca^2+^ oscillations compared to altering glycolysis. It is important to further dissect these pathways, as changing specific components of glucose metabolism can elicit different effects. For example, activating glucokinase (GK, which is rate limiting for glycolysis) and activating pyruvate kinase (PK, an ATP-generating enzyme which directly controls K_ATP_ channel closure in the final step of glycolysis) both reduce the silent duration by accelerating K_ATP_ channel closure. GK and PK activation also alter the active duration and the plateau fraction, however they do so in opposite directions (6). Activating GK increases the active duration and oscillation period, while activating PK decreases those same parameters via Ca^2+^ extrusion (7). The incretin hormones GLP-1 and GIP reduce silent duration and oscillation period, most likely due to their ability to activate Epac2 and sensitize K_ATP_ channels to ATP-dependent closure (44, 45). Thus, PK and GLP1 have a common target (i.e. K_ATP_), and therefore a similar effect on silent duration. These examples illustrate the benefit of analyzing multiple Ca^2+^ parameters for understanding pathways of interest.

The importance of analyzing a variety of Ca^2+^ parameters is further supported by the insulin secretion measurements in the male WSB and 129 mice. While average Ca^2+^ is a common metric used to predict insulin secretion, relying on only this metric would suggest that the two strains secrete insulin similarly **(Supplemental Figure 4)**. However, the WSB mice secreted significantly more insulin in 8G, 8G/QLA, and 8G/QLA/GIP (**Figure 3**). Based on our correlation analysis between Ca^2+^ parameters and insulin secretion across each sex and strain, active duration, and pulse duration in 8G more accurately predicted insulin secretion and may be highly informative when used with other data (**Figure 4**). This is similar to results published by some other groups, suggesting that average Ca^2+^ does not correlate well with insulin secretion (46).

### Strains segregate by their phylogenetic origins

Notably, several of the strains appeared to cluster together with similar responses. One such group is composed of three classical strains (A/J, B6, and 129), which had relatively similar waveforms that were dominated by slow oscillations. These differed from a second group containing the wild-derived strains (CAST, WSB, and PWK) which closely matched one another and were dominated by faster oscillations. Additionally, even with the clear separation between the clusters, inter-strain variation was still observed within the clusters (e.g. more 129 islets had plateau responses to 8G/QLA than the B6 or AJ).

The classic strains have been highly inbred (>150+ generations) and descend from related common ancestors, the “fancy mice.” They also have extremely low genetic diversity, with 97% of their genomes explained by fewer than 10 haplotypes (3, 47, 48). By contrast, wild-derived strains are each independent in their parental origin, inbred for far fewer generations than the classic strains (20 vs. >150), and include significant contributions from other subspecies of *Mus musculus* than the predominant subspecies (*M. m. domesticus*) in the classic strains, particularly CAST (*M. m. castaneous)* and PWK (*M. m. musculus*) (3, 47, 48). It is thus unsurprising that the two primary Ca^2+^ response clusters were composed of the classic and wild-derived strains. Multiple loci have already been linked to islet dysfunction and differential metabolic homeostasis in the classic strains (3). Our work here highlights the promise in using wild-derived strains to unmask previously underappreciated islet phenomena, something we and others have previously shown (17, 49, 50).

While considered one of the classic strains, the NOD mice differed from the two primary clusters noted above. They displayed a combination of features from both groups and had a high degree of inter-islet variability, especially the female mice. NOD share common ancestors with the Swiss-Webster mice, which do not share parental origin with the other classic strains (47) and also display a “mixed” phenotype consisting of islet Ca^2+^ oscillations in response to glucose, with both slow and fast components (23).

Additionally, while all NOD mice were normoglycemic, a heterogeneous response was observed in islets from female, but not male, NOD mice. Female NOD mice are known to develop islet immune infiltration and subsequent autoimmune diabetes whereas males are largely protected from this (51). Male NOD islets were largely consistent in their Ca^2+^ waveforms. In the females, however, a high degree of heterogeneity in responses was observed across the female’s islets (**Supplemental Figure 2**). For any NOD female, some islets resembled those from NOD males in their clear oscillations, others largely lacked oscillatory behavior other than a strong pulse in response to 8G/QLA, and still others had an intermediate response. These observations may reflect varying degrees of dysfunction in the NOD female islets as the mice progress to diabetes, though we cannot say whether this results from variation in beta-cell intrinsic defects or islet immune infiltration.

The NZO mice also varied from the two clusters previously discussed. Male, and all but one of the female NZO mice were diabetic. Islets from diabetic mice had reduced amplitude and oscillatory behavior, other than a single pulse in 8G/QLA. This pattern is similar to the patterns observed in many of the NOD female islets. On the other hand, islets of the one non-diabetic female NZO mouse demonstrated clear, slow oscillations (**Supplemental Figure 1B**), which was surprising given reports of low K_ATP_ abundance due to *Abcc8* mutations in the NZO (52) and the strong role of K_ATP_ channels in regulating islet Ca^2+^(11, 32, 33, 53). While not of the same lineage as the NOD, the NZO do exhibit some autoimmune infiltration in the pancreas (54), and the marked difference between the non-diabetic and diabetic NZOs, along with the variation in female NOD islet responses, further suggests that intra-islet variability for the NOD mice may be the result of disease progression.

Understanding the genetic variation driving islet responses in the founders may be informative beyond these specific strains. Screens in the DO mice and similar outbred populations can track SNPs associated with trait variation to their parental inbred strain of origin. Our previous genetic screen for drivers of islet function observed that many of the quantitative trait loci (QTL) appearing for *ex vivo* islet traits had effects driven by the SNPs from the wild-derived strains as opposed to the classic inbred groups (16). For example, the QTL mediated by *Zfp148*, which drives Ca^2+^ oscillation and insulin secretion phenotypes in β-cells (41), also had strong strain effects from wild-derived strains (16).

Previous studies of islet Ca^2+^ have largely been confined to a handful of strains, and many studies by individual labs tend to use the strains linked with their projects. While this does include a few outbred stocks (e.g. NMRI (55–57), CD-1 (9, 58, 59)), direct comparisons of these to traditional inbred lines are rare (55), and studies of specific genes often use traditional inbred lines, as wild-derived lines do not respond as well to the conventional assistive reproductive technologies required for genome editing and transgenesis (60, 61).

One study, comparing the outbred NMRI stock to the C57BL6/J and C57BL/6N strains (55), found that the NMRI displayed significantly lower frequencies than the C57 lines, particularly in physiological glucose ranges and had similar active periods. While highly informative, there were important differences between these studies and our studies here. Of note, the studies were done in acute slice culture, in only one sex, and the frequencies detected did not resemble the (at least for the C57 lines) the slow oscillations observed in isolated islets from these inbred strains (e.g. **Figure 1**, (6, 41)).

### Mouse-to-human integration nominates novel islet drivers

One limitation of our current study is that the association between islet proteins and Ca^2+^ waveforms is correlative and therefore cannot distinguish proteins that are causal for the differences in islet Ca^2+^ between strains from proteins that change because of these differences. One approach to discriminate cause from effect, and establish the relevance to humans, is to identify whether genes encoding human orthologues of these proteins are associated with glycemic traits in human GWAS. SNPs for glycemic traits (**Table 2**), particularly those involving insulin, suggest that alterations in these proteins may impart disease risk, which is less likely for proteins that do not play critical regulatory roles. Thus, filtering our candidates for glycemic trait associations in human GWAS, while not definitive, suggests a likely causal role for these proteins in mediating differences in islet Ca^2+^ and insulin secretion among the different mice. Integrating human GWAS data with the proteins most correlated to Ca^2+^ dynamics nominated ∼650 protein candidates, of which approximately one-third have been previously shown to have roles in islet biology. These include well-established drivers of insulin secretion; e.g. SUR1, GLUT2, and GNAS. Other previously unknown candidates show promise for validation, as they are already targets of small molecule compounds (e.g. ACP1 and others, (62–67)), are secreted (e.g. COBLL1 and others, (68–75)), or have been knocked out in mice, resulting in metabolic phenotypes (**Figure 7B**, **Table 3**, **Table 4** (76)).

Our approach to merge human GWAS with our findings in mouse assumes that the glycemic-related SNPs we nominated alter the abundance or function of the human orthologues. Most SNPs that are strongly associated with phenotypes in human GWAS are noncoding, residing within introns, promoters, 3’UTRs, or intergenic regions (e.g. **Figure 6**). Therefore, a limitation of our approach is the assumption that SNPs regulate the gene they are proximal to, which is not always accurate (77–79). To infer a more direct link between SNPs and potential target genes, we incorporated human islet chromatin data (38). Physical contact between a region containing SNPs and a distal gene supports a regulatory role, as for ACP1 (**Figure 6B**). Additionally, SNPs within regions of open chromatin (ATAC-seq) and actively transcribed regions (histone markers) suggest a higher likelihood of regulating transcription factor access. While this approach does not conclusively show a link between the SNPs and expression of the orthologue for our candidate proteins, these chromatin data more strongly suggest that the orthologue expression may be regulated by the candidates’ SNPs.

### Exploiting strain and sex-dependent differences in Ca^2+^ dynamics for model system selection

In addition to the candidate regulators with potential relevance to human islet biology, we provide a user-friendly web interface to our data where users can determine whether their gene of interest has a potential regulatory role in islets. Multiple inferences regarding the roles of specific pathways are possible via analysis of Ca^2+^ oscillations in islets (6, 8, 9, 19), and our protein correlation data provides a resource to identify which parameter most closely correlates to a number of Ca^2+^ traits. Additionally, it highlights strain/sex outliers for a given trait or gene product, which can be used to select which strain/sex is best to explore that gene’s role (e.g. **Figure 7C**). Newer technologies in reproductive assistance, transgenesis, and gene editing, together with more accurate genome sequencing and single mutations conferring docility, are quickly making utilization of the wild-derived mice more practical (60, 61, 80–82). As many of the QTL identified in DO-based studies often have strong driver SNPs from the wild-derived strains, a further understanding of which experimental questions might be best addressed by use of these strains will be important.

We have previously provided user-friendly web interfaces that allow searches of gene expression as a function of diet (WD vs chow, (83, 84)) and background (e.g. BTBR and B6, (83, 84)), correlation and QTL scans in F2 intercrosses of these mice (85), and where these may align with QTL in our DO studies (16). Many of our candidates are strongly altered by diet and have strong correlations in the F2 data for certain clinical traits including insulin and glycemic parameters (85). Here, we provide the correlation data for islet proteins against multiple parameters describing islet Ca^2+^ responses between strains (DOI: https://doi.org/10.5061/dryad.j0zpc86jc, https://data-viz.it.wisc.edu/FounderCalciumStudy/, https://github.com/byandell/FounderCalciumStudy, https://rstudio.it.wisc.edu/FounderCalciumStudy). These will enable researchers to better identify proteins or parameters of interest as well as appropriate background strains with which to determine the functions of these proteins.

## MATERIALS & METHODS

### Chemicals

All general chemicals, amino acids, BSA, DMSO, glucose, glucose-dependent insulinotropic polypeptide (GIP, G2269), cOmplete Mini EDTA-free Protease Inhibitor Cocktail Tablets (11836170001), and heat-inactivated FBS (12306C) were purchased from Sigma Aldrich. RPMI 1640 base medium (11-875-093), antibiotic–antimycotic solutions (15240112), NP-40 Alternative (492016), Fura Red Ca^2+^ imaging dye (F3020), DiR (D12731), and agarose (BP1356-500) were purchased from ThermoFisher. Glass-bottomed culture dishes were ordered from Mattek (P35G-0-14-C). Fura Red stocks were prepared at 5 mM concentrations in DMSO, aliquoted into light-shielded tubes, and stored at –20°C until day of use (5 μM final concentration). DiR was prepared in DMSO at 2 mg/ mL, aliquoted to light-shielded tubes, and stored at 4°C until use. All imaging solutions were prepared in a bicarbonate/HEPES-buffered imaging medium (formula in **Table 1**). Amino acids were prepared as 100× stock in the biocarbonate/HEPES-buffered imaging medium, aliquoted into 1.5 mL tubes, and frozen at –20°C until day of use. Aliquots of GIP stock were prepared at 100 μM in water and kept at -20°C until day of use.

### Animals

Animal care and experimental protocols were approved by the University of Wisconsin-Madison Animal Care and Use Committee. Most strains (B6, AJ, 129, NOD, PWK, and WSB) were bred in-house, although two strains (CAST and NZO) were purchased from Jackson Laboratory (Bar Harbor, ME). All mice were fed a high-fat, high-sucrose Western-style diet (WD, consisting of 44.6% kcal fat, 34% carbohydrate, and 17.3% protein) from Envigo Teklad (TD.08811) beginning at 4 weeks and continuing until sacrifice (aged ∼19-20 weeks for all strains except the NZO males). The NZO males were sacrificed at 12 weeks of age owing to complications from severe diabetes. For each strain, 3-7 males and females from at least 2 litters were analyzed. Animals were sacrificed by cervical dislocation prior to islet isolation.

### In vivo measurements

Fasting blood glucose and insulin levels were measured in mice at 19 weeks of age, except for the NZO males which were measured at 12 weeks of age. Glucose was analyzed by the glucose oxidase method using a commercially available kit (TR15221, Thermo Fisher Scientific), and insulin was measured by radioimmunoassay (RIA; SRI-13K, Millipore). This is the same assay that was used to measure plasma insulin for the previously published cohort used for the correlation analysis in Figure 4 (17).

### Islet imaging

Islets were isolated as previously described (86) and incubated in recovery medium (RPMI 1640, 11.1 mM glucose, 1% antibiotic/antimycotic, 10% FBS) overnight at 37°C and 5% CO_2_. Islets were then incubated with Fura Red (5 μM in recovery medium) at 37°C for 45 minutes. Imaging dishes were created from glass-bottomed 10 cm^2^ dishes that had been filled with agarose. A channel with a central well was cut into the agarose with expanded ports on either side of the well for inflow and outflow lines. Prior to loading the chambers were perfused with the initial imaging solution (8 mM glucose in imaging medium). Islets were then loaded into these dishes. The imaging chamber was placed on a 37°C– heated microscope stage (Tokai Hit TIZ) of a Nikon A1R-Si+ confocal microscope. The solutions included 8 mM glucose (8G), 8 mM glucose + 2 mM glutamine, 0.5 mM leucine, and 1.5 mM alanine (8G/QLA), 8G/QLA + 10 nM glucose-dependent insulinotropic polypeptide (8G/QLA/GIP), and 2 mM glucose (2G), each of which were kept in a 37°C water bath. Solutions were perfused through the chamber at 0.25 mL/min for 40 minutes each, with constant flow controlled by a Fluigent MCFS-EZ and M-switch valve assembly (Fluigent). The scope was integrated with a Nikon Eclipse-Ti Inverted scope and equipped with a Nikon CFI Apochromat Lambda D 10x/0.45 objective (Nikon Instruments), fluorescence spectral detector, and multiple laser lines (Nikon LU-NV laser unit; 405, 440, 488, 514, 561, 640nm). Bound dye was excited with the 405nm laser and the spectral detector’s variable filter was set to 620-690nm. The free dye was excited with the 488nm laser and the variable filter collected from 640-690nm. Images were collected at 1 frame/sec at 6 second intervals. Each islet was considered a region of interest for further analysis. ROI intensity was collected by NIS Elements and exported for further analysis. All microscopy was performed at the University of Wisconsin-Madison Biochemistry Optical Core, which was established with support from the University of Wisconsin-Madison Department of Biochemistry Endowment.

### Islet perifusion

Isolated islets were kept in RPMI-based medium (see above) overnight prior to perifusion, which was performed as previously described, with minor modifications (41, 87). Islets were equilibrated in 2 mM glucose for 55 minutes, after which 100 μL fractions were collected every minute with the perifusion solutions set at a flow rate of 100 μL/min. All solutions and islet chambers were kept at 37°C. After the final fraction was collected, islet chambers were disconnected, inverted, and flushed with 2 mL of NP-40 Alternative lysis buffer containing protease inhibitors for islet insulin extraction.

### Secreted insulin assay

Insulin in each perifusion fraction and islet insulin content were determined using a custom assay, as previously described (17).

### Imaging data analysis

Trace segments for each solution condition were analyzed using Matlab and R. Traces were detrended using custom R scripts and Graphpad PRISM. Custom Matlab scripts (7) (https://github.com/hrfoster/Merrins-Lab-Matlab-Scripts, also stored on Dryad https://doi.org/10.5281/zenodo.6540721) determined oscillation peak amplitude, pulse duration, active duration (the time when Ca^2+^ is above 50% peak amplitude), silent duration (the difference between period and active duration), plateau fraction (the fraction of overall time per pulse spent in the active duration), pulse period and other parameters. Spectral density deconvolution for the trace segments to determine principal frequencies was done using R. Animal averages for the different parameters defined by Matlab and R were computed and graphed using custom R scripts. Figures were created using CorelDraw and Biorender.com. All R scripts and the citations for the relevant packages used to generate them are available via Dryad (https://doi.org/10.5061/dryad.j0zpc86jc).

### Correlation and Z-score calculations

Correlation analysis was performed using the imaging data measurements and our published islet protein abundance data, *ex vivo* static insulin secretion measurements, and *in vivo* measurements made in a separate cohort of mice on the WD from the same strains and sexes used in these studies (17). For each imaging parameter or previously published measurement, the Z-score was calculated using the formula *z = (x-μ) / σ* where *z* is the Z-score, *x* is the animal average for that trait given the strain and sex, *μ* is the average of all animals’ values for that trait, and *σ* is the standard deviation for all animals’ values for that trait. Z-scores were computed in R and excel for the imaging parameters and the previously published (17) islet proteomic, *ex vivo* secretion, and *in vivo* measurements.

Correlation coefficients between the Z-score values of the imaging parameters and Z-scores of the previously published protein abundance, islet secretion, and *in vivo* traits were computed in Excel using the CORREL function. The equation used for this function is:

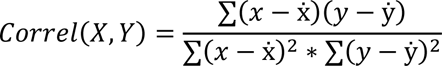

Where *X* and *Y* are the Z-scores for the correlated traits/parameters, *ẋ* is the population average for trait X and ẏ is the population average for trait Y. Traits were considered highly correlated if absolute value for their Z-score correlation coefficients was ≥ 0.5.

### Gene enrichment and human GWAS analysis

Proteins highly correlated or anticorrelated to imaging parameters were further analyzed using pathway enrichment and presence of human GWAS SNPs. Briefly, for a given parameter, pathway analysis for the highly correlated or anti-correlated proteins to that parameter was done using Enrichr (37, 88). Enrichr links for the subsets of proteins highly correlated to specific calcium parameters are provided in **Supplemental Table 1**, which is stored on Dryad (https://doi.org/10.5061/dryad.j0zpc86jc).

For GWAS analysis, human orthologues for genes encoding the previously measured islet proteins were identified using BioMart (89). For highly correlated proteins, the protein was deemed of human interest if its orthologue had SNPs for glycemia-related traits (see table 2) either along the gene body, within +/- 100 kbp of the gene start or end, or if any region in the gene body was connected to regions with SNPs by chromatin looping. SNPs were queried using Lunaris tool of the Common Metabolic Diseases Knowledge Portal (cmdkp.org). Chromatin loop anchor points for the relevant gene orthologues were identified using previously published human islet promoter-capture HiC data (38) and the alignment between these anchor loops and orthologues of interest was done using R scripts.

For those proteins having orthologues with SNPs via this analysis, we conducted further literature searches using Pubmed, Google Scholar, ChEMBL (64, 65, 90), canSAR (63), Uniprot (68), Tabula Muris (91), and the Human Protein Atlas (69, 70), and other resources (71, 92, 93) to determine tissue expression and identify any prior roles in islet biology. Figures for the relevant protein examples were created using GraphPad Prism, CorelDraw, and the WashU Epigenome Browser (94).

We further narrowed this list by searching each of the genes and their aliases in the PubMed, Google Scholar, and Google Search Engines along with “insulin secretion.” This allowed us to identify which genes have a known role in altering the insulin secretory pathway, and which genes may be understudied (**Figure 7 A**).

### Web resource

A web resource was created to explore the islet calcium and proteomic data and their relationships (https://data-viz.it.wisc.edu/FounderCalciumStudy, https://rstudio.it.wisc.edu/FounderCalciumStudy). This resource sits on an RStudio/Connect server (see https://posit.co/). It enables the user to select traits from the calcium and protein datasets to plot by strain, sex, and calcium parameters. Distinct mice were assayed for **calcium** and **protein**. Individual strains can be selected on the main menu using the checkboxes, or all strains (default) can be viewed.

The different datasets available in the main menu are:

1. **calcium**: calcium parameters & spectral density data, with stimulatory secretion conditions
2. **protein**: islet proteomic measurements
3. **basal**: average calcium in 2mM glucose

The **calcium** data have three stimulatory conditions (8G, 8G/QLA, and 8G/QLA/GIP), which are displayed together for each calcium parameter. The proteomic data (**protein**) are displayed for each identified peptide. In rare cases of multiple peptides per gene, both gene symbol and peptide identifier (PP number) are included (e.g. Pkm_PP_1521 for the M1 isoform of the protein PKM). Desired proteins can be selected simultaneously with desired calcium parameters for correlation analysis and paired display by both datasets. The **basal** elements retained from the **calcium** data include the Average Calcium measurement for 2mM glucose. Proteomic data were log10-transformed. All traits were transformed into normal scores, keeping the sample mean and variance the same.

Scatter plots display data across sex and calcium conditions. When plotting calcium against protein or basal traits, means by strain and sex are used, as the two experiments used different mice. Correlation of selected traits with all other traits in the resource use Pearson correlation on pairwise-complete data. The user can order traits by their significance or by their correlation to other selected traits.

Statistical modeling terms include strain, sex, and the strain:sex interaction, plus additional terms for comparing calcium condition with respect to strain and sex. Users can view volcano plots displaying deviation of term effects, measured as the standard deviation (SD = square root of mean square error) divided by the raw SD for that trait, against their significance (p-value after adjusting for all other model terms, presented on -log10 scale). In addition to the terms, a composite “signal” captures the combined effect of terms strain:condition + strain:condition:sex using a general F test computation.

All data handling and web app construction for the resource was performed using R scripts in publicly available GitHub repositories, with specifics for the calcium study at https://github.com/byandell/FounderCalciumStudy and the general purpose analysis and web deployment package at https://github.com/byandell/foundr.

### Statistics

For the islet perifusion insulin measurements, statistics were determined in GraphPad Prism. Fractional secretion area-under-the-curve (AUC) was determined using Prism and differences in AUCs analyzed using post-tests following 2-way ANOVA for the indicated trace segments. Islet total insulins between strains were compared using a two-tailed Student’s t-test with Welch’s correction. For **Supplemental Figure 5**, the graphs, Pearson’s R, and regression lines were created in Prism.

## Data Availability

All R scripts and raw data are available via Dryad (https://doi.org/10.5061/dryad.j0zpc86jc). Code for the web resource is available via Github (https://github.com/byandell/FounderCalciumStudy). Image files are available upon request.

## Study approval

All protocols were approved by the University of Wisconsin-Madison IACUC (Protocol A005821-R01)

## Author contributions

C. Emfinger, L. Clark, M. Merrins, M. Keller, and A. Attie designed experiments. C. Emfinger, L. Clark, K. Schueler, S. Simonett, D. Stapleton, and K. Mitok performed experiments. C. Emfinger, L. Clark, and B. Yandell wrote the paper. C. Emfinger, L. Clark, B. Yandell, and M. Keller performed data analysis. B. Yandell constructed the shiny app interface for the resource and incorporated the relevant data. C. Emfinger, L. Clark, M. Merrins, B. Yandell, M. Keller, and A. Attie edited the paper.

## Funding sources

This work was supported by NIH R01DK101573, R01DK102948, and RC2DK125961 to A. Attie; NIH R01DK113103, NIH R01DK127637, and VA I01B005113 to M. Merrins, ADA 7-21-PDF-157 to C. Emfinger, and by the University of Wisconsin–Madison, Department of Biochemistry and Office of the Vice Chancellor for Research and Graduate Education with funding from the Wisconsin Alumni Research Foundation to M. Keller.

Portions of the figures were created with Biorender.com. License files are PC24P29AVO, UA24P29B1E, TK24P29B44, QM24P29H2P, WF24P29HB3, OM24P2BPIL, and BN24P2C01S. We would also like to thank Dr. Michael Lloyd at The Jackson Laboratory for his help with setting up the Lunaris queries.

## ABBREVIATIONS

COBLL1: Cordon-Bleu-like Like 1
ACP1: Acid phosphatase 1
GALNS: galactosamine (N-acetyl)-6-sulfatase
SUR1: Sulfonylurea receptor 1
GNAS: Guanine nucleotide binding protein, alpha stimulating
GLUT2: Glucose transporter 2
GK: Glucokinase
PKM: Pyruvate Kinase M
GWAS: Genome-wide association study
SNP: single-nucleotide polymorphism

## REFERENCES

1. Dimas AS, Lagou V, Barker A, Knowles JW, Mägi R, Hivert MF, et al. Impact of type 2 diabetes susceptibility variants on quantitative glycemic traits reveals mechanistic heterogeneity. Diabetes. 2014;63(6):2158–71.

2. Wood AR, Jonsson A, Jackson AU, Wang N, van Leewen N, Palmer ND, et al. A Genome-Wide Association Study of IVGTT-Based Measures of First-Phase Insulin Secretion Refines the Underlying Physiology of Type 2 Diabetes Variants. Diabetes. 2017;66(8):2296–309.

3. Clee SM, and Attie AD. The genetic landscape of type 2 diabetes in mice. Endocr Rev. 2007;28(1):48–83.

4. Kebede MA, and Attie AD. Insights into obesity and diabetes at the intersection of mouse and human genetics. Trends in endocrinology and metabolism: TEM. 2014;25(10):493–501.

5. Sittig LJ, Carbonetto P, Engel KA, Krauss KS, Barrios-Camacho CM, and Palmer AA. Genetic Background Limits Generalizability of Genotype-Phenotype Relationships. Neuron. 2016;91(6):1253–9.

6. Lewandowski SL, Cardone RL, Foster HR, Ho T, Potapenko E, Poudel C, et al. Pyruvate Kinase Controls Signal Strength in the Insulin Secretory Pathway. Cell metabolism. 2020;32(5):736–50.e5.

7. Foster HR, Ho T, Potapenko E, Sdao SM, Huang SM, Lewandowski SL, et al. β-cell deletion of the PKm1 and PKm2 isoforms of pyruvate kinase in mice reveals their essential role as nutrient sensors for the K(ATP) channel. Elife. 2022;11.

8. Merrins MJ, Corkey BE, Kibbey RG, and Prentki M. Metabolic cycles and signals for insulin secretion. Cell metabolism. 2022;34(7):947–68.

9. Dahlgren GM, Kauri LM, and Kennedy RT. Substrate effects on oscillations in metabolism, calcium and secretion in single mouse islets of Langerhans. Biochimica et biophysica acta. 2005;1724(1-2):23–36.

10. Krippeit-Drews P, Düfer M, and Drews G. Parallel oscillations of intracellular calcium activity and mitochondrial membrane potential in mouse pancreatic B-cells. Biochemical and biophysical research communications. 2000;267(1):179–83.

11. Marinelli I, Thompson BM, Parekh VS, Fletcher PA, Gerardo-Giorda L, Sherman AS, et al. Oscillations in K(ATP) conductance drive slow calcium oscillations in pancreatic β-cells. Biophys J. 2022;121(8):1449–64.

12. Henquin JC. Regulation of insulin secretion: a matter of phase control and amplitude modulation. Diabetologia. 2009;52(5):739–51.

13. Chen C, Chmelova H, Cohrs CM, Chouinard JA, Jahn SR, Stertmann J, et al. Alterations in β-Cell Calcium Dynamics and Efficacy Outweigh Islet Mass Adaptation in Compensation of Insulin Resistance and Prediabetes Onset. Diabetes. 2016;65(9):2676–85.

14. Threadgill DW, Miller DR, Churchill GA, and de Villena FP. The collaborative cross: a recombinant inbred mouse population for the systems genetic era. Ilar J. 2011;52(1):24–31.

15. Svenson KL, Gatti DM, Valdar W, Welsh CE, Cheng R, Chesler EJ, et al. High-resolution genetic mapping using the Mouse Diversity outbred population. Genetics. 2012;190(2):437–47.

16. Keller MP, Rabaglia ME, Schueler KL, Stapleton DS, Gatti DM, Vincent M, et al. Gene loci associated with insulin secretion in islets from nondiabetic mice. The Journal of Clinical Investigation. 2019;129(10):4419–32.

17. Mitok KA, Freiberger EC, Schueler KL, Rabaglia ME, Stapleton DS, Kwiecien NW, et al. Islet proteomics reveals genetic variation in dopamine production resulting in altered insulin secretion. The Journal of biological chemistry. 2018;293(16):5860–77.

18. Nunemaker CS, Wasserman DH, McGuinness OP, Sweet IR, Teague JC, and Satin LS. Insulin secretion in the conscious mouse is biphasic and pulsatile. American Journal of Physiology-Endocrinology and Metabolism. 2006;290(3):E523–E9.

19. Kennedy RT, Kauri LM, Dahlgren GM, and Jung SK. Metabolic oscillations in beta-cells. Diabetes. 2002;51 Suppl 1:S152–61.

20. Lang DA, Matthews DR, Peto J, and Turner RC. Cyclic oscillations of basal plasma glucose and insulin concentrations in human beings. N Engl J Med. 1979;301(19):1023–7.

21. Colsoul B, Schraenen A, Lemaire K, Quintens R, Van Lommel L, Segal A, et al. Loss of high-frequency glucose-induced Ca2+ oscillations in pancreatic islets correlates with impaired glucose tolerance in Trpm5-/- mice. Proceedings of the National Academy of Sciences of the United States of America. 2010;107(11):5208–13.

22. Corbin KL, Waters CD, Shaffer BK, Verrilli GM, and Nunemaker CS. Islet Hypersensitivity to Glucose Is Associated With Disrupted Oscillations and Increased Impact of Proinflammatory Cytokines in Islets From Diabetes-Prone Male Mice. Endocrinology. 2016;157(5):1826–38.

23. Nunemaker CS, Bertram R, Sherman A, Tsaneva-Atanasova K, Daniel CR, and Satin LS. Glucose modulates [Ca2+]i oscillations in pancreatic islets via ionic and glycolytic mechanisms. Biophys J. 2006;91(6):2082–96.

24. Marinelli I, Parekh V, Fletcher P, Thompson B, Ren J, Tang X, et al. Slow oscillations persist in pancreatic beta cells lacking phosphofructokinase M. Biophys J. 2022;121(5):692–704.

25. Nunemaker CS, Zhang M, Wasserman DH, McGuinness OP, Powers AC, Bertram R, et al. Individual mice can be distinguished by the period of their islet calcium oscillations: is there an intrinsic islet period that is imprinted in vivo? Diabetes. 2005;54(12):3517–22.

26. Bertram R, Satin LS, and Sherman AS. Closing in on the Mechanisms of Pulsatile Insulin Secretion. Diabetes. 2018;67(3):351–9.

27. Bertram R, Sherman A, and Satin LS. Metabolic and electrical oscillations: partners in controlling pulsatile insulin secretion. American journal of physiology Endocrinology and metabolism. 2007;293(4):E890–900.

28. Whitticar NB, and Nunemaker CS. Reducing Glucokinase Activity to Enhance Insulin Secretion: A Counterintuitive Theory to Preserve Cellular Function and Glucose Homeostasis. Frontiers in endocrinology. 2020;11:378-.

29. Koneshamoorthy A, Seniveratne-Epa D, Calder G, Sawyer M, Kay TWH, Farrell S, et al. Case Report: Hypoglycemia Due to a Novel Activating Glucokinase Variant in an Adult - a Molecular Approach. Front Endocrinol (Lausanne). 2022;13:842937.

30. Tengholm A, and Gylfe E. cAMP signalling in insulin and glucagon secretion. *Diabetes*, Obesity and Metabolism. 2017;19(S1):42–53.

31. Shuai H, Xu Y, Ahooghalandari P, and Tengholm A. Glucose-induced cAMP elevation in β-cells involves amplification of constitutive and glucagon-activated GLP-1 receptor signalling. *Acta physiologica (Oxford*, England). 2021;231(4):e13611.

32. Koster JC, Marshall BA, Ensor N, Corbett JA, and Nichols CG. Targeted Overactivity of β Cell KATP Channels Induces Profound Neonatal Diabetes. Cell. 2000;100(6):645–54.

33. Remedi MS, and Nichols CG. In: Pitt GS ed. Ion Channels in Health and Disease. Boston: Academic Press; 2016:199–221.

34. Jennings RE, Scharfmann R, and Staels W. Transcription factors that shape the mammalian pancreas. Diabetologia. 2020;63(10):1974–80.

35. Rutter GA, Georgiadou E, Martinez-Sanchez A, and Pullen TJ. Metabolic and functional specialisations of the pancreatic beta cell: gene disallowance, mitochondrial metabolism and intercellular connectivity. Diabetologia. 2020;63(10):1990–8.

36. Stijnen P, Ramos-Molina B, O’Rahilly S, and Creemers JW. PCSK1 Mutations and Human Endocrinopathies: From Obesity to Gastrointestinal Disorders. Endocr Rev. 2016;37(4):347–71.

37. Chen EY, Tan CM, Kou Y, Duan Q, Wang Z, Meirelles GV, et al. Enrichr: interactive and collaborative HTML5 gene list enrichment analysis tool. BMC Bioinformatics. 2013;14:128.

38. Miguel-Escalada I, Bonas-Guarch S, Cebola I, Ponsa-Cobas J, Mendieta-Esteban J, Atla G, et al. Human pancreatic islet three-dimensional chromatin architecture provides insights into the genetics of type 2 diabetes. Nature genetics. 2019.

39. Bergman RN, Zaccaro DJ, Watanabe RM, Haffner SM, Saad MF, Norris JM, et al. Minimal model-based insulin sensitivity has greater heritability and a different genetic basis than homeostasis model assessment or fasting insulin. Diabetes. 2003;52(8):2168–74.

40. Carter JD, Dula SB, Corbin KL, Wu R, and Nunemaker CS. A practical guide to rodent islet isolation and assessment. Biol Proced Online. 2009;11:3–31.

41. Emfinger CH, de Klerk E, Schueler KL, Rabaglia ME, Stapleton DS, Simonett SP, et al. β Cell– specific deletion of Zfp148 improves nutrient-stimulated β cell Ca2+ responses. JCI Insight. 2022;7(10).

42. El K, Gray SM, Capozzi ME, Knuth ER, Jin E, Svendsen B, et al. GIP mediates the incretin effect and glucose tolerance by dual actions on α cells and β cells. Sci Adv. 2021;7(11).

43. Capozzi ME, Svendsen B, Encisco SE, Lewandowski SL, Martin MD, Lin H, et al. β Cell tone is defined by proglucagon peptides through cAMP signaling. JCI Insight. 2019;4(5).

44. Kang G, Leech CA, Chepurny OG, Coetzee WA, and Holz GG. Role of the cAMP sensor Epac as a determinant of KATP channel ATP sensitivity in human pancreatic beta-cells and rat INS-1 cells. J Physiol. 2008;586(5):1307–19.

45. Holz GGt, Kühtreiber WM, and Habener JF. Pancreatic beta-cells are rendered glucose-competent by the insulinotropic hormone glucagon-like peptide-1(7-37). Nature. 1993;361(6410):362–5.

46. Heart E, Corkey RF, Wikstrom JD, Shirihai OS, and Corkey BE. Glucose-dependent increase in mitochondrial membrane potential, but not cytoplasmic calcium, correlates with insulin secretion in single islet cells. American Journal of Physiology-Endocrinology and Metabolism. 2006;290(1):E143–E8.

47. Beck JA, Lloyd S, Hafezparast M, Lennon-Pierce M, Eppig JT, Festing MFW, et al. Genealogies of mouse inbred strains. Nature genetics. 2000;24(1):23–5.

48. Yang H, Wang JR, Didion JP, Buus RJ, Bell TA, Welsh CE, et al. Subspecific origin and haplotype diversity in the laboratory mouse. Nature genetics. 2011;43(7):648–55.

49. Lee KT, Karunakaran S, Ho MM, and Clee SM. PWD/PhJ and WSB/EiJ mice are resistant to diet-induced obesity but have abnormal insulin secretion. Endocrinology. 2011;152(8):3005–17.

50. Kreznar JH, Keller MP, Traeger LL, Rabaglia ME, Schueler KL, Stapleton DS, et al. Host Genotype and Gut Microbiome Modulate Insulin Secretion and Diet-Induced Metabolic Phenotypes. Cell Reports. 2017;18(7):1739–50.

51. Pearson JA, Wong FS, and Wen L. The importance of the Non Obese Diabetic (NOD) mouse model in autoimmune diabetes. J Autoimmun. 2016;66:76–88.

52. Andrikopoulos S, Fam BC, Holdsworth A, Visinoni S, Ruan Z, Stathopoulos M, et al. Identification of ABCC8 as a contributory gene to impaired early-phase insulin secretion in NZO mice. The Journal of endocrinology. 2016;228(1):61–73.

53. Ashcroft FM, Puljung MC, and Vedovato N. Neonatal Diabetes and the K ATP Channel: From Mutation to Therapy. Trends in Endocrinology & Metabolism. 2017;28(5):377–87.

54. Junger E, Herberg L, Jeruschke K, and Leiter EH. The Diabetes-Prone NZO/Hl Strain. II. Pancreatic Immunopathology. Laboratory Investigation. 2002;82(7):843–53.

55. Pohorec V, Križančić Bombek L, Skelin Klemen M, Dolenšek J, and Stožer A. Glucose-Stimulated Calcium Dynamics in Beta Cells From Male C57BL/6J, C57BL/6N, and NMRI Mice: A Comparison of Activation, Activity, and Deactivation Properties in Tissue Slices. Front Endocrinol (Lausanne). 2022;13:867663.

56. Šterk M, Križančić Bombek L, Skelin Klemen M, Slak Rupnik M, Marhl M, Stožer A, et al. NMDA receptor inhibition increases, synchronizes, and stabilizes the collective pancreatic beta cell activity: Insights through multilayer network analysis. PLoS computational biology. 2021;17(5):e1009002.

57. Stožer A, Skelin Klemen M, Gosak M, Križančić Bombek L, Pohorec V, Slak Rupnik M, et al. Glucose-dependent activation, activity, and deactivation of beta cell networks in acute mouse pancreas tissue slices. American journal of physiology Endocrinology and metabolism. 2021;321(2):E305–e23.

58. Hauke S, Rada J, Tihanyi G, Schilling D, and Schultz C. ATP is an essential autocrine factor for pancreatic β-cell signaling and insulin secretion. Physiol Rep. 2022;10(1):e15159.

59. Scarl RT, Corbin KL, Vann NW, Smith HM, Satin LS, Sherman A, et al. Intact pancreatic islets and dispersed beta-cells both generate intracellular calcium oscillations but differ in their responsiveness to glucose. Cell Calcium. 2019;83:102081.

60. Hirose M, Hasegawa A, Mochida K, Matoba S, Hatanaka Y, Inoue K, et al. CRISPR/Cas9- mediated genome editing in wild-derived mice: generation of tamed wild-derived strains by mutation of the a (nonagouti) gene. Scientific Reports. 2017;7(1):42476.

61. Mochida K, Hasegawa A, Otaka N, Hama D, Furuya T, Yamaguchi M, et al. Devising Assisted Reproductive Technologies for Wild-Derived Strains of Mice: 37 Strains from Five Subspecies of Mus musculus. PloS one. 2014;9(12):e114305.

62. Stanford SM, Diaz MA, Ardecky RJ, Zou J, Roosild T, Holmes ZJ, et al. Discovery of Orally Bioavailable Purine-Based Inhibitors of the Low-Molecular-Weight Protein Tyrosine Phosphatase. J Med Chem. 2021;64(9):5645–53.

63. Coker EA, Mitsopoulos C, Tym JE, Komianou A, Kannas C, Di Micco P, et al. canSAR: update to the cancer translational research and drug discovery knowledgebase. Nucleic acids research. 2019;47(D1):D917–D22.

64. Davies M, Nowotka M, Papadatos G, Dedman N, Gaulton A, Atkinson F, et al. ChEMBL web services: streamlining access to drug discovery data and utilities. Nucleic acids research. 2015;43(W1):W612–W20.

65. Gaulton A, Hersey A, Nowotka M, Bento AP, Chambers J, Mendez D, et al. The ChEMBL database in 2017. Nucleic acids research. 2017;45(D1):D945–D54.

66. Santos R, Ursu O, Gaulton A, Bento AP, Donadi RS, Bologa CG, et al. A comprehensive map of molecular drug targets. Nat Rev Drug Discov. 2017;16(1):19–34.

67. Zhou Y, Zhang Y, Lian X, Li F, Wang C, Zhu F, et al. Therapeutic target database update 2022: facilitating drug discovery with enriched comparative data of targeted agents. Nucleic acids research. 2022;50(D1):D1398–D407.

68. The UniProt C. UniProt: the universal protein knowledgebase in 2021. Nucleic acids research. 2021;49(D1):D480–D9.

69. Thul PJ, Åkesson L, Wiking M, Mahdessian D, Geladaki A, Ait Blal H, et al. A subcellular map of the human proteome. Science (New York, NY). 2017;356(6340).

70. Uhlén M, Fagerberg L, Hallström BM, Lindskog C, Oksvold P, Mardinoglu A, et al. Proteomics. Tissue-based map of the human proteome. Science (New York, NY). 2015;347(6220):1260419.

71. Uhlén M, Karlsson MJ, Hober A, Svensson AS, Scheffel J, Kotol D, et al. The human secretome. Science signaling. 2019;12(609).

72. Navajas R, Ramos-Fernandez A, Herraiz I, Galindo A, Bartha JL, Corrales F, et al. Quantitative proteomic analysis of serum-purified exosomes identifies putative pre-eclampsia-associated biomarkers. Clin Proteomics. 2022;19(1):5.

73. Wang F, Xu Z, Zhou J, Lo W-S, Lau C-F, Nangle LA, et al. Regulated capture by exosomes of mRNAs for cytoplasmic tRNA synthetases. The Journal of biological chemistry. 2013;288(41):29223–8.

74. Chen G, Chen J, Liu H, Chen S, Zhang Y, Li P, et al. Comprehensive Identification and Characterization of Human Secretome Based on Integrative Proteomic and Transcriptomic Data. Frontiers in Cell and Developmental Biology. 2019;7.

75. Gonzales PA, Pisitkun T, Hoffert JD, Tchapyjnikov D, Star RA, Kleta R, et al. Large-Scale Proteomics and Phosphoproteomics of Urinary Exosomes. Journal of the American Society of Nephrology. 2009;20(2).

76. Groza T, Gomez FL, Mashhadi HH, Muñoz-Fuentes V, Gunes O, Wilson R, et al. The International Mouse Phenotyping Consortium: comprehensive knockout phenotyping underpinning the study of human disease. Nucleic acids research. 2023;51(D1):D1038–D45.

77. Nyaga DM, Vickers MH, Jefferies C, Perry JK, and O’Sullivan JM. Type 1 Diabetes Mellitus-Associated Genetic Variants Contribute to Overlapping Immune Regulatory Networks. Frontiers in genetics. 2018;9:535.

78. Chen X-F, Guo M-R, Duan Y-Y, Jiang F, Wu H, Dong S-S, et al. Multiomics dissection of molecular regulatory mechanisms underlying autoimmune-associated noncoding SNPs. JCI Insight. 2020;5(17).

79. Smemo S, Tena JJ, Kim KH, Gamazon ER, Sakabe NJ, Gómez-Marín C, et al. Obesity-associated variants within FTO form long-range functional connections with IRX3. Nature. 2014;507(7492):371–5.

80. Karunakaran S, and Clee SM. Genetics of metabolic syndrome: potential clues from wild-derived inbred mouse strains. Physiol Genomics. 2018;50(1):35–51.

81. Chao T, Liu Z, Zhang Y, Zhang L, Huang R, He L, et al. Precise and Rapid Validation of Candidate Gene by Allele Specific Knockout With CRISPR/Cas9 in Wild Mice. Frontiers in genetics. 2019;10:124.

82. Chang PL, Kopania E, Keeble S, Sarver BAJ, Larson E, Orth A, et al. Whole exome sequencing of wild-derived inbred strains of mice improves power to link phenotype and genotype. Mammalian genome : official journal of the International Mammalian Genome Society. 2017;28(9-10):416–25.

83. Keller MP, Choi Y, Wang P, Davis DB, Rabaglia ME, Oler AT, et al. A gene expression network model of type 2 diabetes links cell cycle regulation in islets with diabetes susceptibility. Genome research. 2008;18(5):706–16.

84. Yau B, Naghiloo S, Diaz-Vegas A, Carr AV, Van Gerwen J, Needham EJ, et al. Proteomic pathways to metabolic disease and type 2 diabetes in the pancreatic islet. iScience. 2021;24(10):103099.

85. Lan H, Chen M, Flowers JB, Yandell BS, Stapleton DS, Mata CM, et al. Combined Expression Trait Correlations and Expression Quantitative Trait Locus Mapping. PLOS Genetics. 2006;2(1):e6.

86. Rabaglia ME, Gray-Keller MP, Frey BL, Shortreed MR, Smith LM, and Attie AD. α-Ketoisocaproate-induced hypersecretion of insulin by islets from diabetes-susceptible mice. American Journal of Physiology-Endocrinology and Metabolism. 2005;289(2):E218–E24.

87. Bhatnagar S, Oler AT, Rabaglia ME, Stapleton DS, Schueler KL, Truchan NA, et al. Positional cloning of a type 2 diabetes quantitative trait locus; tomosyn-2, a negative regulator of insulin secretion. PLoS Genet. 2011;7(10):e1002323.

88. Kuleshov MV, Jones MR, Rouillard AD, Fernandez NF, Duan Q, Wang Z, et al. Enrichr: a comprehensive gene set enrichment analysis web server 2016 update. Nucleic acids research. 2016;44(W1):W90–7.

89. Smedley D, Haider S, Ballester B, Holland R, London D, Thorisson G, et al. BioMart--biological queries made easy. BMC Genomics. 2009;10:22.

90. Jupp S, Malone J, Bolleman J, Brandizi M, Davies M, Garcia L, et al. The EBI RDF platform: linked open data for the life sciences. Bioinformatics. 2014;30(9):1338–9.

91. Schaum N, Karkanias J, Neff NF, May AP, Quake SR, Wyss-Coray T, et al. Single-cell transcriptomics of 20 mouse organs creates a Tabula Muris. Nature. 2018;562(7727):367–72.

92. Varshney A, Scott LJ, Welch RP, Erdos MR, Chines PS, Narisu N, et al. Genetic regulatory signatures underlying islet gene expression and type 2 diabetes. Proceedings of the National Academy of Sciences of the United States of America. 2017;114(9):2301–6.

93. Lawlor N, George J, Bolisetty M, Kursawe R, Sun L, Sivakamasundari V, et al. Single-cell transcriptomes identify human islet cell signatures and reveal cell-type-specific expression changes in type 2 diabetes. Genome research. 2017;27(2)y:208–22.

94. Li D, Hsu S, Purushotham D, Sears RL, and Wang T. WashU Epigenome Browser update 2019. Nucleic acids research. 2019;47(W1):W158–W65.

95. Blake JA, Baldarelli R, Kadin JA, Richardson JE, Smith CL, and Bult CJ. Mouse Genome Database (MGD): Knowledgebase for mouse-human comparative biology. Nucleic acids research. 2021;49(D1):D981–d7.

