## Supplemental Figures for "Novel regulators of islet function identified from genetic variation in mouse islet Ca^2+^ oscillations"

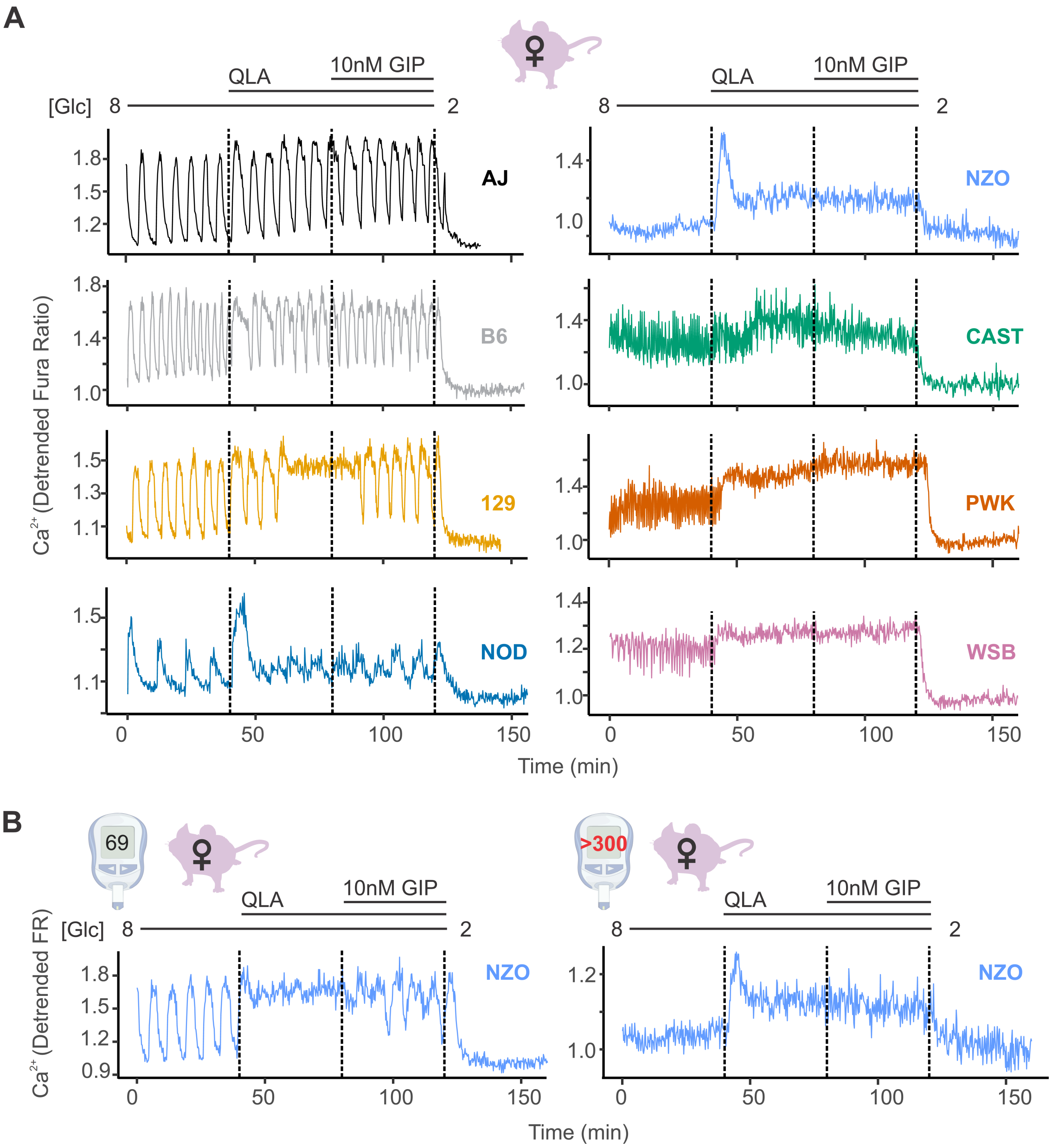


**Supplemental Figure 1. The high diversity in Ca$^{2+}$ oscillation in males is also observed in female mice. (A)** Representative Ca$^{2+}$ traces for female mice (n = 3-7 mice per strain, and 11-94 islets per mouse) exhibit a high degree of variability across the eight strains. Abbreviations: '[Glu]' = 'concentration of glucose in mM' (**B**) Representative Ca$^{2+}$ trace for non-diabetic NZO female (left panel, fasting plasma glucose 69 mg/dL) and a diabetic NZO female (right trace, fasting plasma glucose > 300 mg/dL). Related to Figure 1.

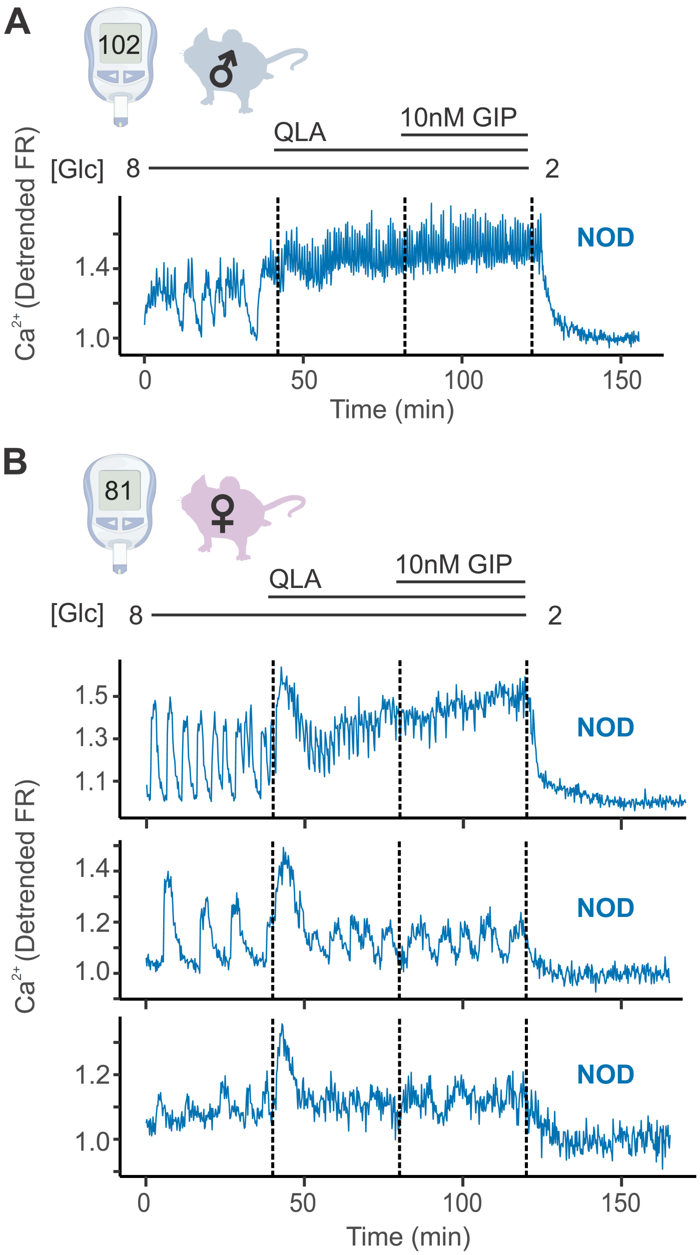


**Supplemental Figure 2. Diverse responses in non-diabetic NOD females' islets. (A)** Representative $Ca^{2+}$ traces for a male NOD mouse, for which islets closely resemble the trace pattern shown. (**B**) Example traces from islets of a single non-diabetic NOD female mouse. The pattern observed, where some of the mouse's islets appeared similar to the NOD male islets (top panel), some appeared similar to the diabetic NZO islets (bottom panel), and some presented with an intermediate phenotype (oscillations present in all stimulatory conditions, but with a more pronounced 8/QLA initial peak and diminished amplitudes; middle panel), was consistently observed for all the NOD females. Abbreviations: '[Glu]' = 'concentration of glucose in mM'. Related to Figure 1.

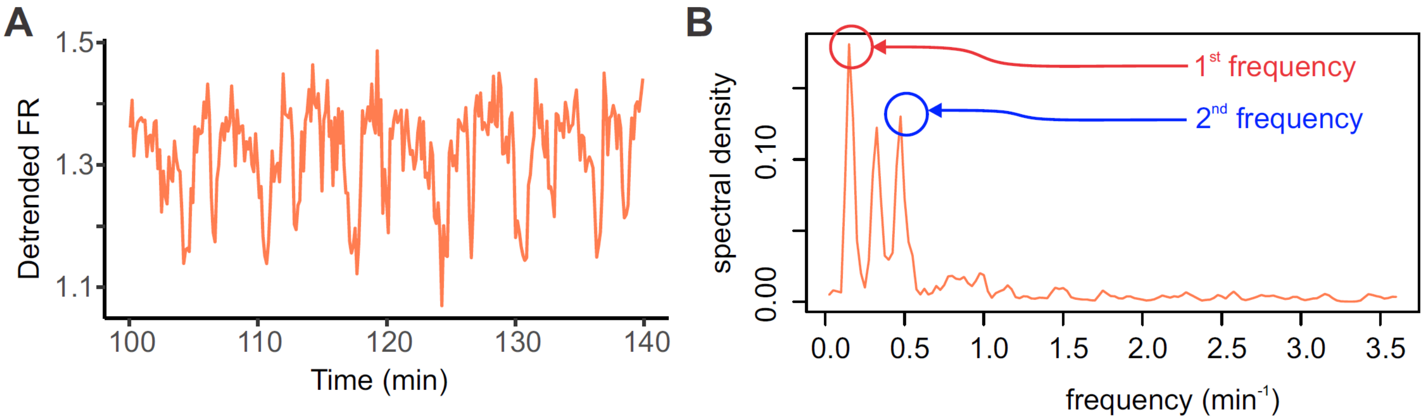


**Supplemental Figure 3. Example of spectral density breakdown for $Ca^{2+}$ traces. (A)** For the trace segment shown, the wave can be broken down into individual periodic waves (e.g. sine waves) of discrete amplitudes that, when added together, reproduce the trace. The frequency of each of these waves and its relative contribution to the overall trace signal is shown in (**B**), with frequency on the x axis and the spectral density, or the strength of each component signal, on the Y axis. This was computed using scripts in R. The strongest 2 frequencies, denoted 1$^{st}$and 2$^{nd}$ respectively, are indicated. Related to Figure 2.

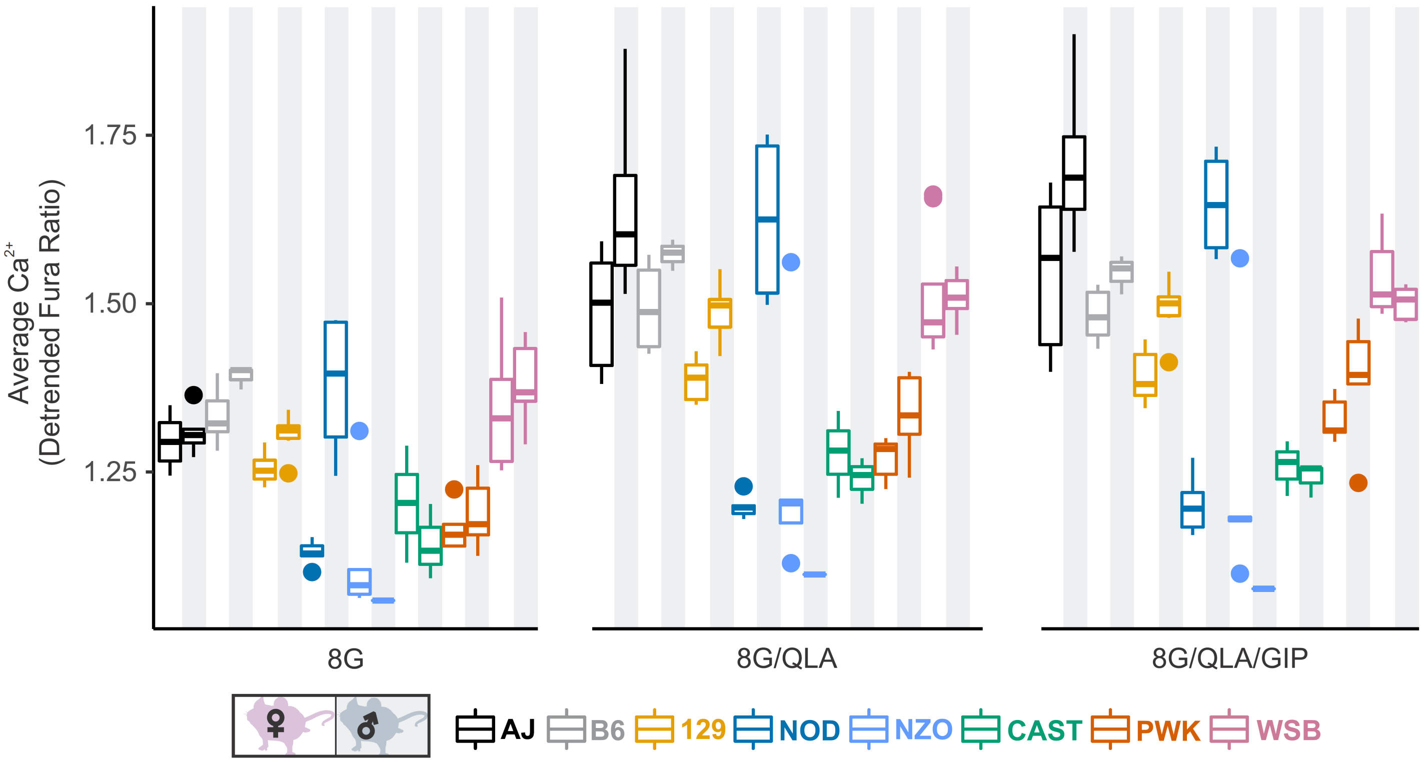


**Supplemental Figure 4. Average $Ca^{2+}$ for the stimulatory conditions.** Average $Ca^{2+}$ (detrended Fura Red ratio) for 8 mM glucose (8G, left panel), 8G with 1.25 mM L-alanine, 2 mM L-glutamine, and 0.5 mM L-leucine (8G/QLA, middle panel), and 8G/QLA with 10nM GIP (8G/QLA/GIP, right panel) are shown for each strain/sex. n = 3-8 mice per strain. Related to Figures 3 and 4.

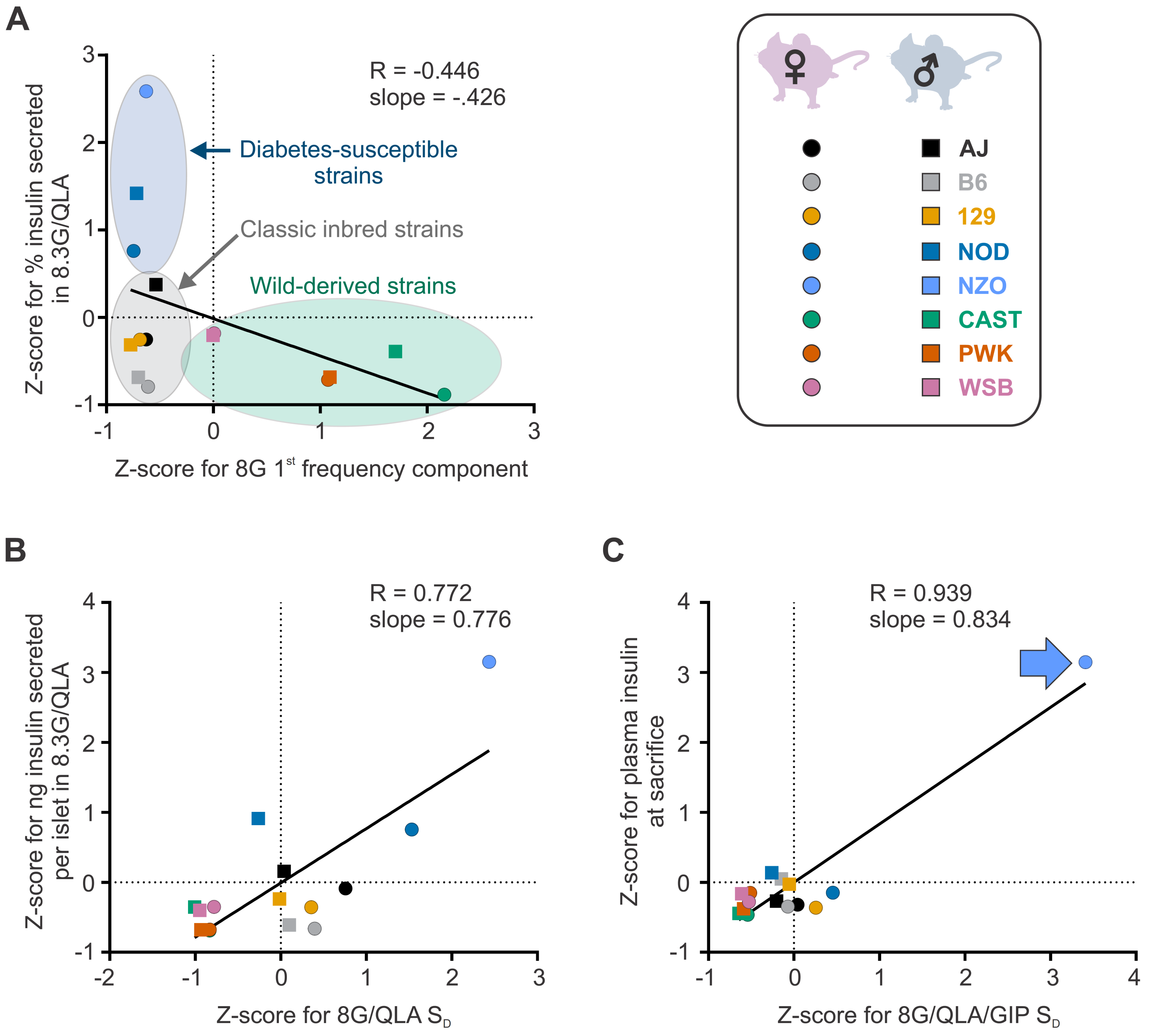


**Supplemental Figure 5. Differential strain and sex effects in correlations between traits.** The correlations between Z-scores for the calcium trait (x axis) and the other indicated trait (y axis) are shown for the eight strains (key, upper right). Males are indicated as boxes (□), females as circles (○) in each plot, where the color indicates the strain. **(A)** The correlation between the Z-scores for the 1st frequency component in 8mM glucose (8G) and the percent insulin secreted in 8.3mM glucose with 1.25 mM L-alanine, 2 mM L-glutamine, and 0.5 mM L-leucine (8.3G/QLA) reveal clustering of strains into the disease susceptible strains (blue ellipse), classic inbred strains (gray ellipse), and wild-derived strains (green ellipse). **(B)** The correlation between the Z-scores for silent duration ($S_D$) in 8G/QLA and the ng of insulin secreted per islet in 8.3G/QLA show less of a separation between the three groups. **(C)** Some correlations, such as that between Z-scores for the $S_D$ in 8mM glucose with 1.25 mM L-alanine, 2 mM L-glutamine, 0.5 mM L-leucine and 10nM GIP (8G/QLA/GIP) and the plasma insulin level at sacrifice show strong effects of single strains (e.g. NZO, indicated by blue arrow). In each plot the Pearson's R value and slope are indicated. Related to Figure 4.

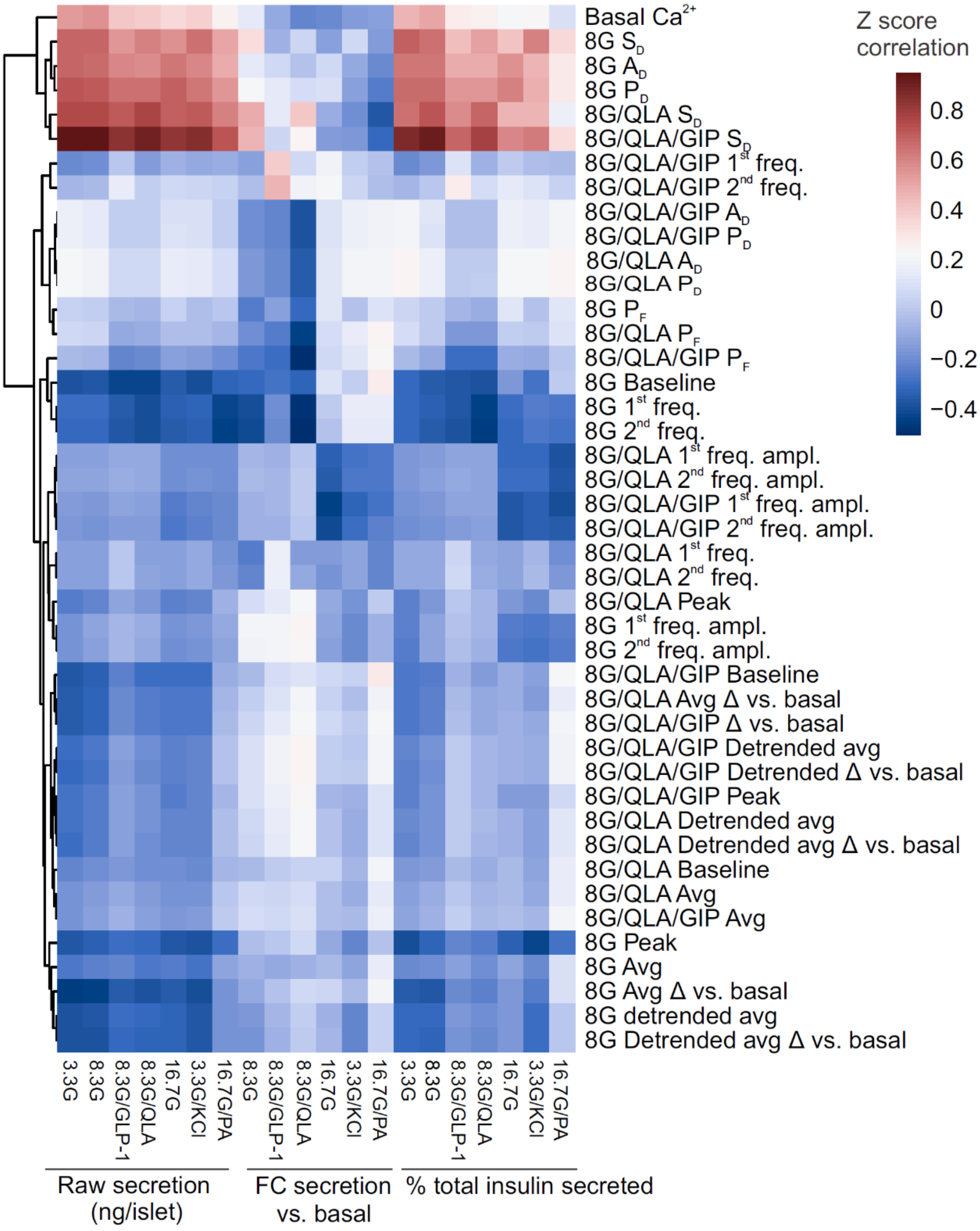


**Supplemental Figure 6. Correlation reveals specific $Ca^{2+}$ parameters highly associated with insulin secretion.** Heatmap displaying the correlation coefficients between the z-scores for $Ca^{2+}$ wave metrics and the z-scores for raw insulin secreted, fold-change (FC) over basal insulin secreted, and % of total islet insulin secreted. Insulin measurements were previously collected (17) for seven different secretagogues (16.7 mM glucose + 0.5 mM palmitic acid (16.7G/PA); 3.3 mM glucose + 50 mM KCl (3.3G/KCl); 16.7 mM glucose (16.7G); 8.3 mM glucose + 1.25 mM L-alanine, 2 mM L-glutamine, and

0.5 mM L-leucine (8.3G/QLA); 8.3 mM glucose + 100 nM GLP-1 (8.3G/GLP-1); 8.3 mM glucose (8.3G); and 3.3 mM glucose (3.3G)). Perifusion conditions included 8mM glucose (8G); 8 mM glucose + 1.25 mM L-alanine, 2 mM L-glutamine, and 0.5 mM L-leucine (8G/QLA); 8 mM glucose + QLA + 10 nM GIP (8G/QLA/GIP). Unsupervised clustering of the $Ca^{2+}$ wave parameters revealed several parameters highly correlated to multiple insulin secretion conditions. Parameters included: average $Ca^{2+}$ in 2 mM glucose (basal $Ca^{2+}$), in 8 mM glucose (8G avg.), in 8G/QLA (8G/QLA avg), and in 8G/QLA/GIP (8G/QLA/GIP avg.); average detrended $Ca^{2+}$ in 8 mM glucose (8G detr. avg.), in 8G/QLA (8G/QLA detr. avg), and in 8G/QLA/GIP (8G/QLA/GIP detr. avg.); average change in $Ca^{2+}$ vs. basal in 8 mM glucose (8G avg. Δ vs. 2G), in 8G/QLA (8G/QLA avg. Δ vs. 2G), and in 8G/QLA/GIP (8G/QLA/GIP avg. Δ vs. 2G); change in detrended average $Ca^{2+}$ vs. basal in 8 mM glucose (8G detr. Δ vs. 2G), in 8/QLA (8G/QLA detr. Δ vs. 2G), and in 8G/QLA/GIP (8G/QLA/GIP detr. Δ vs. 2G); average oscillation peak $Ca^{2+}$ in 8G (8G peak), in 8G/QLA (8G/QLA peak), and in 8G/QLA/GIP (8G/QLA/GIP peak); average oscillation baseline $Ca^{2+}$ in 8G (8G baseline), in 8G/QLA (8G/QLA baseline), and in 8G/QLA/GIP (8G/QLA/GIP baseline); pulse duration in 8G (8G $P_D$), in 8G/QLA (8G/QLA $P_D$), and in 8G/QLA/GIP (8G/QLA/GIP $P_D$); active duration in 8 mM glucose (8G $A_D$), in 8G/QLA (8G/QLA $A_D$), and in 8G/QLA/GIP (8G/QLA/GIP $A_D$); silent duration in 8 mM glucose (8G $S_D$), in 8G/QLA (8G/QLA $S_D$), and in 8G/QLA/GIP (8G/QLA/GIP $S_D$); plateau fraction in 8 mM glucose (8G $P_F$), in 8G/QLA (8G/QLA $P_F$), and in 8G/QLA/GIP (8G/QLA/GIP $P_F$); spectral density 1st component frequency in 8 mM glucose (8G 1st freq.), in 8G/QLA (8G/QLA 1st freq.), and in 8G/QLA/GIP (8G/QLA/GIP 1st freq.); spectral density 2nd component frequency in 8 mM glucose (8G 2nd freq.), in 8G/QLA (8G/QLA 2nd freq.), and in 8G/QLA/GIP (8G/QLA/GIP 2nd freq.); contribution of the 1st component to the $Ca^{2+}$ waveform for 8 mM glucose (8G 1st freq. amp.), for 8G/QLA (8G/QLA 1st freq. amp.), and for 8G/QLA/GIP (8G/QLA/GIP 1st freq. amp.); and contribution of the 2nd component to the $Ca^{2+}$ waveform for 8 mM glucose (8G 2nd freq. amp.), for 8G/QLA (8G/QLA 2nd freq. amp.), and for 8G/QLA/GIP (8G/QLA/GIP 2nd freq. amp.). Related to Figure 4.

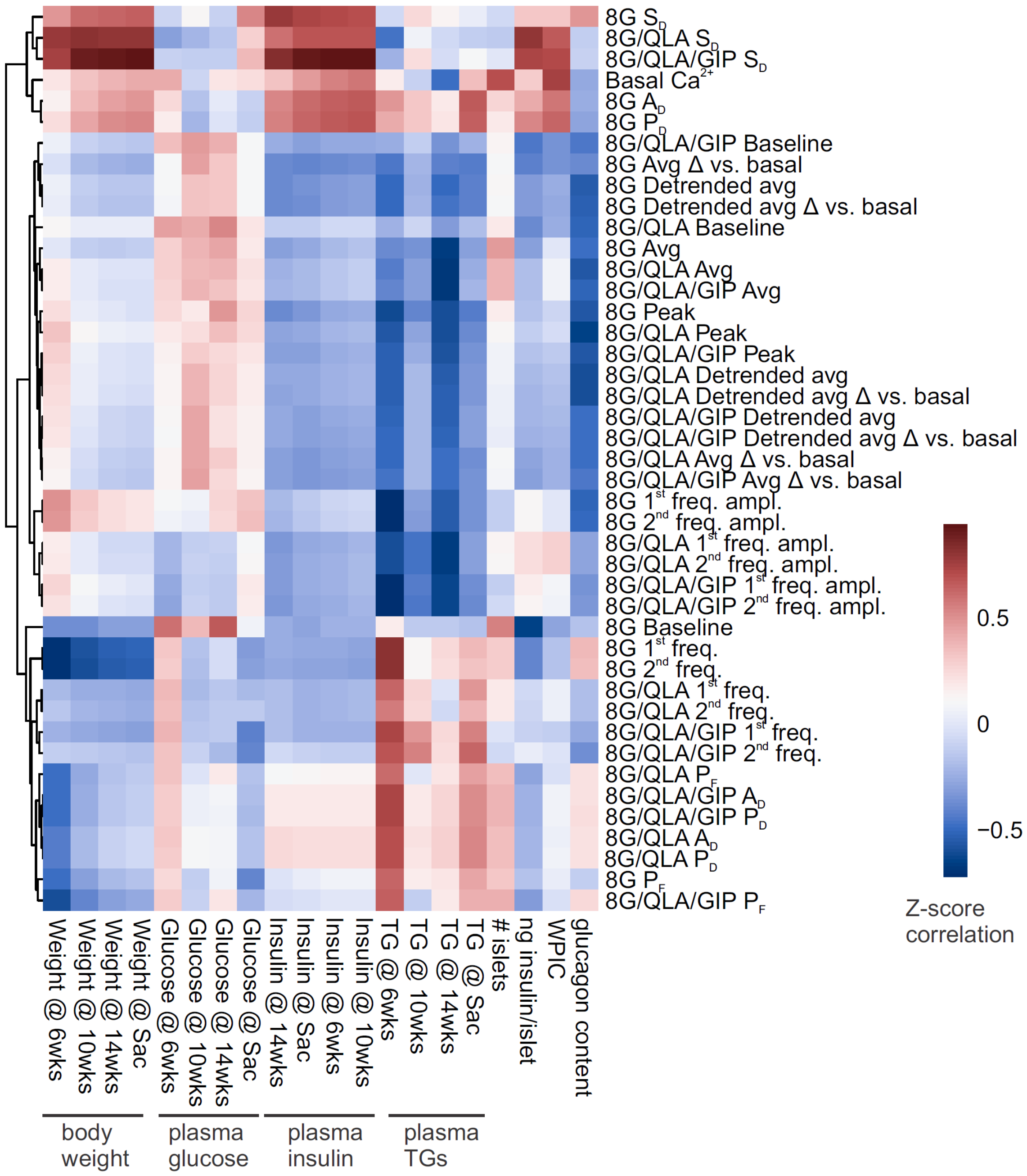


**Supplemental Figure 7. Correlation reveals specific $Ca^{2+}$ parameters highly associated with *in vivo* traits.** Heatmap displaying the correlation coefficients between the z-scores for $Ca^{2+}$ wave metrics and the z-scores for mouse *in vivo* metrics previously collected (17) for the same strains and sexes. These traits included plasma insulin, glucose, and triglycerides (TGs) (measured at 6, 10, and 14 weeks of age as well as at sacrifice) as well as whole body weights at those time points. Whole pancreas insulin content (WPIC), islet glucagon, islet number, and islet insulin content were also determined. $Ca^{2+}$ perifusion conditions included 8mM glucose (8G); 8 mM glucose + 1.25 mM L-alanine, 2 mM L-glutamine, and 0.5 mM L-leucine (8G/QLA); 8 mM glucose + QLA + 10 nM GIP (8G/QLA/GIP). Unsupervised clustering of the $Ca^{2+}$ wave parameters revealed several parameters highly correlated to multiple *in vivo* traits. Parameters included: average $Ca^{2+}$ in 2 mM glucose (basal $Ca^{2+}$), in 8 mM glucose (8G), in 8G/QLA (8G/QLA), and in 8G/QLA/GIP (8G/QLA/GIP); average detrended $Ca^{2+}$ in 8 mM glucose (8G detr.), in 8G/QLA (8G/QLA detr.), and in 8G/QLA/GIP (8G/QLA/GIP detr.); average change in $Ca^{2+}$ vs. basal in 8 mM glucose (8G Δ vs. basal), in 8G/QLA (8G/QLA Δ vs. basal), and in 8G/QLA/GIP (8G/QLA/GIP Δ vs. basal); change in detrended average $Ca^{2+}$ vs. basal in 8 mM glucose (8G detr. Δ vs. basal), in 8/QLA (8G/QLA detr. Δ vs. basal), and in 8G/QLA/GIP (8G/QLA/GIP detr. Δ vs. basal); average oscillation peak $Ca^{2+}$ in 8G (8G peak), in 8G/QLA (8G/QLA peak), and in 8G/QLA/GIP (8G/QLA/GIP peak); average oscillation baseline $Ca^{2+}$ in 8G (8G baseline), in 8G/QLA (8G/QLA baseline), and in 8G/QLA/GIP (8G/QLA/GIP baseline); pulse duration in 8G (8G $P_D$), in 8G/QLA (8G/QLA $P_D$), and in 8G/QLA/GIP (8G/QLA/GIP $P_D$); active duration in 8 mM glucose (8G $A_D$), in 8G/QLA (8G/QLA $A_D$), and in 8G/QLA/GIP (8G/QLA/GIP $A_D$); silent duration in 8 mM glucose (8G $S_D$), in 8G/QLA (8G/QLA $S_D$), and in 8G/QLA/GIP (8G/QLA/GIP $S_D$); plateau fraction in 8 mM glucose (8G $P_F$), in 8G/QLA (8G/QLA $P_F$), and in 8G/QLA/GIP (8G/QLA/GIP $P_F$); spectral density 1st component frequency in 8 mM glucose (8G 1st freq.), in 8G/QLA (8G/QLA 1st freq.), and in 8G/QLA/GIP (8G/QLA/GIP 1st freq.); spectral density 2nd component frequency in 8 mM glucose (8G 2nd freq.), in 8G/QLA (8G/QLA 2nd freq.), and in 8G/QLA/GIP (8G/QLA/GIP 2nd freq.); contribution of the 1st component to the $Ca^{2+}$ waveform for 8 mM glucose (8G 1st freq. amp.), for 8G/QLA (8G/QLA 1st freq. amp.), and for 8G/QLA/GIP (8G/QLA/GIP 1st freq. amp.); and contribution of the 2nd component to the $Ca^{2+}$ waveform for 8 mM glucose (8G 2nd freq. amp.), for 8G/QLA (8G/QLA 2nd freq. amp.), and for 8G/QLA/GIP (8G/QLA/GIP 2nd freq. amp.). Related to Figure 4.

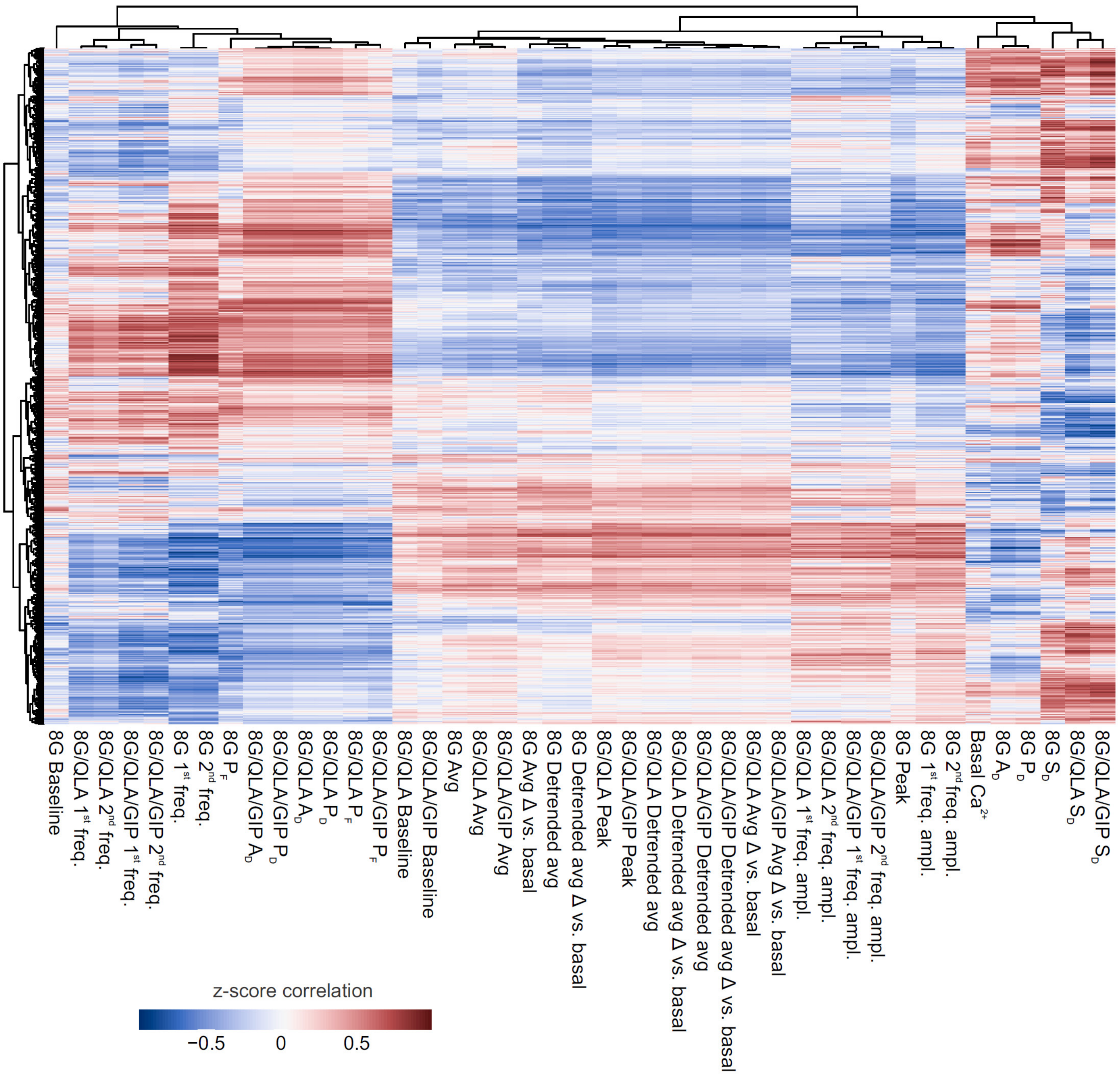


**Supplemental Figure 8. Correlation reveals proteins highly associated with specific $Ca^{2+}$ parameters.** Heatmap displaying unsupervised clustering of the correlation coefficients between the z-scores for $Ca^{2+}$ wave metrics and the z-scores for normalized islet protein abundance. Islet proteins were quantified previously (17). Perifusion conditions included 8mM glucose (8G); 8G mM glucose + 1.25 mM L-alanine, 2 mM L-glutamine, and 0.5 mM L-leucine (8G/QLA); 8 mM glucose + QLA + 10 nM

GIP (8G/QLA/GIP). Unsupervised clustering of the $Ca^{2+}$ wave parameters revealed several parameters highly correlated to multiple insulin secretion conditions. Parameters included: average $Ca^{2+}$ in 2 mM glucose (basal $Ca^{2+}$), in 8 mM glucose (8G avg.), in 8G/QLA (8G/QLA avg), and in 8G/QLA/GIP (8G/QLA/GIP avg.); average detrended $Ca^{2+}$ in 8 mM glucose (8G detr. avg.), in 8G/QLA (8G/QLA detr. avg), and in 8G/QLA/GIP (8G/QLA/GIP detr. avg.); average change in $Ca^{2+}$ vs. basal in 8 mM glucose (8G avg. Δ vs. 2G), in 8G/QLA (8G/QLA avg. Δ vs. 2G), and in 8G/QLA/GIP (8G/QLA/GIP avg. Δ vs. 2G); change in detrended average $Ca^{2+}$ vs. basal in 8 mM glucose (8G detr. Δ vs. 2G), in 8G/QLA (8G/QLA detr. Δ vs. 2G), and in 8G/QLA/GIP (8G/QLA/GIP detr. Δ vs. 2G); average oscillation peak $Ca^{2+}$ in 8G (8G peak), in 8G/QLA (8G/QLA peak), and in 8G/QLA/GIP (8G/QLA/GIP peak); average oscillation baseline $Ca^{2+}$ in 8G (8G baseline), in 8G/QLA (8G/QLA baseline), and in 8G/QLA/GIP (8G/QLA/GIP baseline); pulse duration in 8G (8G $P_D$), in 8G/QLA (8G/QLA $P_D$), and in 8G/QLA/GIP (8G/QLA/GIP $P_D$); active duration in 8G (8G $A_D$), in 8G/QLA (8G/QLA $A_D$), and in 8G/QLA/GIP (8G/QLA/GIP $A_D$); silent duration in 8 mM glucose (8G $S_D$), in 8G/QLA (8G/QLA $S_D$), and in 8G/QLA/GIP (8G/QLA/GIP $S_D$); plateau fraction in 8 mM glucose (8G $P_F$), in 8G/QLA (8G/QLA $P_F$), and in 8G/QLA/GIP (8G/QLA/GIP $P_F$); spectral density 1st component frequency in 8 mM glucose (8G 1st freq.), in 8G/QLA (8G/QLA 1st freq.), and in 8G/QLA/GIP (8G/QLA/GIP 1st freq.); spectral density 2nd component frequency in 8 mM glucose (8G 2nd freq.), in 8G/QLA (8G/QLA 2nd freq.), and in 8G/QLA/GIP (8G/QLA/GIP 2nd freq.); contribution of the 1st component to the $Ca^{2+}$ waveform for 8 mM glucose (8G 1st freq. amp.), for 8G/QLA (8G/QLA 1st freq. amp.), and for 8G/QLA/GIP (8G/QLA/GIP 1st freq. amp.); and contribution of the 2nd component to the $Ca^{2+}$ waveform for 8 mM glucose (8G 2nd freq. amp.), for 8G/QLA (8G/QLA 2nd freq. amp.), and for 8G/QLA/GIP (8G/QLA/GIP 2nd freq. amp.). Related to Figure 5.

**TABLES:**

**Supplemental Table 1: Enrichments for the highly correlated and anticorrelated proteins.** The Enrichr tool (37, 88) queries multiple databases for information regarding gene lists and queries can be stored for access later using hyperlinks. This Excel file contains 5 tabs. The "Key" tab indicates the contents of the file. The "Uniprot_IDs_correlated" and "Uniprot_IDs_anticorrelated" tabs each respectively contain in their columns lists of the Uniprot IDs for peptides correlated (coefficient > 0.5) or anticorrelated (coefficient < -0.5) to specific $Ca^{2+}$ parameters listed in the row labeled "Traits." These Uniprot IDs were queried to determine the gene names. The "Enrichr_correlated" and "Enrichr_anticorrelated" tabs each respectively contain these corresponding gene names for those proteins correlated (coefficient > 0.5) or anticorrelated (coefficient < -0.5) to specific $Ca^{2+}$ parameters listed in the row labeled "Traits." In the row "Enrichr Link" contains the Enrichr query hyperlinks for each protein list. For example, the Enrichr_correlated column B has the link https://maayanlab.cloud/Enrichr/enrich?dataset=affcd1912271cd603ec6e26304ac789e which is the Enrichr database search for the proteins listed in that column that correlate with the 1st frequency component in 8G.
